# Characterization of Pro-Fibrotic Signaling Pathways using Human Hepatic Organoids

**DOI:** 10.1101/2023.04.25.538102

**Authors:** Yuan Guan, Zhuoqing Fang, Angelina Hu, Sarah Roberts, Meiyue Wang, Wenlong Ren, Patrik K. Johansson, Sarah C. Heilshorn, Annika Enejder, Gary Peltz

**Author notes:** Address Correspondence to 300 Pasteur Dr. Room L232 Stanford, CA 94305. Author Contributions*. Y.G, S.R., A.H., P.K.J generated experimental data; Z.F., Y.G., P.K.J., A.E., M.W., W.R., S.C.H. and G.P. analyzed data; The paper was written by Y.G. and G.P with input from all authors.

## Abstract

Due to the limitations of available *in vitro* systems and animal models, we lack a detailed understanding of the pathogenetic mechanisms and have minimal treatment options for liver fibrosis. To overcome this barrier, we engineered a live cell imaging system that identifies collagen producing cells in a human multi-lineage hepatic organoid. This system was adapted for use as a microwell-based platform (i.e., microHOs) where exposure to PDGF or TGFβ1 induced the formation of thick collagen fibers. Transcriptomic analysis revealed that TGFβ1 exposure converted mesenchymal cells into myofibroblast-like cells that contribute to the development of liver fibrosis. When pro-fibrotic intracellular signaling pathways were examined using pharmacological probes, the anti-fibrotic effect of receptor-specific tyrosine kinase inhibitors was limited to the fibrosis induced by the corresponding growth factor, which indicates that their anti-fibrotic efficacy would be limited to fibrotic diseases that were solely mediated by that growth factor. Transcriptomic and transcription factor activation analyses were used to identify pathways that were jointly activated by PDGF and TGFβ1. GSK3β or p38 MAPK inhibitors could prevent TGFβ1- or PDGF-induced fibrosis in microHOs because they block intracellular signaling pathways that are commonly utilized by the TGFβ1 and PDGF receptors. Hence, these studies identified GSK3β and p38 MAPK inhibitors as potential new broad-spectrum therapies for liver fibrosis, and it is likely that other new therapies could subsequently be identified using this microHO system.

---

Liver fibrosis is a pathological condition caused by the accumulation of extracellular matrix (**ECM**) that develops in response to chronic liver injury ^1, 2^. It results from an interaction between parenchymal and nonparenchymal liver cells and involves endogenous and liver infiltrating immune cells ^3, 4, 5^. The key non-parenchymal cell is the myofibroblast, which is generated when a fibrogenic stimulus induced by liver injury causes hepatic stellate cells (**HSC**) to transdifferentiate into myofibroblasts that produce fibril-forming collagens and other ECM components ^5, 6, 7, 8, 9^. Before organoid methodology was available, *in vitro* models often did not have the spectrum of cell types that mediate fibrogenesis, which limited our ability to explore fibrogenic mechanisms. Commonly used rodent models require an injury inducing agent (carbon tetrachloride, bile duct ligation, or dietary modulation); are very time consuming and expensive to run; and any conclusions drawn from them are limited by concerns about their fidelity with the processes mediating the commonly occurring forms of human liver fibrosis ^10^. However, the pathogenesis of liver fibrosis can best be studied using a multi-lineage human hepatic organoid (**HO**) that has the cellular components and the geometry of hepatic lobules, which enables the effect that biochemical signals have on fibrosis to be quantitatively characterized. We previously developed an *in vitro* model system that differentiates human induced pluripotent stem cells (**iPSCs**) into a multi-lineage HO that has hepatocytes, cholangiocytes, HSCs, and cells of other lineages (fibroblasts, macrophages, and endothelial cells); and bile ducts and other structures found in the liver lobule are formed in these HOs ^11, 12^. HOs engineered to express a causative mutation for Autosomal Recessive Polycystic Kidney Disease (**ARPKD**) developed an extensive fibrosis ^12^ characterized by a marked increase in thick collagen fiber formation, which resembled that in liver tissue obtained from ARPKD patients. The myofibroblasts in ARPKD organoids resembled those in the livers of patients with viral-induced cirrhosis or advanced non-alcoholic steatohepatitis (**NASH**) ^12^. Here, we engineered a live cell imaging system for identification of the collagen producing cells in hepatic organoids that was adapted for use in a microwell-based platform. This system was used to characterize the signals driving liver fibrosis at the whole organoid, single cell, and molecular levels. Moreover, when the interaction between the intracellular signaling pathways activated by two pro-fibrotic factors were characterized, two potential new approaches for treating liver fibrosis were identified.

## Results

### A live cell imaging system for hepatic fibrosis

Collagen is a triple helical protein that is composed of two COL1A1 (α1) and one COL1A2 (α2) chains, and increased collagen synthesis by activated myofibroblasts is a major contributor to liver fibrosis ^2^. To characterize fibrogenic mechanisms, we developed a live cell imaging system, which enabled collagen producing cells in HOs to be identified and quantitatively analyzed. A CRISPR-assisted insertion tagging system (CRISPaint) ^13^ was used to insert a Clover expression cassette (***COL1A1*-P2A Clover**) at the COOH terminus of the endogenous *COL1A1* gene in a control iPSC line. Since the insert has a self-cleaving P2A peptide, *COL1A1* mRNA expressing cells in the HOs produced from *COL1A1*- P2A Clover iPSCs were labelled with a fluorescent intracellular protein (**Fig 1A-B**). The Clover^+^ cell population increased in HO cultures at the hepatoblast stage (day 9) (**Fig. 1C-D**). Immunostaining and flow cytometry revealed that Clover^+^ cells: were markedly increased in day 21 HOs that were treated with TGFβ1 or PDGFβ (on day 13); co-expressed PDGFRβ and COL1A1; were present throughout the organoid; and were distinct from hepatocytes or cholangiocytes (**Figs. 1E-G, S1; Tables S1-S2**). Flow cytometry results confirm that Clover fluorescence is cell based (Figs. 1D, 1G). The pro-fibrotic effect of TGFβ and PDGF was consistent with the temporal pattern of expression of the mRNAs encoding their receptors (*PDGFRA, PDGFRB, TGFBR1, TGFBR2, TGFBR3, BMPR1B*) and intracellular signaling proteins (STATs, SMADs), which were expressed at the hepatoblast stage (day 9) (i.e., prior to when microHOs are exposed to PDGF or TFGβ on day 13) and in mature HOs (**Fig. S2**). Clover protein was not expressed in iPSC or hepatoblasts, but was expressed in control, TGFβ- and PDGF-treated HOs (**Fig. 1H**).

**Figure 1.**
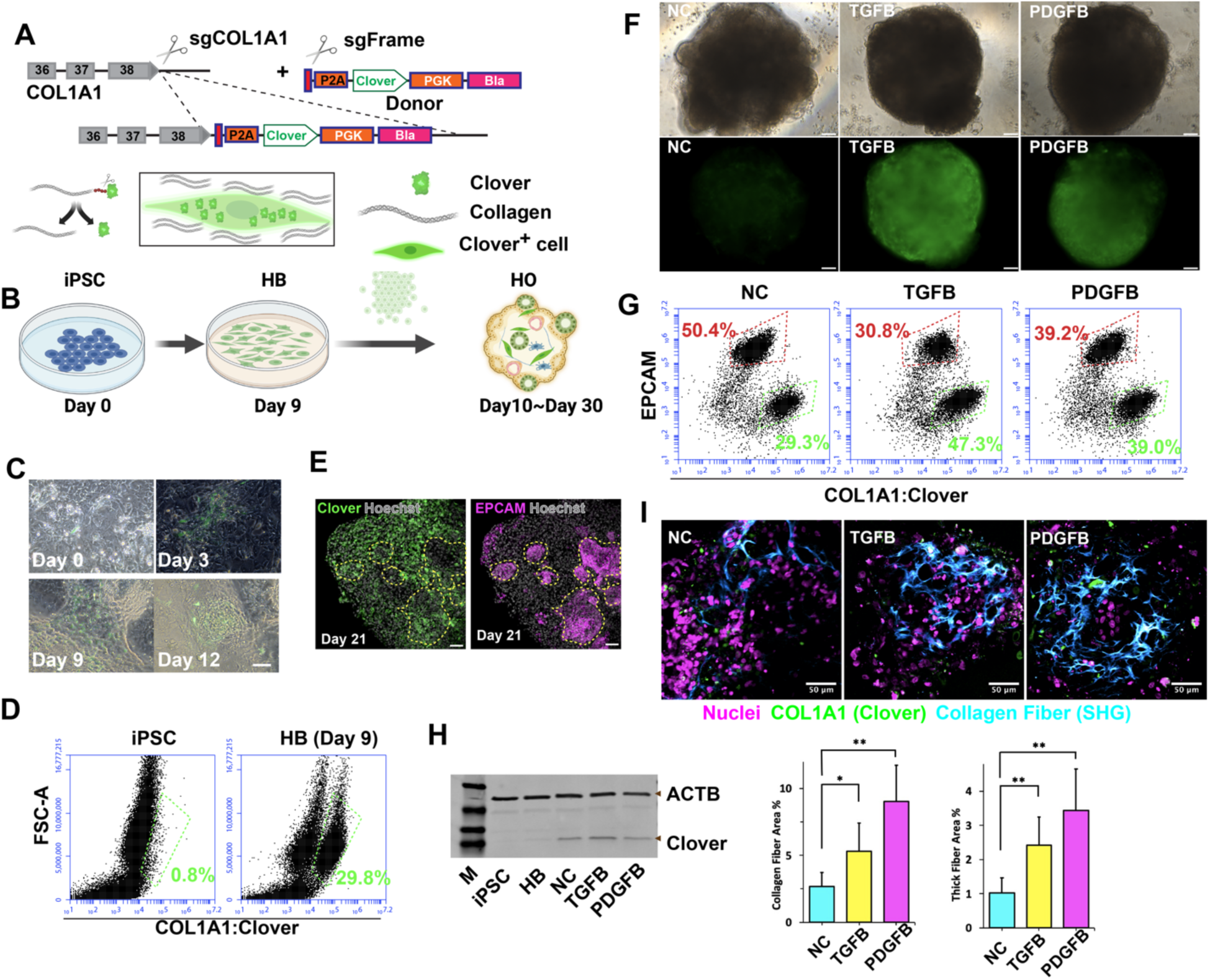
**(A)** *Top:* A diagram of the CRISPaint system used to insert P2A-Clover at the 3’ end of *COL1A1* in an iPSC line. After blasticidin selection, cells were cloned to generate the COL1A1-P2A-Clover iPSC line. *Bottom*: The self-cleaving P2A peptide enables *COL1A1* expressing cells to be labelled with a fluorescent intracellular protein (Clover). (**B**) The collagen producing cells in a HO produced from COL1A1-P2A-Clover iPSCs are labelled with an intracellular fluorescent Clover protein. (**C**) An overlay of bright field and fluorescent images of differentiating COL1A1-P2A Clover HO cultures reveals that the number of Clover^+^ cells increase between days 3 and 12. Scale bar: 50 μm. (**D**) Flow cytometry scatter plots show that the Clover^+^ cell population is absent in iPSC (day 0) but appears in (day 9) hepatoblasts in differentiating *COL1A1*-P2A Clover HO cultures. (**E**) Stacked confocal images show the location of antibody-stained EPCAM^+^ (bottom) or fluorescent Clover^+^ (top) cells in day 20 COL1A1-P2A-Clover organoids. The yellow dashed circles (bottom) indicate areas with EPCAM^+^ (i.e., hepatocytes or cholangiocytes) cells; and Clover^+^ cells are not found within those areas but are present in other areas of the HO. Scale bar: 50 μm. (**F**) Bright field (top) and fluorescence (bottom) images of day 21 *COL1A1*-P2A Clover HOs treated on day 13 with no addition (NC), 50 ng/ml TGFβ1 or 50 ng/ml PDGFβ. Both growth factors induced a marked increase in COL1A1^+^ cells. Scale bars, 50 μm. (**G**) Flow cytometry scatter plots show that Clover^+^ cells are increased in mature day 21 *COL1A1*-P2A Clover HOs after TGFβ or PDGF exposure. Also, the Clover^+^ cells are distinct from the EPCAM^+^ (hepatocytes or cholangiocytes) cells in the HOs. (**H**) Immunoblot showing Clover protein expression in differentiating HO cultures (iPSC, day 9 hepatoblasts (HB)); and in day 21 control (NC), PDGF- or TGFβ-treated HOs. The positions of Clover and β-actin (ACTB) proteins are indicated. (**I**) SHG analysis of the collagen fibers formed in human HOs. *Top:* Depth color-coded projections of collagen fibers within day 21 control organoids, or day 21 organoids that were treated with 50 ng/ml TGFβ1 or 50 ng/ml PDGF on day 13. Control organoids (left) have isolated regions with relatively thin collagen fibers (shown in blue). In contrast, the TGFβ1 or PDGF-treated organoids form a network of thick collagen fibers that extend throughout the entire organoid. Collagen producing cells (green) can also be seen in these images. Scale bars, 50 μ. *Bottom:* A quantitative comparison of collagen fiber area in SHG images was performed for control, TGFβ1- or PDGF-treated hepatic organoids (n = 5 per measurement) on day 21. There is a statistically significant increase in total collagen abundance (collagen fiber area=total area with collagen fibers/area of the imaged organoids) in the organoids after exposure to TGFβ1 (*, p < 0.05) or PDGF (**, p <0.01, Welch t-test). There was also a statistically significant increase in the abundance of thick collagen fibers (i.e., those fibers > 3 um diameter) in the TGFβ1 and PDGF-treated hepatic organoids.

The spatial distribution and morphology of the collagen fibers in HOs were examined using Second Harmonic Generation (**SHG**) microscopy ^14, 15^, which has previously been used to quantitatively analyze liver fibrosis in human patients ^16, 17^ and to monitor fibrosis in ARPKD HOs^12^. Of importance, SHG measures thick collagen fiber formation; which is a complex biochemical pathway that requires the coordinated activity of multiple liver enzymes that include proteases and other post-translational processing enzymes that act on collagen, and subsequently the activity of lysyl hydroxylases and lysyl oxidases to generate thick collagen fibers. ^18^. SHG images revealed that control HOs had sporadic networks of cross-linked collagen fibers surrounding the cells in some isolated regions, while the TGFβ1 or PDGF-treated organoids had more extensive networks of thick collagen fibers throughout the entire HO. A quantitative comparison indicated that TGFβ1 (p < 0.05) or PDGF (p <0.01) exposure induced statistically significant increases in total collagen abundance and in the abundance of thick collagen fibers (i.e., > 3 um diameter) formed in HOs (p<0.01) (**Fig. 1I**). The SHG and immunostaining results indicate that PDGF or TGFβ1 increases the numbers of collagen synthesizing cells and the amount of thick collagen fibers formed in HOs.

### A high content screening system (**HCS**) for serially monitoring the effect of biochemical signals on liver fibrosis

Culture conditions were modified to enable HOs to reproducibly grow to a standard size in microwells, which we refer to as ‘*microHOs*’ (**Fig. 2A-B**). (**Fig. 2A-B**). Trichrome staining confirmed that TGFβ or PDGF induced a large increase in collagen in microHOs (**Fig. 2C**). The TGFβ effect on thick collagen fiber formation was confirmed by 4-hydroxyproline (4OH-Pro) measurements: 4OH-Pro was absent in iPSC; it increased in day 9 hepatoblasts and in day 21 control HOs; but was significantly increased after PDGF or TGFβ exposure (p<0.001 vs control) (**Fig. 2D**). HCS was performed using a confocal fluorescence microplate imager with an automated program that we developed for analyzing multiple Z-stacked confocal images. TGFβ1 addition on day 13 induced a sustained increase in COL1A1^+^ cells in microHOs that was blocked by co-administration of TGFRβ1 tyrosine kinase inhibitors (10 uM SB431542 ^19, 20^ or A83-01 ^21^) (**Fig. 2E-F**). PDGFβ addition on day 13 also increased the number of COL1A1^+^ cells in microHOs, which was blocked by addition of a PDGFRβ tyrosine kinase inhibitor (10 uM Imatinib ^22, 23^) (**Fig. 3, top**). The microHO system was also used to examine the effect of five cytokines (IL-4 ^24^, IL-6 ^25^, IL-13, IL-33 ^26, 27^, TNFα ^28^) and three growth factors (IGF1 ^29^, VEGF ^30^, CCL3 ^31^) that had been directly or indirectly implicated in the pathogenesis of NASH or liver fibrosis. None of these agents induced fibrosis in microHOs (**Fig. S3**). Analysis of the temporal pattern of receptor expression in differentiating HO cultures using previously obtained transcriptomic data ^12^ revealed that the mRNAs encoding the IGF-1 (*IGF1R*), IL13 (*IL13RA1*), and VEGF (*VEGFR-1* (*FLT1*); *VEGFR-2 (KDR*)) receptors were expressed in day 9 hepatoblasts and in mature (day 21) organoids. In contrast, the mRNAs encoding the receptors for IL33 (*IL1RL1 or ST2*), IL4 (*IL4R*), or for the common gamma chain that is used for intra-cellular signaling by multiple cytokine receptors (*IL2RG*) were not expressed in day 9 hepatoblasts (**Fig. S4A**). Of note, the mRNAs encoding most of these receptors are abundantly expressed in mature HOs (**Fig. S4B**). Hence, the lack of a profibrotic effect of IGF-1 and VEGF is not due to the absence of receptor expression on day 9, which is prior to when microHOs were exposed to these agents. However, the lack of a profibrotic effect of IL-4 and IL-13 could result from the absence of functional receptors on day 9, which is just before the time of exposure to those cytokines.

**Figure 2.**
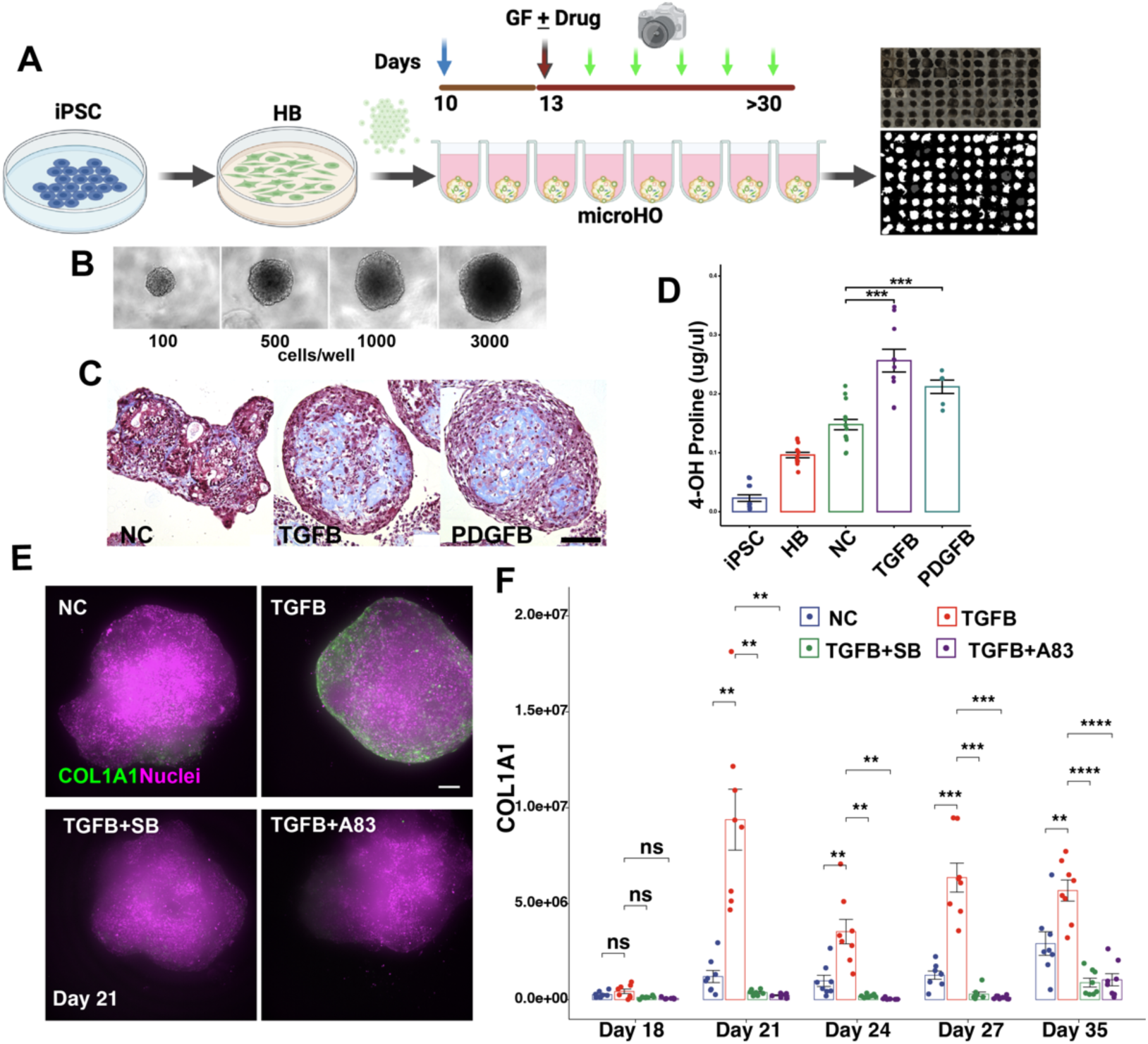
(**A**) A diagram of the HCS platform used for analysis of microHOs. COL1A1-P2A iPSCs are differentiated into hepatoblasts (HB), and 10,000 HBs are placed in each microwell on day 10. Growth factors (GF) and/or drugs are added on day 13, and the cultures are differentiated into hepatic organoids. Culture fluorescence is serially assessed to measure the number of COL1A1-producing cells in the microHO. Bright field (top) and fluorescence (bottom) images of a microwell plate with microHOs is shown on the right. (**B**) A cell number titration experiment was performed by adding the indicated numbers of HBs to microwells and the cells were differentiated through day 12. Bright field images on day 12 reveal that microHOs have a uniform shape and their size is dependent upon the number of input cells. (**C**) Trichrome staining reveals a marked increase in collagen (blue areas) in PDGF or TGFβ-treated microHOs relative to control (NC) microHOs. Scale bar, 50 um. (**D**) The 4-Hydroxy-proline (4OH-Proline) concentration was measured in differentiating HO cultures (iPSC, day 9 HB); and in day 21 control (NC), PDGF- or TGFβ-treated HOs. The bar graph shows the mean ± SE of measurements performed on a total of 15 HOs, which were generated in two separate experiments. There was a significant increase in 4-OH-Proline in the PDGF- or TGFβ-treated day 21 HOs (***, p <0.001 vs NC, t-test). (**E**) Maximum intensity projection (MIP) images of z stack sections obtained from the indicated type of microHOs on day 21. Scale bar 100 um. Clover expression is green, and nuclei stained with Hoechst 33342 are purple. (**F**) TGFβ1- induced fibrosis in microHOs is blocked by TGFRβ1 inhibitors. COL1A1-P2A Clover HOs were formed by adding 10,000 HBs to each microwell. Then, either nothing (NC), or 50 ng/ml TGFβ1 ± 10 uM TGFRβ1 inhibitors (SB431542 (SB) or A83-01 (A83)) was added to each microwell on day 13. The amount of COL1A1^+^ cells within a microHO were serially measured on days 18 through 35. Each dot represents a measurement made on one microHO, and 8 microHOs per treatment were assessed per condition. The significance indicators are ns, not significant; *, p<0.05; **, p<0.01; ***, p<0.001 or ****, p<0.0001 for measurements in TGFβ1 treated vs NC microHOs; and for microHOs that were treated with TGFβ1 and drug, the p-values are calculated relative to the TGFβ1 treated microHOs. A two-way ANOVA indicates that drug treatment and time are two variables that have a significant interaction on the fluorescence measurements (p=1.66 x 10^-15^) (**Table S9**).

**Figure 3.**
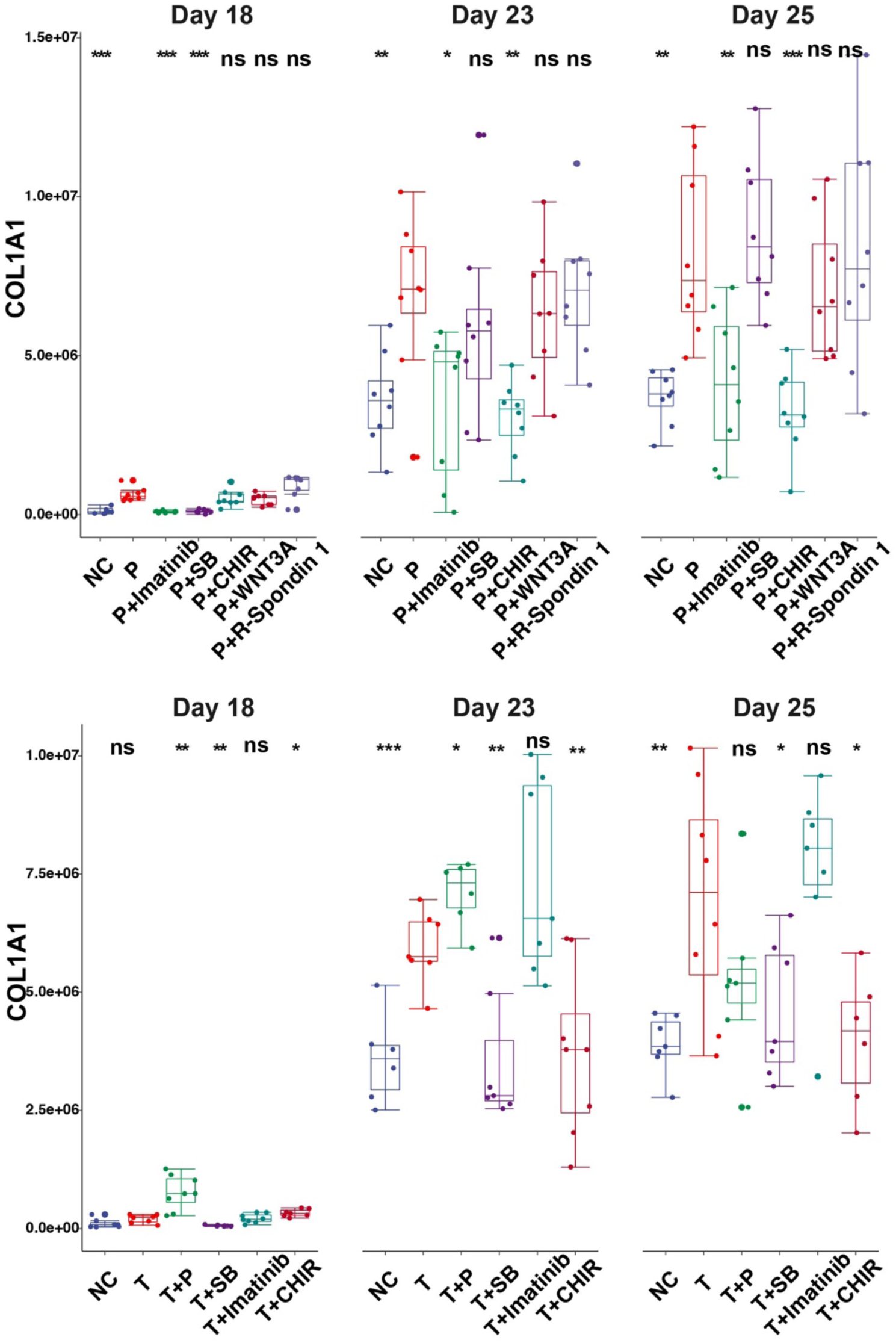
*Top:* The PDGFβ-induced increase in COL1A1^+^ cells is blocked by PDGFRβ and GSK3β inhibitors. *COL1A1*-P2A Clover HOs were formed by adding 10,000 cells from day 9 organoid cultures to each microwell. Then, either nothing (NC), 50 ng/ml PDGFβ (P) or 50 ng/ml PDGFβ with 10 μM imatinib, 10 uM SB431542 (SB), 10 uM CHIR99021 (CHIR), 100 ng/ml Wnt3a or 100 ng/ml R-spondin 1 was added to each microHO on day 13. *Bottom:* The TGFβ1 induced increase in COL1A1^+^ cells was blocked by TGFRβ and GSK3β inhibitors. Either nothing (NC), 50 ng/ml TGFβ1 (T), 50 ng/ml TGFβ1 with 50 ng/ml PDGF (T+P), or 50 ng/ml TGFβ1 and either 10 μM imatinib, 10 uM SB431542 (SB), or 10 uM CHIR99021 (CHIR) was added to each microHO on day 13. microHO fluorescence, which indicates the amount of COL1A1^+^ cells, was serially measured on days 18 through 25. Each dot represents a measurement made on a microHO, the thick line is the median for 8 microHOs that were assessed, and the box plot shows the 25 to 75% range for the measurements. In the panels: ns, not significant; *, p<0.05; or **, p<0.01, ***, p<0.001 for the measurement relative to the value in the NC; or when growth factors or inhibitors were added, the p-values were calculated relative to that of the PDGFβ or TGFβ1 treated cultures. A two-way ANOVA indicates that drug treatment and time are two variables that have a significant interaction on the fluorescence measurements (Table S9).

Since hedgehog signaling pathway activation increases *COL1A1* mRNA expression in myofibroblasts ^32, 33^ and promotes myofibroblast accumulation and liver fibrosis ^34^, we examined whether a hedgehog pathway activator would induce fibrosis in microHOs. Smoothened (**SMO**) is a seven transmembrane protein ^35, 36^ that accumulates in the primary cilium, and transduces a Hedgehog pathway signal intracellularly. However, addition of a SMO agonist (SAG), which activates Hedgehog signaling ^37^, did not induce fibrosis in microHO cultures (Fig. S3).

We next examined whether addition of seven free fatty acids (at 0 to 400 uM concentrations), which included their saturated and corresponding cis-9 (or trans-9) monounsaturated forms, on day 13 to microHO cultures had a pro-fibrotic effect. Previously obtained transcriptomic data from developing and mature HO cultures revealed that the mRNAs encoding proteins required for free fatty acid uptake (i.e., *SLC27A1, SCC27A5, FABP5, and Acyl-CoA Binding protein* (*ACBP*)) are expressed in day 9 HO cultures (**Fig. S5**). Even when the free fatty acids were added at concentrations below those previously reported to induce fibrosis in other types of HOs ^38^, the cells within microHOs abundantly accumulated fatty acids from the medium and there was evidence of lipotoxicity (**Fig. S6**). Somewhat unexpectedly based upon the results presented in ^38^, free fatty acid addition did not have a pro-fibrotic effect in microHOs (**Fig. S7**). The disparate findings may result from their using organoid stiffness ^38^ (rather than an increase in collagen thick fibers) as the criteria for fibrosis. Our results, which are consistent with recent findings by others ^39^, indicate that free fatty acid addition causes steatosis and lipotoxicity, but does not cause the fibrosis that develops in the advanced clinical stages of NASH. Altered differentiation conditions, adding other factors, or co-culture with other cell types may be required to enable HOs to model NASH-associated fibrosis.

#### Single cell Transcriptomic analysis

To characterize the pro-fibrotic effects of PDGFβ and TGFβ1, scRNA-seq data was generated from day 21 control (22,211 cells), TGFβ1-treated (17,554 cells) or PDGFβ-treated (18,013 cells) microHOs. Eleven cell clusters were identified by analysis of this scRNA-seq dataset (**Fig. 4A, Table S3**). The differentially expressed genes and the gene ontology biological process annotations (GO) in each cluster were analyzed to identify the cell type represented by each cluster (**Figs. 4B, S8** and **Table S4**). Based upon expression of canonical mRNAs and by the level of concordance between the transcriptomes of the cell clusters with the cell types in human liver, the microHOs have cholangiocyte (Cho1-3), hepatocyte (Hep), mesenchymal cell (Mes1-4) and myofibroblast (MyoF_T1-2, MyoF_P) clusters. Also, *COL1A1* and *Clover* mRNAs have an identical pattern of expression in the microHOs: both mRNAs are expressed in myofibroblasts and mesenchymal cells. In contrast, an epithelial mRNA (*EPCAM)* that is a hepatoblast and cholangiocyte marker was not expressed in the *Clover* mRNA^+^ cells (**Fig. 4C-D).** Many mRNAs found in cholangiocytes (*KRT19*, *KRT7* and *CFTR)* or in hepatocytes and cholangiocytes (*HNF4A, AFP, CDH1, ALDHA1, APOB, FGB*) were also not expressed in the *Clover* mRNA^+^ cells in the microHOs (**Fig. S9**). Myofibroblast marker mRNA (*ACTA2, PDGFRA, FAP*) expression was very low in control microHOs, but were predominantly expressed in the myofibroblast (MyoF_T1/2, MyoF_P) clusters in PDGF- or TGFβ-treated microHOs (**Fig. S10**). Consistent with the profile of *FAP* mRNA expression in microHOs (Fig. S10), FAP is absent in most tissues under basal conditions but is highly expressed on activated myofibroblasts ^40, 41^. FAP is a well-established marker for activated fibroblasts ^42^ and its serine protease activity promotes liver fibrosis by activating HSC and promoting macrophage infiltration after liver injury ^43^.

**Figure 4.**
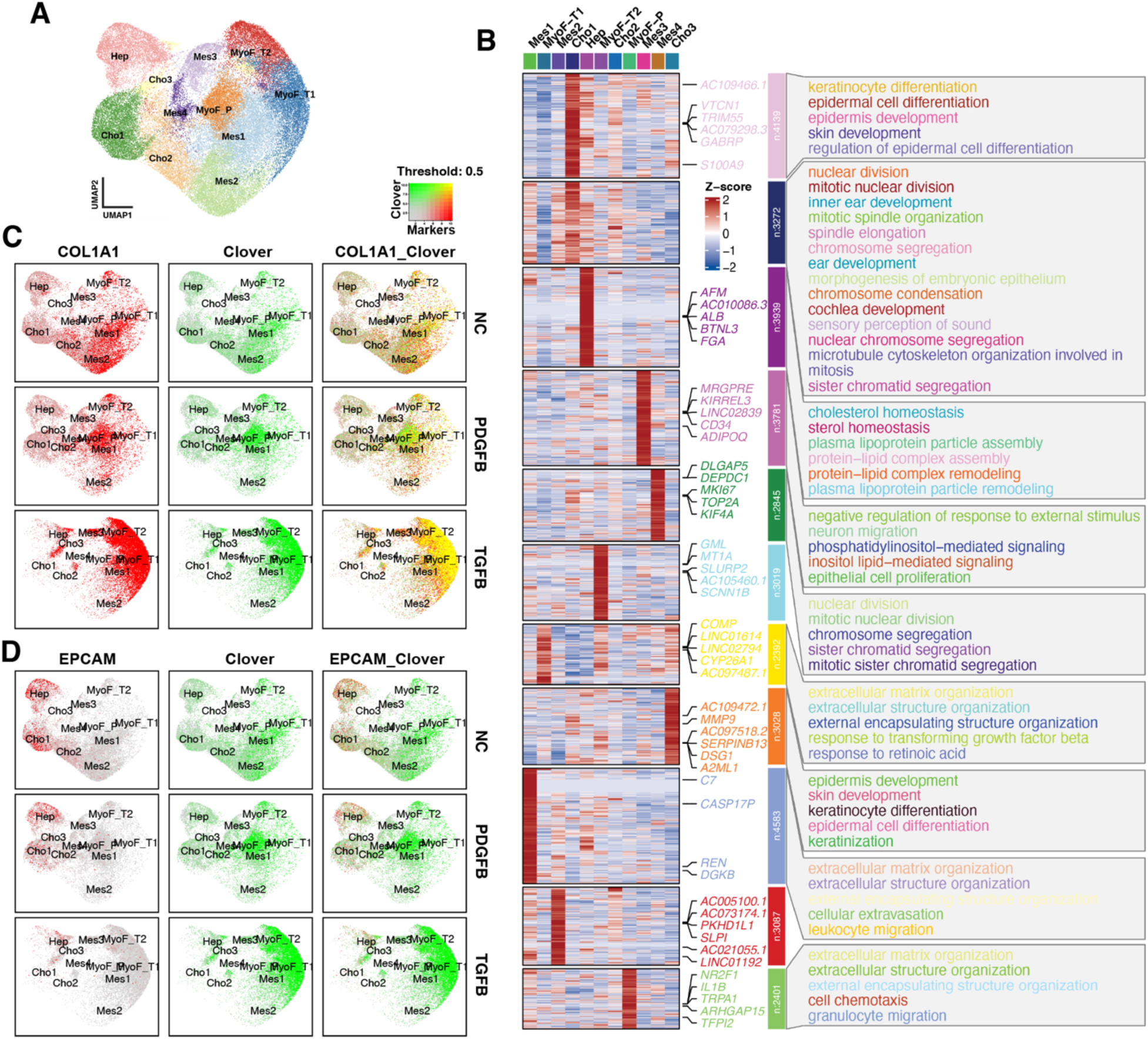
(**A**) A combined sample UMAP plot shows the 11 cell clusters identified in day 21 control (NC), PDGFβ- (P) and TGFβ1 (T)-treated microHO cultures. (**B**) Heatmaps show the DEGs and annotated genome ontology pathways identified for each cluster. (**C, D**) These feature plots show the level of expression of *COL1A1* and *Clover* mRNAs (D), or *EPCAM* and *Clover* mRNAs (E) in the UMAP plot shown in (A). As shown in the color threshold diagram, dot colors represent level of mRNA expression. *COL1A1 and Clover* mRNAs have an overlapping expression pattern; they are predominantly expressed in myofibroblasts, and in mesenchymal cells. In contrast, *EPCAM* mRNA is expressed in the hepatocyte and cholangiocyte clusters; and its expression does not overlap with that of *Clover* mRNA.

### Changes in mesenchymal and epithelial cell clusters in fibrotic microHOs

To systematically examine the changes in the cell types present in PDGFB- or TGFβ1-treated microHOs (vs NC), we ANOVA and ‘sccomp’ (a statistical model using a constrained Beta-binomial distribution ^44^) were used to statistically analyze the changes in cell type proportions. The cells in PDGF- or TGFβ1-treated microHOs were reproducibly different from those in control microHOs (**Fig. 5A-B, Fig. S11**). One myofibroblast cluster (MyoF_T1) was far more abundant in TGFβ1-treated microHOs than in control or PDGF-treated microHOs (37.5% of total vs <2%), while another myofibroblast cluster (MyoF_P) was more abundant in PDGF-treated microHOs than in control or TGFβ1-treated microHOs (20% of total vs <2%). The abundance of a mesenchymal cluster (Mes2) was markedly decreased in TGFβ or PDGF-treated microHOs (25% vs <5%) (**Fig 5C, Table S3**). A one-way ANOVA comparing the different types of cells in each of 5-independent biological replicates analyzed by scRNA-Seq indicated that MyoF_T1 (p<0.015) and MyoF_P (p<0.05) abundances were significantly increased, while Mes2 abundance was decreased (p=0.004) in TGFβ- or PDGF-treated microHOs. The ‘sccomp’ analysis indicated that the Cho1- 2 clusters were significantly decreased, while Cho3 was significantly increased in TGFβ-treated microHOs (**Table S5, Fig. S12**).

**Figure 5.**
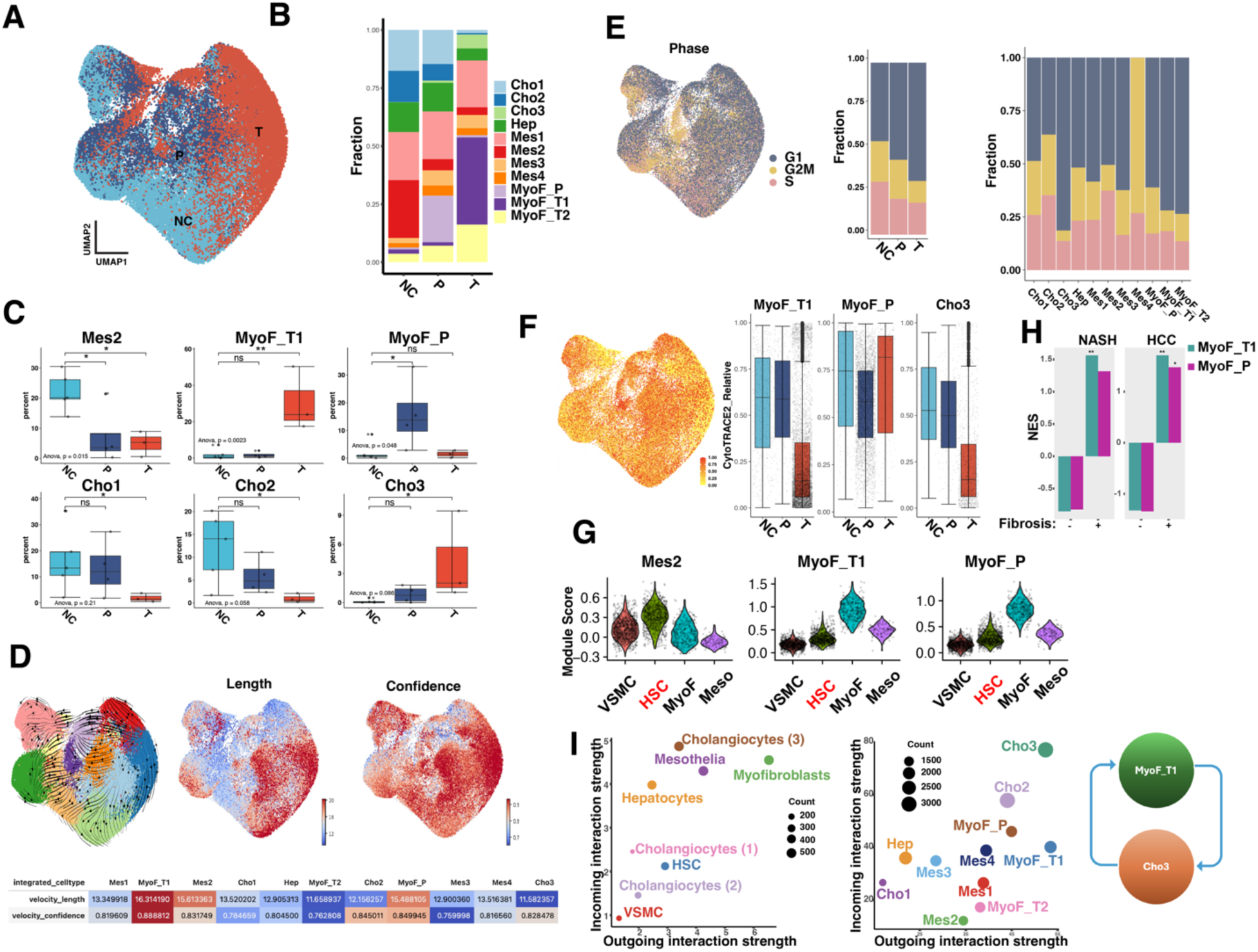
(**A**) A combined sample UMAP plot for the scRNA-seq data obtained from the cells in day 21 control (NC), PDGFβ- (P) or TGFβ1 (T)-treated microHO cultures. (**B**) The cell percentages for each of the 11 clusters in the different types of microHOs. (**C**) Boxplots show the changes in the percent of each indicated cell cluster in NC, PDGFB and TGFB treated microHOs. ns, not significant; *, p<0.05; or **, p<0.01. (**D**) RNA velocity results plotted on a UMAP plot. *Left*: stream embedding; *Middle*: velocity length indicates the rate of differentiation; *Right*: confidence, which is determined by the correlation of the velocity determined for neighboring clusters in the UMAP. *Bottom*: heatmap showing the velocity summary for all of the clusters. (**E**) Cell cycle phase is shown on the UMAP (Left); and the percentage of cells in each phase of the cell cycle is shown for NC, PDGFB- and TGFβ-treated microHOs (Middle) or for all 11 clusters. (**F**) The Cytotrace score is plotted on the UMAP or as boxplots generated for the MyoF_T1, MyoF_P and Cho3 cells in NC, PDGFB- and TGFβ-treated microHOs. (**G**) Among the cell types in human liver, the MyoF_T1 and MyoF_P transcriptomes are most similar to myofibroblasts, while Mes2 is most similar to HSC. These violin plots show the results when the Mes2, MyoF_T1, and MyoF_P gene signatures were compared with the transcriptomes of the following cell types in normal and cirrhotic (GSE136103 ^91^) human liver: VSMC, vascular smooth muscle; HSC, MyoF; and Meso, mesothelium. Each dot shows the module score of individual cells obtained for each of the four cell types. The MyoF_T1 and MyoF_P module scores are most similar to MyoF, and Mes2 has the most similarity with HSC in human liver. The MyoF module scores for MyoF_T1 and MyoF_P are 3-fold higher (p<1×10-^10^) than that of HSC, and the HSC gene signature module score for Mes2 is 4.8-fold (p<1×10-^10^) higher than for MyoF in human liver. (**H**) The bar graphs show the normalized enrichment score (NES) of the GSEA examining the association of the MyoF_T1 and MyoF_P gene signatures with liver tissue obtained from non-fibrotic (stage 1) or fibrotic (stage 4) NASH liver (GSE135251), and with resected hepatocarcinoma (HCC) tissue that was classified as non-fibrotic or fibrotic (GSE6764). The false discovery rate (FDR) p-value is indicated by the symbol above each bar: *, P < 0.05; **, P<0.01. The MyoF_T signature was strongly associated with fibrotic NASH liver; and both signatures were strongly associated with fibrotic HCC tissue but not with non-fibrotic NASH or HCC tissue. (**I**) *Left*: Cellchat scatter plots show the cell types that are the dominant senders (sources) and receivers (targets) of information that regulate cellular interactions. The axes represent the total outgoing or incoming information associated with each cell group as determined by scRNA-seq analysis. The dot size is proportional to the number of inferred links (both outgoing and incoming) for each cell group; the dot colors indicate the different cell groups. *Right*: Illustration for the hypothesis emerging from the Cellchat analysis: that communication between MyoF_T1 and cholangiocytes is key for fibrogenesis. The key pathways examined by Cellchat include cytokines and chemokine pathways that promote cell growth and repair; extracellular matrix proteins that provide structural support and influence cell behavior; and adhesion molecules that direct cell-cell contact.

To better understand the TGFβ1 on the mesenchymal and epithelial cell clusters, their differentiation state and other properties were characterized using three different computational methods that analyze scRNA-Seq data. (i) RNA velocity ^45^, which assessed the differentiation state of the cell clusters, indicated that MyoF_T1 was the most highly differentiated cluster (**Fig. 5D**). (ii) Cell cycle analysis indicated that TGFβ-treated microHOs had the highest percentage of cells in the G1 phase, which indicates that the cells were less proliferative. At the cluster level, MyoF-T1 and Cho3 had the highest percentage of cells in G1 and the lowest percentage of G2/M and S phase cells, which indicates that those clusters were less proliferative (**Fig. 5E**). (iii) Consistent with the RNA velocity results, Cytotrace2 ^46^ results indicated that MyoF_T1 and MyoF_P were the most differentiated of the mesenchymal cell clusters, whereas Cho3 was the most differentiated epithelial cluster (**Fig. 5F**). These results indicate that MyoF-T1 cells are the most differentiated of the mesenchymal cell populations in fibrotic microHOs, and they have a low level of proliferation after they exit the cell cycle exit (Fig. 5E).

Transcriptomic comparisons with various cell types in control and cirrhotic human liver were performed to further characterize the three clusters whose abundance was significantly altered by TGFβ or PDGF exposure. The results revealed that MyoF_T1 and MyoF_P clusters were most like that of myofibroblasts, whereas Mes2 was most similar to HSC in human liver (**Fig. 5G, Fig. S13, Tables S6-S7)**. As described in supplemental note 1, Gene Set Enrichment Analysis (**GSEA**) ^47^ revealed that the gene expression signature of MyoF_T1 in microHOs resembled those found in fibrotic human liver tissue caused by NASH or hepatocellular carcinoma (**Figs. 5H and S14**, **Table S8**). While the MyoF_P gene signature was positively associated with NASH or HCC-induced liver fibrotic tissue, those GSEA associations did not achieve statistical significance. Also, the pattern of expression of 9 mesenchyme and myofibroblast-specific mRNAs in mesenchymal cells and myofibroblasts in human liver was retained in Mes2 and MyoF_T1 cells in microHOs (**Fig. S15**). Taken together, the transcriptomic results indicate that TGFβ1 or PDGF exposure converts a population of HSC-like cells into myofibroblast-like cells in microHOs. Since nonparenchymal cells (i.e., the mesenchymal cells in microHOs) and epithelial cell interactions play a key role in liver fibrosis, ‘Cellchat’ ^48^ was used to examine cell cluster interactions in fibrotic microHOs and in cirrhotic human liver. It was noteworthy that the myofibroblasts in cirrhotic liver (or MyoF-T1 in microHOs) and cholangiocytes in cirrhotic liver (or Cho3 in microHOs) had the strongest incoming and outgoing interactions (**Fig. 5I**), which indicates that these cells and pathways may be targeted for pharmacologic intervention.

### Identification of druggable targets by analysis of transcription factor (TF) activation in microHOs and in cirrhotic human liver tissue

TF activity in fibrotic microHOs and in NASH-induced cirrhotic human liver tissue was examined by analysis of scRNA-seq data using the ‘decoupleR’ method ^49^. Multiple TFs were activated in fibrotic microHOs and in the cirrhotic liver tissue. Among the clusters in microHOs, MyoF_T1 and Cho3 cells exhibited the widest spectrum of TF activation (**Fig. 6A**). Of interest, SMAD4, STAT1 and JUN activity was increased in MyoF_T1 and Cho3 cells after TGFβ treatment (versus control, **Figs. 6B-C**). These TFs were also activated cirrhotic NASH liver tissue, and liver myofibroblasts displayed the highest level of activation of these TFs (versus vascular smooth muscle or HSC, **Figs. 6D-E)**. PDGF and TGFβ1 are well established drivers of liver fibrosis ^50^. TGFRB activation leads to activation of SMAD signaling pathways ^51, 52^ and PDGFR cross-linking activates the STAT3 pathway ^53^ ^54^, and both of these pathways have well-known roles in promoting hepatic fibrosis ^54^. However, it is possible that other pathways, which are jointly activated by these profibrotic agents, could provide new targets for anti-fibrotic drugs. Both TGFRB ^51, 52, 55^ and PDGFR ^56, 57^ activate the p38 Mitogen Activated Protein Kinase (MAPK) and PI3K–Akt–mTOR intracellular signaling pathways, which are also pro-fibrotic (**Fig. 6F**). Of relevance, several TFs in the p38 pathway were activated in MyoF-T1 in fibrotic microHOs (JUN, AP1, FOS) and in the myofibroblasts in cirrhotic human liver (JUNB, FOSB) (Figs. 6A, 6D). We propose that effective anti-fibrotic agents could be produced by targeting pathways (i.e., p38 or PI3K–Akt–mTOR) that are jointly activated by TGFRBs and PDGFRs.

**Figure 6.**
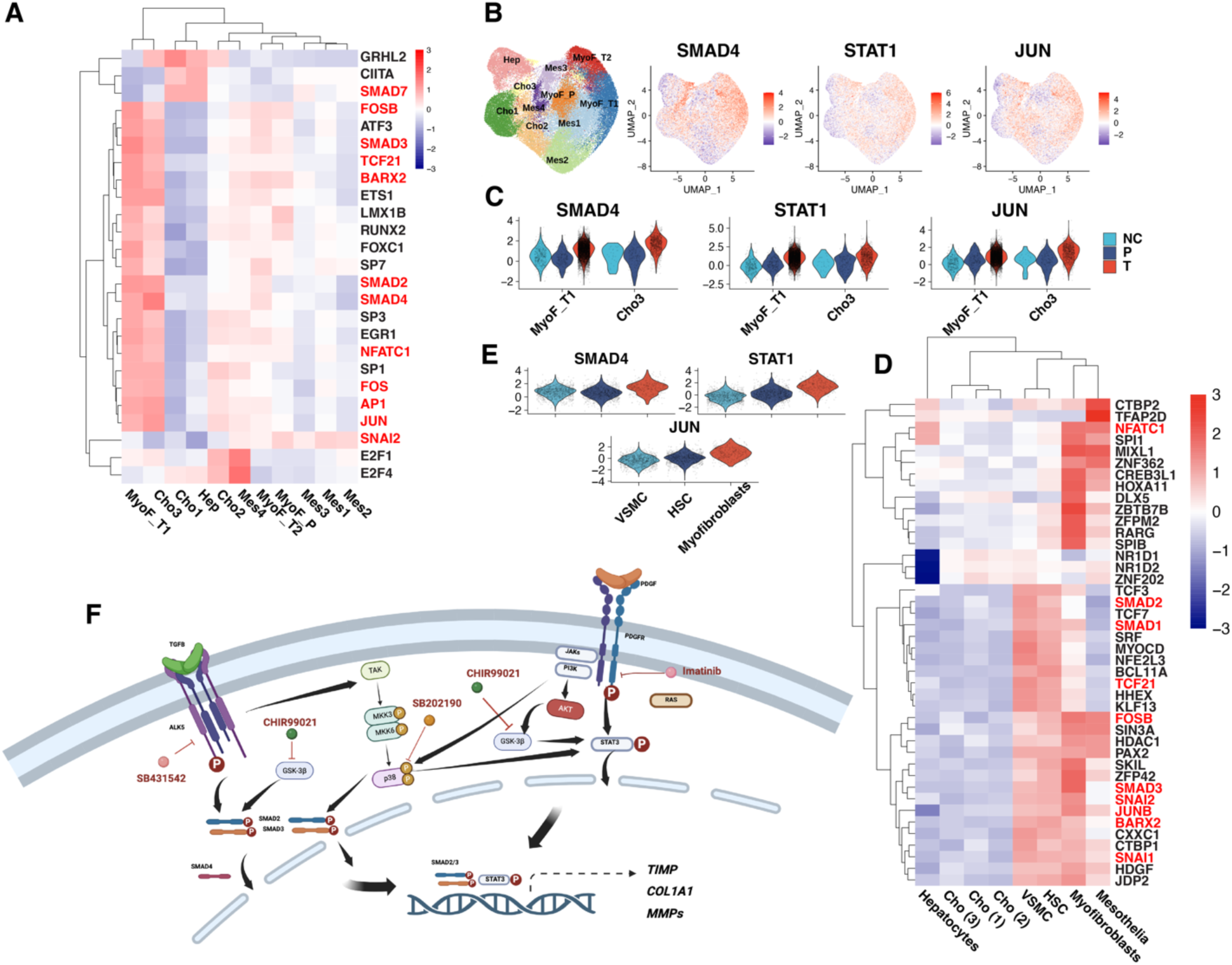
The transcription factors (TFs) activated in fibrotic microHOs and in fibrotic human liver. (**A**) A heat map shows the mean activity of the 25 TFs that were identified by decoupleR as being the ones that were most activated in fibrotic microHOs. The scaled mean activity for each TF in a cell cluster is indicated by the color of the square. Of note, MyoF_T1 and Cho3 had highest level of activation of these TFs. (**B**) SMAD4, STAT1 and JUN activity was plotted on the UMAP, and the cluster regions are indicated on the UMAP shown on the left. The scaled mean activity for each TF is indicated by dot color. The MyoF_T1 cluster had the highest level of activity of these TFs. (**C**) Violin plots show the TGFβ-induced increased TF activity in the MyoF_T1 and Cho3 clusters. (**D**) A heat map shows the mean activity of the 42 TFs that were identified by decoupleR as being the ones that were most activated in fibrotic human liver. The scaled mean activity for each TF in a cell type is indicated by the color of the square. Myofibroblasts have the most activated TFs, and there is overlap with the TFs activated in MyoF1 in fibrotic microHOs (highlighted in red). (**E**) Violin plots show that myofibroblasts have an increased level of SMAD4, STAT1 and JUN activity versus that of vascular smooth muscle cells or HSC. (**F**) Potential targets for anti-fibrotic drugs within the intracellular signaling pathways that are activated by TGFβ or PDGF. Pro-fibrotic agents activate the SMAD and STAT pathways, which can also activate the p38 MAPK and GSK3β pathways as shown in the diagram. Several potential targets for anti-fibrotic agents within SMAD, STAT and interacting (p38 MAPK and GSK3β) pathways are also indicated in the diagram.

### Pharmacologic characterization of pro-fibrotic signaling pathways

Since PDGFRβ and TGFβR1 inhibitors could block the PDGFβ and TGFβ1-induced increase in COL1A1^+^ cells, respectively; the interaction between these intracellular signaling pathways was examined in microHOs. The TGFRβ inhibitor did not block the PDGFβ-induced increase, and the PDGFRβ inhibitor did not block the TGFβ1-induced increase in COL1A1^+^ cells (Fig. 3). Also, addition of both PDGFβ and TGFβ1 did not significantly increase the number of COL1A1^+^ cells in microHOs relative to that caused by either factor alone (Fig. 3, bottom). Although the anti-fibrotic effect of the receptor tyrosine kinase inhibitors was growth factor dependent, other signaling pathways that are jointly activated by the TGFβ1 and PDGF receptors could be essential for fibrosis. Therefore, we examined whether co-administration of PI3K-Akt-mTOR (rapamycin) or MAPK pathway (SB202190) inhibitors with PDGF or TGFβ1 on day 13 could reduce fibrosis. Co-administration of 10 uM SB202190 – an ATP pocket binding and highly selective p38α and β kinase inhibitor ^58^– inhibited both TGFβ1 and PDGF-induced fibrosis in microHOs (**Fig. 7A-C**). Thus, p38 MAPK pathway activity is essential for either PDGF- or TGFβ1-driven fibrosis. Rapamycin partially inhibited PDGF-induced fibrosis but did not inhibit TGFβ1-induced fibrosis (**Fig. 7D-F**). The lack of inhibition of TGFβ1-induced fibrosis by rapamycin indicated that the downstream part of the PI3K-Akt-mTOR pathway was not essential for TGFβ1-induced fibrosis, but it may be required for PDGF-induced fibrosis.

**Figure 7.**
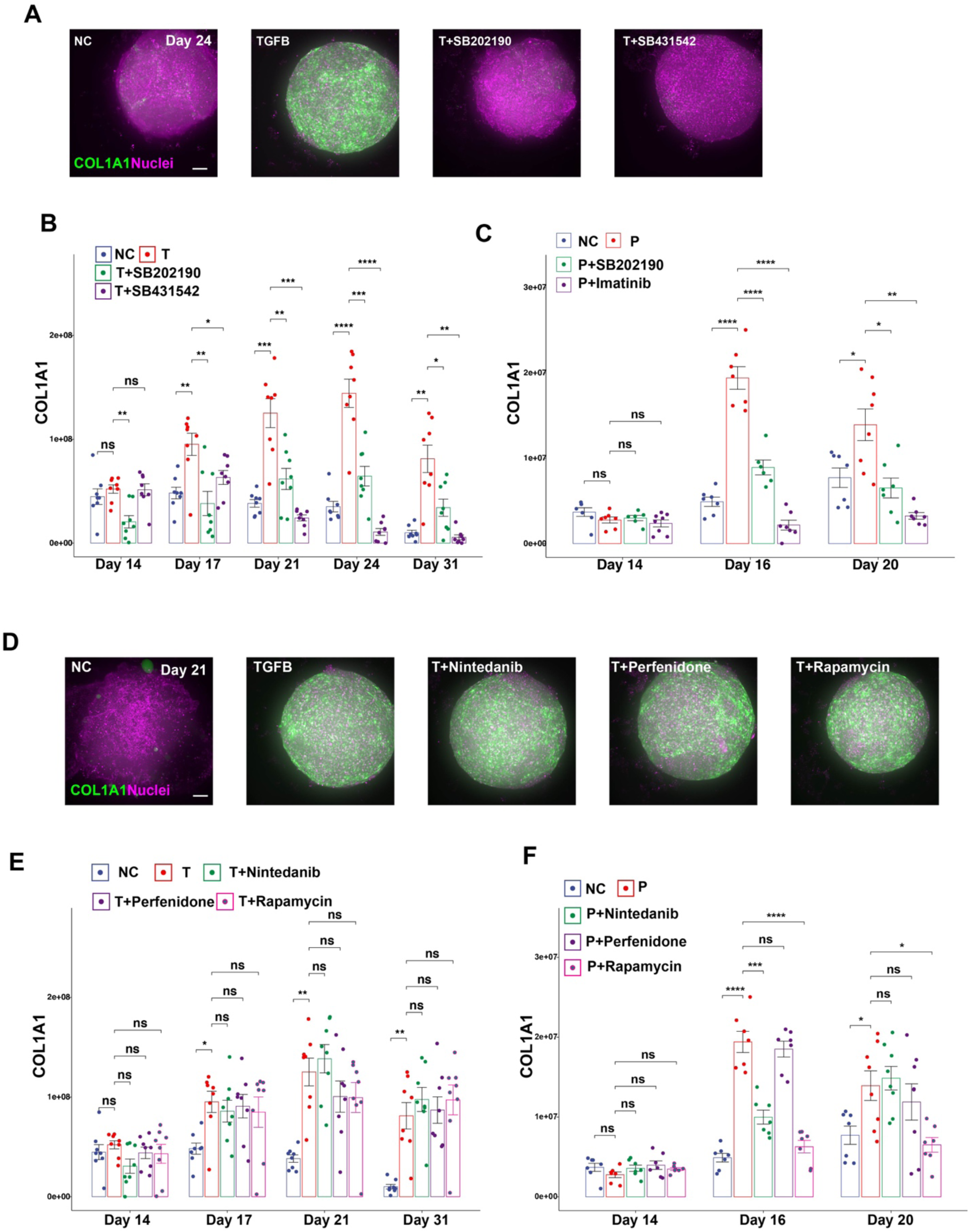
TGFβ1- or PDGF-induced fibrosis in microHOs is inhibited by a p38 MAPK (SB202190) inhibitor (A-C); but only the PDGF-induced fibrosis is inhibited by pirfenidone or rapamycin (D-F). *COL1A1*-P2A Clover microHOs were formed by adding 10,000 HBs to each microwell. Then, either nothing (NC), 50 ng/ml TGFβ1 (T) or 50 ng/ml (PDGF) was added on day 13 to each microwell in the presence or absence of 10 uM of the following drugs: SB431542 (TGFRβ inhibitor), SB202190 (p38 α inhibitor), nintedanib, pirfenidone or rapamycin. (A, D) Maximum intensity projection (MIP) images of z stack sections obtained from microHOs treated with no addition (NC), or 50 ng/ml TGFβ ± 10 uM of the indicated drugs. Scale bar 100 um. Clover expression is green, and nuclei stained with Hoechst 33342 are purple. (B-C, E-F) The amount of COL1A1^+^ cells within a microHO was serially measured on days 14 through 31. Each dot represents a measurement made on a microHO, the thick line is the median of 8 microHOs that were assessed, and the box plot shows the 25 to 75% range for all measurements per condition. When only growth factors were added, the p-values were relative to the NC, and the significance indicators are ns, not significant; *, p<0.05; **, p<0.01; ***, p<0.001 or ****, p<0.0001. When growth factors and inhibitors were added, the p-values were calculated relative to the PDGFβ or TGFβ1 treated microHOs (without added drug). A two-way ANOVA analysis indicates that drug treatment and time are two variables that have a significant interaction on the fluorescence measurements (Table S9).

WNT/β-catenin signaling contributes to the pathogenesis of liver fibrosis and NASH ^59, 60^, which led to the suggestion that it could be targeted by anti-fibrotic therapies ^61^. Since glycogen synthase kinase 3β (GSK3β)-mediated phosphorylation of β-catenin targets it for ubiquitin-dependent proteasomal degradation, GSK3β inhibitors will stabilize β-catenin and promote the transcription of β-catenin target mRNAs ^62, 63^. Therefore, we examined the effect that co-administration of a Wnt/β-catenin pathway activator (Wnt3a) or of a GSK3β enzyme inhibitor (10 uM CHIR99021), which has been shown to activate the Wnt/β-catenin pathway ^64^, had on fibrosis in microHOs. Wnt3a co-administration had a minimal effect on the extent of PDGF- or TGFβ1-driven fibrosis (Fig. S3). The GSK3β inhibitor by itself did not increase COL1A1^+^ cells in the organoids (Fig. S3). However, co-administration of the GSK3β inhibitor with either PDGF or TGFβ1 blocked fibrosis in microHOs (Fig. 3). Of importance, cell viability in microHOs was not reduced by addition of TGFβ1, nor by adding the GSK3β or TGFRβ inhibitors **(Fig. S16).** Our results indicate that p38 MAPK or GSK3β kinase inhibitors can block liver fibrosis irrespective of whether it is driven by either PDGF or TGFβ1.

This microHO system could also be used to test the potential anti-fibrotic efficacy of candidate medications for liver fibrosis. For example, nintedanib and pirfenidone are the only two FDA-approved drugs for treatment of pulmonary fibrosis ^65^. Both inhibit multiple tyrosine kinases and have been shown to reduce lung fibrosis in animal models ^66, 67, 68^ and in patients with interstitial lung fibrosis ^69, 70, 71, 72, 73^. Although the mechanism for their anti-fibrotic effect is unknown, it has been suggested that they could also be used to treat liver fibrosis ^54^. However, co-administration of 10 uM pirfenidone or 10 uM nintedanib did not inhibit the TGFβ1-induced fibrosis in microHOs. While pirfenidone did not inhibit PDGF-induced fibrosis, nintedanib delayed the onset of PDGF-induced fibrosis in microHOs (Fig. 7D-F). Of note, these concentrations were >10-fold above the nintedanib concentration that inhibited PDGF-stimulated cellular proliferation (IC_50_ 64 nM) and PDGFR auto-phosphorylation (IC_50_ 22-39 nM) ^74^ and of the pirfenidone concentration (1 uM) that inhibited proliferation and TGFb mRNA expression ^75^ in cultured human fibroblasts. The partial inhibitory effect of nintedanib on PDGF-induced fibrosis is consistent with its inhibition of the PDGF receptor tyrosine kinase activity (IC_50_ 60 nM) ^76^. WNT/β-catenin signaling contributes to the pathogenesis of liver fibrosis and NASH ^59, 60^, which led to the suggestion that it could be targeted by anti-fibrotic therapies ^61^. Since glycogen synthase kinase 3β (GSK3β)-mediated phosphorylation of β-catenin targets it for ubiquitin-dependent proteasomal degradation, GSK3β inhibitors will stabilize β-catenin and promote the transcription of β-catenin target mRNAs ^62, 63^. Therefore, we examined the effect that co-administration of a Wnt/β-catenin pathway activator (Wnt3a) or of a GSK3β enzyme inhibitor (10 uM CHIR99021), which has been shown to activate the Wnt/β-catenin pathway ^64^, had on fibrosis in microHOs. Wnt3a co-administration had a minimal effect on the extent of PDGF- or TGFβ1-driven fibrosis (Fig. S3). The GSK3β inhibitor by itself did not increase COL1A1^+^ cells in the organoids (Fig. S3). However, co-administration of the GSK3β inhibitor with either PDGF or TGFβ1 blocked fibrosis in microHOs (Fig. 3). Of importance, cell viability in microHOs was not reduced by addition of TGFβ1, nor by adding the GSK3β or TGFRβ inhibitors **(Fig. S16).** Our results indicate that p38 MAPK or GSK3β kinase inhibitors can block liver fibrosis irrespective of whether it is driven by either PDGF or TGFβ1.

This microHO system could also be used to test the potential anti-fibrotic efficacy of candidate medications for liver fibrosis. For example, nintedanib and pirfenidone are the only two FDA-approved drugs for treatment of pulmonary fibrosis ^65^. Both inhibit multiple tyrosine kinases and have been shown to reduce lung fibrosis in animal models ^66, 67, 68^ and in patients with interstitial lung fibrosis ^69, 70, 71, 72, 73^. Although the mechanism for their anti-fibrotic effect is unknown, it has been suggested that they could also be used to treat liver fibrosis ^54^. However, co-administration of 10 uM pirfenidone or 10 uM nintedanib did not inhibit the TGFβ1-induced fibrosis in microHOs. While pirfenidone did not inhibit PDGF-induced fibrosis, nintedanib delayed the onset of PDGF-induced fibrosis in microHOs (**Fig. 7D-F**). Of note, these concentrations were >10-fold above the nintedanib concentration that inhibited PDGF-stimulated cellular proliferation (IC_50_ 64 nM) and PDGFR auto-phosphorylation (IC_50_ 22-39 nM) ^74^ and of the pirfenidone concentration (1 uM) that inhibited proliferation and TGFb mRNA expression ^75^ in cultured human fibroblasts. The partial inhibitory effect of nintedanib of PDGF-induced fibrosis is consistent with its inhibition of the PDGF receptor tyrosine kinase activity (IC_50_ 60 nM) ^76^.

To determine if the anti-fibrotic effects of the TGFRβ, GSK3β or p38 inhibitors was dependent upon the genetic background of the *COL1A1*-P2A Clover line, we examined their anti-fibrotic effect in microHOs generated from iPSC lines (C1, C2) prepared from two other donors with different genetic backgrounds ^11^. As visualized by trichrome staining, TGFβ induced a marked increase in collagen-rich connective tissue in the C1 and C2 microHOs (p<0.001) that was markedly inhibited by addition of TGFRβ, GSK3β, or p38 inhibitors (p<0.001) (**Fig. 8**); which indicates that anti-fibrotic effect of these drugs was independent of genetic background.

**Figure 8.**
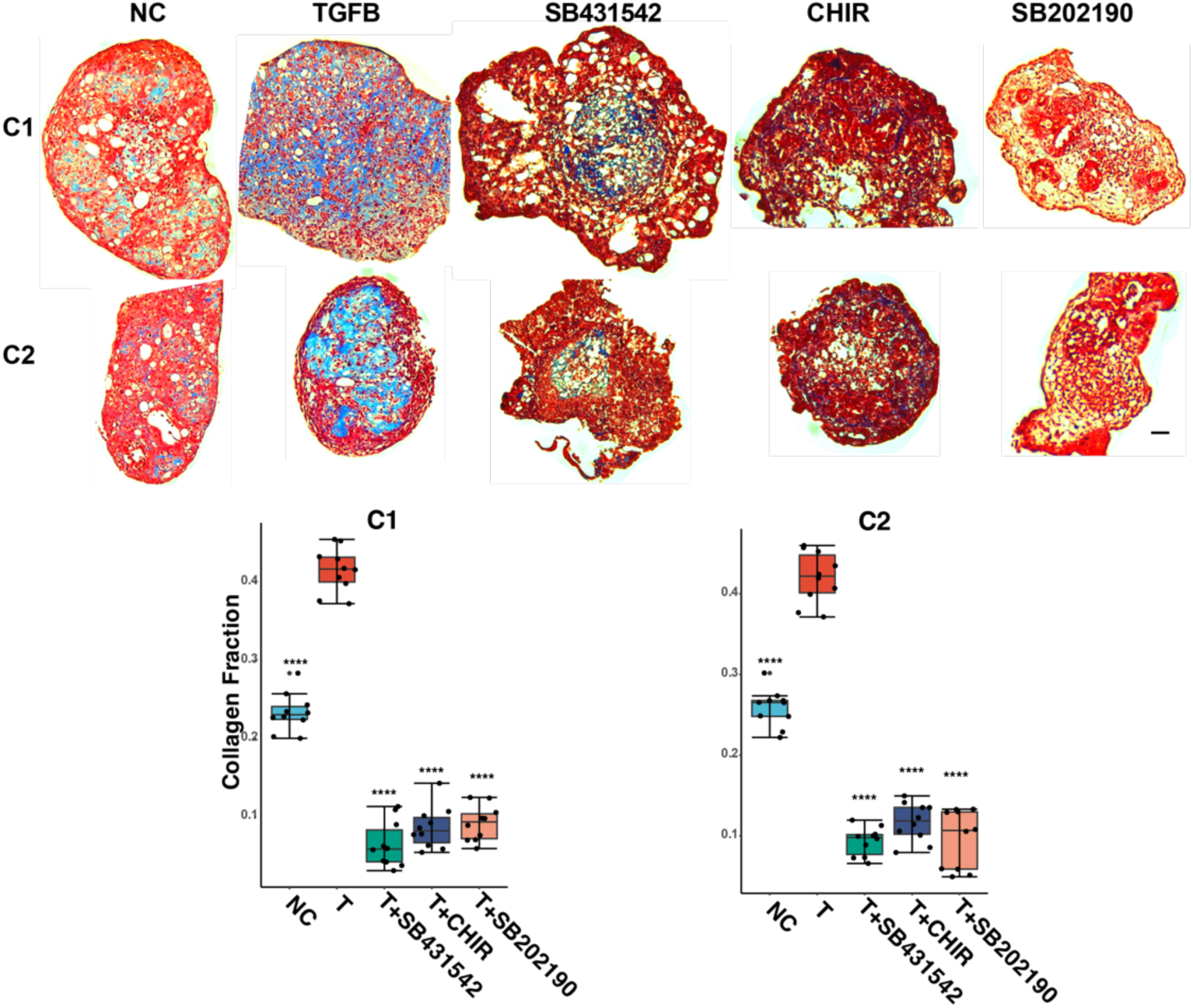
Anti-fibrotic drugs inhibit fibrosis in microHOs produced from iPSC of different genetic backgrounds. microHOs produced from iPSC lines generated from two donors (C1, C2) ^11^ were treated with no addition (NC), TGFβ (T) (50 ng/ml), or TGFβ and one of the following drugs on day 13: TGFBR inhibitor (10 μM SB431542), GSK3β inhibitor (3 μM CHIR) or a p38 inhibitor (10 μM SB202190). The microHOs were analyzed by trichrome staining on day 21. **(A)** Images of trichrome-stained TGFβ-treated microHOs show a marked increase in collagen-rich connective tissue (blue stained regions) relative to control (NC) microHOs, which only had a thin layer of connective tissue. The TGFβ-induced increased in collagen was markedly inhibited by addition of the TGFBR, GSK3β, or p38 inhibitors. The scale bar on the left: 50 μm. (**B**) Boxplots show the area in microHOs that received indicated treatments (n=6-9 per group) occupied by collagen (collagen fraction). The means for each group were compared using a one-way ANOVA and Tukey’s post-test: ****, p<0.0001 (for T vs NC, or T vs T+drug).

## Discussion

We demonstrate that microHOs and the live cell imaging system developed here can be used to characterize the biochemical signals and intracellular signaling pathways that drive liver fibrosis and for assessing potential anti-fibrotic therapies. The TGFβ1 and PDGF-induced fibrosis in microHOs was shown to resemble human fibrotic liver disease based upon the formation of thick collagen fibers (SHG results), increased collagen crosslinking (4-OHPro measurement) and increased collagen deposition (trichrome staining) in microHOs; and by transcriptomic comparisons demonstrating that the transcriptomes of microHO myofibroblasts resembled the myofibroblasts in fibrotic human liver tissue. Of the twenty agents tested using microHOs – which included cytokines, growth factors, fatty acids, a hedgehog agonist, and Wnt ligands – only PDGF and TGFβ1 had a strong pro-fibrotic effect. For several of the cytokines, growth factors and fatty acids tested, the absence of a pro-fibrotic effect in the organoid could be due to the absence of an immune system and/or of other cell types that could be required for a pro-fibrotic effect. However, a key finding emerging from this study is that the anti-fibrotic efficacy of PDGFR and TGFRβ tyrosine kinase inhibitors was dependent upon the factor driving the fibrosis, and this could explain why it has been so difficult to develop therapies for liver fibrosis. When fibrosis is driven by PDGFRβ-STAT3 pathway activation – as occurs in ARPKD liver fibrosis ^12^ or in murine models of lung and bone marrow fibrosis ^77^ – PDGFR inhibitors will exhibit anti-fibrotic efficacy; but they will be less effective when the fibrosis is driven by other mechanisms. There are organ-specific and species-specific differences in the mechanisms mediating tissue fibrosis ^10^ and different types of liver injury produces different patterns of liver fibrosis in humans ^78^. Consistent with differences in the factors that drive different types of fibrosis, microHO data indicates that at least one (and possibly both) of the FDA approved drugs for treatment of lung fibrosis drugs may not be effective for liver fibrosis. As another example, a combination treatment with PDGFR (imatinib) and TGFRβ (galunisertib ^79^) inhibitors was more effective than blockade of either pathway alone in a murine radiation-induced lung fibrosis model ^80^. To optimally treat liver fibrosis, therapies may have to be adjusted based upon the pathogenic driving factors, which may vary in different patients or in response to different inciting causes. There is already evidence that Diabetes patients can be subdivided into distinct sub-groups based upon clinical features and biomarker results ^81^, and efforts are underway to optimize treatment selection for the different types of diabetes patients ^82^. Just as in other diseases, application of the principles of ‘*precision medicine’* (i.e., using genetic or genomic information to optimize treatment selection ^83^) could enable therapies for liver fibrosis to be successfully developed.

Another key finding was that GSK3β and p38 MAPK inhibitors potently blocked the fibrosis induced by TGFβ1 or PDGF. The GSK3β inhibitor effect was unexpected since modulating Wnt/β-catenin signaling has had variable effects on liver fibrosis in mouse models ^84, 85^; and in a murine bile duct ligation model, GSK3β inhibitor administration produced an increase the extent of liver fibrosis ^86^. As discussed in supplemental note 2, there are multiple nodes where the TGFβ, Wnt/β-catenin and the p38 MAPK pathways interact. These interactions could explain how a GSK3β inhibitor could block TGFβ-induced fibrosis; but the mechanism for the GSK3β inhibitor effect on PDGF-induced fibrosis is less clear. Nevertheless, our results indicate that the p38 MAPK pathway, which is activated by both TGFβ1 and PDGF receptors, plays a key role in fibrosis. Although there was no prior data demonstrating that it had an effect on liver fibrosis, the ability of SB202190 to inhibit fibrosis in microHOs is consistent with prior studies showing that this agent inhibited the development of renal interstitial ^87^ and corneal ^88^ fibrosis in animal models, and the conversion of human corneal fibroblasts into myofibroblasts *in vitro* ^89^. However, additional studies will have to be performed to characterize the mechanism(s) by which the Wnt/β-catenin and p38 MAPK pathways, along with the SMAD and STAT3 pathways that are also activated by TGFβ1 and PDGF receptors, jointly contribute to liver fibrosis. Moreover, microHOs can also be used in conjunction with other recently developed genomic methods to provide a deeper understanding of the mechanisms mediating liver fibrosis ^90^. In summary, live cell imaging of human microHOs has identified GSK3β and p38 MAPK inhibitors as potential new therapies for liver fibrosis, and it is likely that other new therapies and their mechanism of action could subsequently be identified using this system.

## Data availability

All raw and processed single cell RNA-seq data were deposited in the Gene Expression Omnibus (GEO) and are available under accession GSE228214.

## Abbreviations

ARPKD: Autosomal Recessive Polycystic Kidney Disease
ECM: extra-cellular matrix
GSEA: gene set enrichment analysis
HCS: high content screening
HO: hepatic organoid
HCC: hepatocellular carcinoma
HSC: hepatic stellate cells
iPSC: induced pluripotent stem cells
MAPK: Mitogen Activated Protein Kinase
NASH: non-alcoholic steatohepatitis
SHG: Second Harmonic Generation
TF: transcription factor.

## Supporting information

Supplemental information, tables and fig;ures

