## Supplemental information, tables and fig;ures for "Characterization of Pro-Fibrotic Signaling Pathways using Human Hepatic Organoids"

***Supplemental note 1***: *Supplemental Note 1*: Gene Set Enrichment Analysis (**GSEA**) [1] was used to investigate whether microHOs fibrosis resembled two commonly occurring forms of human liver fibrosis. To do this, genes whose expression levels were increased in MyoF_T1 (n=527) or MyoF_P (n=1716) relative to Mes2 were used to form myofibroblast-specific gene expression signatures. GSEA calculates a normalized expression score (**NES**), which indicates whether the signature genes were enriched in fibrotic or non-fibrotic liver tissue. GSEA was first performed using expression data obtained from early (non-fibrotic) stage 1 and late (fibrotic) stage 4 NASH liver tissue (GSE13525 [2]). Liver fibrosis develops as NASH liver disease advances; myofibroblast activation is key to its pathogenesis [3, 4, 5]; and the extent of liver fibrosis is the major determinant of NASH outcome [6, 7]. GSEA results indicate that the MyoF_T1 (NSE 1.56; false discovery rate (FDR) 0.0178) gene expression signature was very strongly associated with stage 4 NASH, but not with early (NES -0.85 FDR 1) stage 1 NASH liver (**Fig. S10A, Table S9)**). The MyoF_T1 (NES 1.71, FDR 0.005) signature was also associated with liver fibrosis caused by hepatocellular carcinoma, whereas it (NES -1.32, FDR 0.17) was not associated with non-fibrotic hepatocarcinoma liver tissue (GSE6764 [8]) (**Fig. S10B**). In contrast, The MyoF_P signature had a positive (but not statistically significant association) with stage 4 NASH (NES 0.45, FDR 0.14) and HCC-induced liver fibrosis (NES 0.36 FDR 0.06). Thus, GSEA analyses indicate that the MyoF_T1 signature in microHOs resemble those in two commonly occurring types of human liver fibrosis.

***Supplemental note 2***: *Interactions between the TGFβ, Wnt/β-catenin and p38 MAPK pathways*. The TGFβ1 and Wnt/β-catenin pathways have several interaction points. (i) TGF-β significantly enhances hepatic WNT-5A expression *in vivo* and in myofibroblasts *in vitro* [9]. (ii) The interdomain regions of SMAD transcription factors have two residues that are phosphorylated by GSK3β in the nucleus, and their phosphorylation state regulates SMAD transcriptional activity and turnover [10]. Hence, the ability of the GSK3β inhibitor to block TGFβ1-induced fibrosis in microHOs could (at least in part) be mediated by altering the phosphorylation state of these two sites, which could alter SMAD transcription factor activity. (iii) During development, receptor tyrosine kinase and Wnt/β-catenin pathway signals were shown to be integrated by the level of SMAD phosphorylation [11].

The Wnt/β-catenin and p38 MAPK pathways also interact at several nodes. (i) Wnt ligands have been shown to activate p38 MAPK activity [12]. (ii) GSK3β phosphorylation by p38 MAPK regulates its kinase activity. MAPK kinases have a Thr–Gly–Tyr dual phosphorylation motif within their kinase activation loop, and both sites must be phosphorylated to fully activate the kinase. The MAP2K kinases MKK3 and MKK6 are the major upstream kinases for p38 activation [13]. However, phosphorylation on $Ser^{389}$ on GSK3β is a major mechanism for downregulating its kinase activity and p38α was shown to inactivate GSK3β by this mechanism: p38-mediated phosphorylation of $Ser^{389}$ on GSK3β was shown to cause β-catenin accumulation and activation of the Wnt signaling pathway [12, 14]. By this mechanism, p38 MAPK inhibitors could reduce Wnt pathway activation, which could prevent the development of a fibrosis that was dependent upon Wnt signaling.

### Materials and Methods

***Chemicals.*** The growth factors, drugs, antibodies, and other staining reagents are shown in Tables S6-S7.

***iPSCs and HO generation.*** The human iPSC line (C3) used in this study was prepared as previously described [15]. The biopsy sample used to generate this iPSC line was obtained according to a protocol (number 10368) approved by the Institutional Review Board at Stanford. The CRISPR-assisted insertion tagging system (CRISPaint) [16] plasmid (pCRISPR-HOT-Clover-BlastR) was obtained from Addgene (Plasmid #138569; http://n2t.net/addgene:138569). A clover expression cassette (**P2A-Clover**) was inserted at the COOH terminus of the endogenous *COL1A1* gene of the iPSC line without a STOP code using a sgRNA (TTGGGATGGAGGGAGTTTAC). After Blasticidin selection, colonies obtained from single cells were picked and genotyped for the correct in-frame insertion using forward (actcccacgtggtaatgccc); and reverse (cttcagggtcagcttgccg) primers. iPSCs were differentiated into hepatic organoids (HOs) via culture in a series of media containing different growth factors using previously described methods [15].

***Second Harmonic Generation (SHG) microscopy***. SHG (collagen fibers) images from day 21 control, PDGF-treated and TGFβ1-treated HOs were collected on a multimodal nonlinear optical microscope setup, consisting of a modified inverted confocal microscope (Nikon, Ti2-E with C2 scanner and 60X water immersion objective, NA=1.27) and a picosecond-pulsed laser source (APE picoEmerald S, 2 ps pulse width, 80 MHz repetition rate, and 10 $cm^{-1}$ bandwidth). As previously described[15], SHG data collection was made in the epi-direction with two narrow bandpass filters (Semrock FF01-400/12, Thorlabs MF390-18) and one shortpass filter (Thorlabs FESH0500). For the quantitative analyses, the area of the collagen fibers (identified by Otsu thresholding) was calculated for each SHG image and divided by the total sample area, forming the collagen area fraction (in %). To determine the diameter of the collagen fibers, the local thickness of the fibers was evaluated by applying the ImageJ Local Thickness routine (see the Analyze menu in Fiji) [17] to the SHG data. The volume fraction of thick collagen fibers (diameter ≥ 3 μm) was estimated by dividing the number of pixels represented by local thickness values ≥ 3 μm by the total number of pixels of the stack. To test whether the volume fractions obtained for control and TGFβ1- or PDGF-treated organoids were significantly different, the Welch's t-test was applied (* $p<0.05$ and ** $p<0.01$).

***microHO generation.*** To produce microHOs, 50~60% confluent iPSC cultures were switched to an endoderm differentiation medium that consisted of DMEM/F12 + ITS (GIBCO) supplemented with 0.1 mM nonessential amino acids, 1 mM pyruvate and 2 mM L-alanyl-L-glutamine dipeptide (GlutaMAX). On days 1 to 2, 100 ng/ml Activin-A (Peprotech, Rocky Hill, NJ), 10 ng/ml BMP4 (Peprotech), 100 ng/ml bFGF (Peprotech), 3 mM CHIR99021 (Selleckchem, Houston, TX) and 10 uM LY294002 (Selleckchem) were added this medium. On day 3, 100 ng/ml Activin-A and 100 ng/ml bFGF were added the definitive endoderm differentiation medium. From days 4 to 9, 20 ng/ml FGF10 (Peprotech) and 20 ng/ml BMP4 were added to the hepatoblast medium, which consisted of which consisted of Advanced RPMI 1640 Medium (Gibco) that was supplemented with GlutaMAX and ITS. After day 9, the hepatoblasts were dissociated to single cells in Accutase (Invitrogen) medium with 10 uM Y-27632 (Santa Cruz Biotechnology, Dallas, Texas). Then, 1000 to 5000 cells/well were re-aggregated in low-cell-adhesion Nunclon™ Sphera™ 96 well Microplates (ThermoFisher) that contained serum-free HO growth and differentiation medium, which consisted of William's E medium supplemented with 0.1% Polyvinyl alcohol (Sigma-Aldrich), 0.1 mM nonessential amino acids, 1 mM pyruvate, 2 mM L-alanyl-L-glutamine dipeptide (GlutaMAX), 10 mM Y-27632, 100 ng/ml EGF, 10 ng/ml HGF, 10 μM Dexamethasone and 10 μM Hydrocortisone (Sigma, St. Louis MO). On day 13, 50 ng/ml of TGFβ1 (Peprotech, 100-21; or Sinobiological, 10804-HNAC) or 50 ng/ml of PDGFB (Miltenyibiotec, 130-108-163; or Peprotech, 100-14B) was added to each microwell.

***microHO analysis***. For COL1A1$^{+}$ and Hoechst 33342 (Invitrogen, H3570) fluorescence measurements, 3D stacked images were captured from day 14 to day 31 cultures using a

Molecular Devices ImageXpress Micro Confocal system. For confocal imaging20 to 40 planes crossing the whole organoid at 20 um intervals were captured using a 10X 0.45 Plan Apo 4mm WD objective with a 60-um pinhole. For each 10X confocal z-stack field (enough to cover one whole organoid), maximum intensity projections (MIPs) were generated from all of the acquired z-planes (the number planes ranged from 20 to 40 depending on the batch size). MIPs from each channel belonging to the same well were used for organoid segmentation and feature extraction by applying the machine learning based 'Trainable Weka Segmentation' [18] plugin from FIJI [19]. Each individual organoid edge was automatically segmented based on the nuclei signal (Hoechst 33342). Cell numbers were determined using the 'Find Maxima' function in defined area of an organoid after segmentation. COL1A1-P2A-Clover$^{+}$ cells were segmented based on fluorescence. For training purposes, each analyzed element in every image (total 12 images) were used to train the classifiers by manual labelling all positive spots. Then, for experimental image analysis, the saved classifier was used to generate the probability map for each image. Segmented areas were then isolated, thresholded and binarized, and the Integrated Density was calculated from segmented area. An unpaired Student's t-test was used to test whether the measurements were significantly different between each comparison group. The macro script used for this analysis is available upon request.

***Immunohistochemistry.*** Fresh organoids were harvested and fixed in 4% paraformaldehyde for 30 minutes. All organoids were either stained directly or embedded in low melting point agarose (IBI Scientific, Dubuque, Iowa). The primary antibodies used for staining are listed in Table S5. The organoids were stained with the primary antibody, and permeabilized by incubation with 1% Triton X-100 (Sigma-Aldrich, St. Louis, MO) and 10% chicken serum (Jackson ImmunoResearch, Bar Harbor, MA) overnight. Then, secondary antibody-staining was performed using an Alexa Fluor labeled chicken anti IgG (H+L), which was cross-adsorbed with a secondary antibody in 10% chicken serum (Invitrogen, Pleasanton, CA).

***4-hydroxyproline quantitation.*** iPSC and day 9 hepatoblast cultures and day 21 control, PDGF- or TGFβ-treated HOs (n=16 per condition) were isolated and snap frozen in liquid nitrogen. 4-hydroxy proline levels were measured using the Hydroxyproline Assay kit (Sigma, MAK008) according to the manufacturer's instructions. In brief, HOs were homogenized in 100 µL of water and transferred to a pressure-tight polypropylene vial. Then, 100 µL of concentrated hydrochloric acid (HCl,12 M) was added, a cap was placed on tightly on the vial, and the sample was hydrolyzed at 120°C for 3 hours. The hydrolysate was centrifuged at 10,000 × g for 3 minutes. The clear supernatant was transferred and used for colorimetric assay of 4-hydroxyproline, which was performed according to the manufacturer's instructions.

***Trichrome staining***. Day 21 control, PDGF- or TGFβ- (+ drug) treated organoids were harvested, allowed to settle by gravity, and were embedded in low melting point agarose (IBI Scientific, Dubuque, Iowa). The embedded organoids were then processed by sectioning of the paraffin-embedded tissue to produce 10-micron tissue sections. To assess the amount of collagen in the organoids, trichrome staining of the tissue sections was performed using the MASSON'S 2000 TRICHROME STAIN KIT according to the manufacturer's instructions.

***Immunoblotting.*** iPSCs and day 9 hepatoblast cultures, and day 21 control, PDGF- or TGFβ-treated HOs (n=16 per condition) were lysed in RIPA buffer. The lysates were analyzed by PAGE on a 4-15% Tris-glycine gel. After blotting onto a membrane, the membranes were incubated with a 1:2000 dilution of an anti-GFP antibody (Clontech 632381, clone JL8) for detection of Clover protein, and the secondary antibodies were IRDye® 680 and 800 (LI-COR Biosciences).

***scRNA-Sequencing.*** Control, PDGFB- or TGFβ1-induced microHOs (n>30 per group) were harvested from multiple independently prepared sets of cultures, and single cell suspensions were prepared by protease digestion as described [15]. The single cell suspensions were visually inspected under a microscope, cells were counted using a Scepter™ 2.0 Handheld Automated Cell Counter (EMD Millipore, Burlington, MA), and then resuspended in PBS with 0.01% BSA. scRNA-seq libraries were prepared by using the spit-pool based scRNA-Seq method (Evercode™ WT Mini v2 from Parse biosciences) according to the manufacturer's instructions. In brief, ~5,000 cells from each group were split into 3 wells for the first round of sample barcoding, and the final pooled cells were divided into two equal libraries for sequencing. The expression matrix was generated using the 'Parse Biosciences analysis pipeline.' A total of 38,716 features that were generated from 13,160 cells were generated from four experimental batches, and 57, 7778 cells were recovered that passed quality control for processing of the scRNA-Seq data.

***Flow cytometry*** iPSCs were dissociated into single cells by incubation with 5 μM ETDA (Invitrogen) in PBS; and hepatoblasts were dissociated using Accutase (Gibco). Single cell suspensions were prepared from microHOs by protease digestion as described [15]. Flow cytometry was performed using at a BD Accuri C6 flow cytometer using the conjugated antibodies listed in Table S2, and the data was analyzed using FCS Express (denovosoftware.com) software. A density plot of forward scatter height (FSC-H) vs. forward scatter area (FSC-A) was used to exclude doublets.

***scRNA-Seq data analysis.*** The scRNA-Seq data was imported into 'Seurat' [20] for the subsequent analysis steps. (i) Cells with unique gene counts <200 or where the percentage of mitochondrial mRNAs was >10% were removed. (ii) A total of 2000 variable genes were identified using the default settings. Four batches of data were integrated by standard processing methods using the Seurat integration function. Unwanted sources of variation were removed by regression analysis, which was performed to mitigate the effect of signals caused by mRNAs with unique molecular identifiers and mitochondrial expression. (iii) Ten principal components were used to construct the shared nearest neighbor (SNN) graph, and the parameter regulating the resolution of the 'Find Clusters' program, was set to 0.5. This resulted in the identification of 11 unique clusters for all 57,778 cells. To identify the cell type of each cluster, differentially expressed genes for the pre-defined cell types were computed using the 'FindAllMarkers' function within the Seurat Package with the following parameters: only.pos = TRUE, min.pct = 0.25, logfc.threshold = 0.25. Seurat then identifies the differentially expressed genes using the non-parametric Wilcoxon rank sum test. The top 100 DEGs were used for the GO biological process analysis, which was performed using the 'clusterProfiler' for GO over-representation analysis [21].

***Module score calculation***. Module scores were calculated to assess the relationship between the transcriptomes of microHO clusters and the different types of cells in human liver tissue. scRNA-Seq data obtained from normal and cirrhotic human liver tissue (GSE136103) [22] was used to identify the different types of liver cells. Differentially expressed genes were calculated for each of the clusters in microHO (MyoF_T1, MyoF_P, Mes2) and human liver tissue (Myofibroblast, HSC, VSMC and Meso), using the 'FindMarkers' function of the R package in Seurat [23]. Then, each of the gene signatures used for determining the module score were selected based on the intersection between the marker genes in microHO and those in the cell types in human liver. There were 53, 84, and 31 shared genes that were used to calculate the module scores for MyoF_T1 vs MyoF in liver, MyoF_P and liver MyoF, and Mes2 and liver HSC, respectively. For comparisons of the scores obtained with vascular smooth muscle cells (VSMC), HSC, Myofibroblasts and Mesothelial cells, a one-way ANOVA was used to compare the means of the measurements for the four groups. The Tukey multiple comparison of the means test was then used to determine whether there was a significant difference between the means of all possible comparisons.

***Gene signature expression analyses (GSEA)***. The myofibroblast-specific gene expression signatures for genes whose expression was upregulated in MyoF_T1 (n=527) or MyoF_P (n=1716) relative to Mes2 (Table S*) were identified using the '*FindMarkers*' function in '*Seurat*' [20] with the parameter 'min.pct' set at 0.25 (which indicates the minimum fraction of cells within a cluster that expressed a given gene); and the default Wilcoxon rank sum test was used to perform this analysis. Two publicly available gene expression datasets for NASH (GSE135251 [2]) and cirrhotic (GSE6764 [8]) liver were obtained from the Gene Expression Omnibus using the '*GEOquery*' as described [24]. Cirrhotic liver tissue was obtained from 10 subjects undergoing liver resection for hepatocellular carcinoma at one of 3 US or European hospitals; and their liver tissues were classified as cirrhotic by the examining pathologist. The hepatocellular carcinoma samples (HCC) samples examined in this study were obtained from eight subjects whose liver was resected for HCC, but these specimens did not have fibrosis or cirrhosis according to the examining pathologist. The 10 normal liver tissues (used for comparison) were obtained from 10 subjects undergoing liver resection for other reasons at the same hospitals, and their liver tissue was classified as normal by the examining pathologist [8]. For the NASH analysis, 216 liver biopsies were examined by histology, which generated the fibrosis stage scores: 4 biopsies were classified as Stage 4, 48 biopsies were classified as Stage 1, and 46 as stage 0.

The GSEA for the myofibroblast signatures were calculated for the gene expression datasets generated from these samples using previously described methods [1]; and 1000 permutations were used for significance assessment for each analysis. The Enrichment score (ES) reflects the degree to which the Myofibroblast gene set is overrepresented at the top or bottom of a ranked list of genes; the Normalized enrichment score (NES) was used to compare analysis results across gene sets; and the false discovery rate (FDR) was used to estimate the probability that a gene set with a given NES represents a false positive finding.

***2-way analysis of variance (ANOVA) calculations***. The 2-way ANOVA was used to compare the means of the COL1A1:Clover signal (calculated as area * mean intensity (IntDen)) between the different drug treatment groups and time points, while also assessing interaction effects between the treatment groups and time. The two-way ANOVA was performed using the 'aov' function of the 'R\stats' package, and both groups and time points were analyzed as categorical independent variables. The null hypothesis was that there is no difference in the means of the different treatment groups or timepoints, and that the treatment groups and time points do not interact in any way. Since the two-way ANOVA of the data shown in Figures 2D, 3A, 3B, 5B, and 5C use separate datasets, a correction for multiple testing is not needed.

***RNA velocity, Cytotrace2, cell-cell communication and TF activity analysis***. RNA velocity analysis infers cell state by measuring the ratio of un-spliced to spliced mRNA transcripts, which provides information about the transcriptional activity of a gene and its direction. The transcript assignment file (tscp_assignment.csv.gz from the Parse pipeline output directory) contains the splicing information for each transcript identified in a Parse assay. The splicing information is used to generate the splice matrices that are required to run scVelo [25]. The Anndata file, which is required for scVelo, was generated using the Seurat object that contains the metadata, and splicing matrix. RNA velocity was estimated with the ‘stochastic model’ (using second-order moments). Cellular potency categories and the absolute developmental potential was assessed with ‘CytoTRACE 2’ [26] using the scRNA-seq data. The predicted potency scores provide a continuous measure of developmental potential; and they range from 0 (differentiated) to 1 (totipotent). The raw count from Seurat object is used to directly to compute CytoTRACE score. The R toolkit ‘CellChat’ v2 was used for inference, visualization and analysis of cell-cell communication based upon the scRNA-seq data [27]. The All CellChatDB, except for the "Non-protein signaling" component, was used for analysis of cell-cell communication. The R implementation of ‘decoupleR’ [28] was used to extract biological activities from the CollecTRI database. CollecTRI is a comprehensive resource containing a curated collection of TFs and their transcriptional targets compiled from 12 different sources [29]. The interactions are weighted based upon their mode of regulation (activation or inhibition).

***Analysis of drug effect in C1 and C2 microHOs.*** The C1 and C2 iPSC lines were generated and characterized as previously described [15] and used to prepare microHOs as described above. Trichrome staining of day 21 microHOs was performed according to the manufacturer’s instructions using MASSON'S 2000 TRICHROME stain (Americanmastertech, Lodi, CA). Image segmentation and quantitative measurement of collagen rich areas was performed using our previously described methods [30]. In brief, Fiji (2.1.0) implementation of ImageJ was used to quantify the areas of positive staining. The ‘Trainable Weka Segmentation’ plug-in was used to train the classifiers and calculate the test experimental image. Statistical analysis was performed using a one-way ANOVA to assess the overall differences between group means, and Tukey's post-test was then used for pairwise comparisons to identify groups with significant differences in their means.

**Table S1**. Drugs and chemicals used in these studies.

| Reagent | Company | Catalog # |
|---|---|---|
| palmic acid | SigmaAldrich | P5585 |
| Oleic acid | SigmaAldrich | O1383 |
| Stearic acid | SigmaAldrich | S4751 |
| Myristic Acid | Caymanchem | 13351 |
| Myristoleic Acid | Caymanchem | 9002461 |
| Palmitoleic Acid | Caymanchem | 10009871 |
| Elaidic Acid | Caymanchem | 90250 |
| CDCA | SigmaAldrich | C9377 |
| Activin A Protein | Sinobiological | 10429-HNAH |
| Animal-Free Recombinant Human EGF | Peprotech | AF-100-15 |
| EGF | Sinobiological | 10605-HNAE |
| FGF4 | Peprotech | 100-31 |
| HGF | Sinobiological | 10463-HNAS |
| FGF10 | Sinobiological | 10573-HNAE |
| FGF2 Protein, Human, Recombinant | Sinobiological | 10014-HNAE |
| FGF10 | Peprotech | 100-26 |
| Human PDGF-BB IS | Miltenyibiotec | 130-108-163 |
| Human VEGF (165) IS | Miltenyibiotec | 130-109-383 |
| Recombinant Human PDGF-BB (carrier-free) | Biolegend | 577302 |
| Recombinant Human PDGF-BB | Peprotech | 100-14B |
| Recombinant Human PDGF-BB Protein, CF | rndsystems | 220-BB-010In Stock |
| TGF beta 1 Protein, Human, Rhesus, Cynomolgus, Canine, Recombinant | Sinobiological | 10804-HNAC |
| Recombinant Human BMP-4 | Peprotech | 120-05 |
| Recombinant Human BMP-4 (carrier-free) | Biolegend | 795606 |
| Recombinant Human/Mouse/Rat Activin A | R&D | 338-AC-010 |
| Recombinant Human FGF-Basic (146 a.a.) | Peprotech | AF-100-18C |
| Wnt3a | Peprotech | 315-20 |
| OSM | Peprotech | 300-10 |
| IL3 Protein, Human, Recombinant (His Tag) | Sinobiological | 11858-H08H |
| IL6 | R&D | 7270-IL-010/CF |
| IL-13 Protein, Human, Recombinant | Sinobiological | 10369-HNAC |
| IL-33 Protein, Human, Recombinant | Sinobiological | 10368-HNAE |
| IGF1 | R&D | 291-G1-200 |
| R-Spondin 1 Protein | R&D | 4645-RS |
| JAG-1 protein | StemRD | Cat#: JAG-1-pep-100 |
| SB202190 (FHPI) | Selleckchem | S1077 |
| SB431542 | Selleck | S1067 |
| Sirolimus (Rapamycin) | Medkoo | 100766 |
| Imatinib mesylate | Medkoo | 100470 |
| DAPT (GSI-IX) | Selleckchem | S2215 |
| LDN-193189 | Selleckchem | S2618 |
| SAG | Selleckchem | S7779 |
| IWR1-en | Selleckchem | S7086 |
| A-83-01 | Santa Cruz Biotechnology | sc-203791 |
| PI-103 | Tocris | 2930 |
| PD0325901 | Santa Cruz Biotechnology | sc-205427 |
| CHIR99021 | Tocris | 4423 |
| Wnt-C59 | Tocris | 5148 |
| BIO | Selleckchem | S7198 |
| Thiazovivin | Medchemexpress | HY-13257 |
| Nintedanib | Tocris | 7049 |
| Pirfenidone | Tocris | 1093 |
| polyvinyl alcohol | SigmaAldrich | P8136 |
| Matrigel | BD | 354234 |

**Table S2**. Antibodies and staining reagents used in these studies.

| Name | Company | Catalog# | Dilution |
|---|---|---|---|
| CK8 | abcam | ab53280 | 1/100 - 1/250 |
| CK8 | DSHB | TROMA-I | 200 |
| COL1 | ABCAM | ab34710 | 1:100 |
| GFP Tag Polyclonal Antibody | thermofisher | A-11122 | 200-2000 |
| GFP Antibody (B-2) | scbt | sc-9996 | 50 |
| Human Albumin Antibody | Bethyl | A80-129A | 1:200 – 1:2,000 |
| PDGFRB | eBioscienc | 14-1402-82 | 100 |
| PDGFRB | ABCAM | ab32570 | 100 |
| PDGFRB | R&D | AF385-SP | 100 |
| PDGFRB | ll Signaling Technolo | 3169T | 100 |
| LipidSpot™ Lipid Droplet Stains | biotium | #70069-T | |
| MitoSox Red | Invitrogen | M36008 | |
| Alexa Fluor® 647 anti-human CD326 (EpCAM) Antibody | biolegend | 324212 | 1:20 |

| Cluster | Cell Type | NC (%) | PDGF (%) | TGFβ (%) |
|---|---|---|---|---|
| 0 | Mes1 | 20.7 | 20.5 | 20.2 |
| 1 | MyoF_T1 | 1.9 | 1.5 | 37.5 |
| 2 | Mes2 | 24.9 | 4.8 | 3.3 |
| 3 | Cho1 | 17.6 | 14.7 | 1.3 |
| 4 | Hep | 12.8 | 12.3 | 5.2 |
| 5 | MyoF_T2 | 3.6 | 7.07 | 16.2 |
| 6 | Cho2 | 13.5 | 7.1 | 0.7 |
| 7 | MyoF_P | 0.8 | 20.1 | 1.0 |
| 8 | Mes3 | 2.2 | 6.5 | 5.7 |
| 9 | Mes4 | 1.9 | 4.4 | 3.0 |
| 10 | Cho3 | 0.1 | 1.0 | 6.0 |

**Table S3**. The cell types and the percentage of the total number of cells in the 11 cell clusters identified by analysis of scRNA-Seq data in day 21 control, PDGF-, and TGFβ1-treated microHOs are shown. The transcriptomes of the 10 other cell clusters in the day 21 microHOs were defined using canonical markers and by comparison with the cells in control and cirrhotic human livers. Based upon the expression of canonical mRNAs and level of concordance between the cell cluster transcriptomes and the cell types identified in human liver, the microHOs have cholangiocytes (Cho1-3), hepatocyte (Hep), mesenchymal cell (Mes1-4) and myofibroblast (MyoF_T1-2, MyoF_P) clusters. A myofibroblast cluster (MyoF_T1) was far more abundant in the TGFβ1-treated microHOs than in control or PDGF-treated microHOs (37.5% of total vs <2%), while another myofibroblast cluster (MyoF_P) was more abundant in the PDGF-treated microHOs than in control or TGFβ1-treated microHOs (20% vs <2% of the total number of cells).

**Table S4** is provided at the end of this file.

**Table S5.** The results of 1-way ANOVA analyses of the cell percentages calculated by analysis of the scRNA-Seq data for the five biological replicates from the control, PDGF- and TGF$\beta$-treated microHO preps shown in Figure S8. The degrees of freedom (DF), the sum of the squares (Sum Sq) and mean squared (mean Sq) of the variance, calculated F-value, and the probability Pr(>F) from the ANOVA are shown. A Pr(>F) < 0.05 indicates that there was a significant effect of the indicated variable. There were significant differences in the % of MyoF_T1, Mes2, MyoF_P and Cho3 cells as indicated by the p-value: ***; 0.001, **; 0.01, *; 0.05. However, Cho3 abundance was $\leq$1% in all three types of microHOs.

| | Df | Sum Sq | Mean Sq | F value | Pr(>F) |
|---|---|---|---|---|---|
| Mes1 | 2 | 5 | 2.6 | 0.009 | 0.991 |
| Residuals | 12 | 3599 | 299.9 | | |
| | Df | Sum Sq | Mean Sq | F value | Pr(>F) |
| **MyoF_T1** | 2 | 1259 | 629.7 | 6.046 | **0.0153 *** |
| Residuals | 12 | 1250 | 104.1 | | |
| | Df | Sum Sq | Mean Sq | F value | Pr(>F) |
| **Mes2** | 2 | 747.4 | 373.7 | 8.941 | **0.00419 ** ** |
| Residuals | 12 | 501.6 | 41.8 | | |
| | Df | Sum Sq | Mean Sq | F value | Pr(>F) |
| Cho1 | 2 | 339.8 | 169.92 | 1.946 | 0.185 |
| Residuals | 12 | 1047.8 | 87.32 | | |
| | Df | Sum Sq | Mean Sq | F value | Pr(>F) |
| Hep | 2 | 134.6 | 67.28 | 0.974 | 0.405 |
| Residuals | 12 | 828.7 | 69.05 | | |
| | Df | Sum Sq | Mean Sq | F value | Pr(>F) |
| MyoF_T2 | 2 | 145.5 | 72.73 | 2.739 | 0.105 |
| Residuals | 12 | 318.7 | 26.56 | | |
| | Df | Sum Sq | Mean Sq | F value | Pr(>F) |
| Cho2 | 2 | 169.3 | 84.63 | 2.754 | 0.104 |
| Residuals | 12 | 368.7 | 30.73 | | |
| | Df | Sum Sq | Mean Sq | F value | Pr(>F) |
| **MyoF_P** | 2 | 425.9 | 213.0 | 3.988 | **0.047 *** |
| Residuals | 12 | 640.8 | 53.4 | | |
| | Df | Sum Sq | Mean Sq | F value | Pr(>F) |
| Mes3 | 2 | 61.31 | 30.66 | 2.503 | 0.123 |
| Residuals | 12 | 146.99 | 12.25 | | |
| | Df | Sum Sq | Mean Sq | F value | Pr(>F) |
| Mes4 | 2 | 22.8 | 11.40 | 1.045 | 0.382 |
| Residuals | 12 | 130.9 | 10.91 | | |
| | Df | Sum Sq | Mean Sq | F value | Pr(>F) |
| **Cho3** | 2 | 90.6 | 45.30 | 4.242 | **0.0404 *** |
| Residuals | 12 | 128.1 | 10.68 | | |

**Table S6.** The gene symbols for 3 sets of genes whose mRNAs were used for calculation of the module scores shown in Figure 4C. (i) The MyoF_T1 mRNAs were derived from the intersection of the differentially expressed marker genes for MyoF_T1 versus Mes2 clusters. (ii) The MyoF_P mRNAs were derived from the intersection of differentially expressed marker genes for MyoF_P vs Mes2 clusters. (iii) The Mes2 mRNAs were derived from the intersection of the differentially expressed marker genes for Mes2 vs MyoF_T1 clusters.

| | **MyoF_T1** | | | **MyoF_P** | | | **Mes2** |
|---|---|---|---|---|---|---|---|
| **1** | OGN | | 1 | PTGDS | | 1 | TM4SF1 |
| **2** | LTBP2 | | 2 | LIFR | | 2 | NPR3.00 |
| **3** | LTBP1 | | 3 | OGN | | 3 | TXNIP |
| **4** | IGFBP7 | | 4 | GPNMB | | 4 | EPAS1 |
| **5** | MMP2 | | 5 | LGALS3BP | | 5 | ARHGAP29 |
| **6** | DPYSL3 | | 6 | PLTP | | 6 | INPP4B |
| **7** | ITGA1 | | 7 | THY1 | | 7 | MEF2C |
| **8** | COL4A2 | | 8 | MATN2 | | 8 | FRZB |
| **9** | COL4A1 | | 9 | ITGA1 | | 9 | ALDH1A1 |
| **10** | GSN | | 10 | LTBP1 | | 10 | RERG |
| **11** | IGFBP4 | | 11 | COL4A1 | | 11 | SYTL2 |
| **12** | SERPINE2 | | 12 | COL4A2 | | 12 | PLA2G5 |
| **13** | COL6A3 | | 13 | TIMP1 | | 13 | ANGPT1 |
| **14** | TIMP1 | | 14 | SPARC | | 14 | ARHGAP15 |
| **15** | VCAN | | 15 | VCAN | | 15 | EBF1 |
| **16** | COL1A2 | | 16 | FN1 | | 16 | NRP1 |
| **17** | COL1A1 | | 17 | HGF | | 17 | AKAP12 |
| **18** | FN1 | | 18 | FBLN5 | | 18 | SLC40A1 |
| **19** | BGN | | 19 | GPC3 | | 19 | RAPGEF5 |
| **20** | THY1 | | 20 | IGFBP3 | | 20 | EZR |
| **21** | FMOD | | 21 | MGP | | 21 | SLC2A3 |
| **22** | PRSS23 | | 22 | PLXDC2 | | 22 | APOA2 |
| **23** | SERPINA1 | | 23 | COL6A3 | | 23 | NFASC |
| **24** | CYP1B1 | | 24 | EDNRB | | 24 | PDE1A |
| **25** | IGFBP6 | | 25 | DPT | | 25 | ALB |
| **26** | ELN | | 26 | PDGFRA | | 26 | HMGB2 |

| | | | | | | | |
|---|---|---|---|---|---|---|---|
| **27** | PDGFRA | | 27 | IGFBP7 | | 27 | APOA1 |
| **28** | SPARC | | 28 | LMCD1 | | 28 | ZBTB16 |
| **29** | F3 | | 29 | C7 | | 29 | HDAC2 |
| **30** | PDLIM3 | | 30 | KRT18 | | 30 | DNAJC15 |
| **31** | FBLN5 | | 31 | SERPINA1 | | 31 | NHSL2 |
| **32** | ECM1 | | 32 | NR2F1 | | | |
| **33** | DPT | | 33 | MASP1 | | | |
| **34** | FAP | | 34 | FMOD | | | |
| **35** | S100A6 | | 35 | NR2F1-AS1 | | | |
| **36** | NR2F1 | | 36 | MFAP4 | | | |
| **37** | MFAP4 | | 37 | PLPP3 | | | |
| **38** | ITGBL1 | | 38 | MIR99AHG | | | |
| **39** | SPON2 | | 39 | C3 | | | |
| **40** | C3 | | 40 | RAPGEF5 | | | |
| **41** | LMCD1 | | 41 | FGF7 | | | |
| **42** | THBS1 | | 42 | PTN | | | |
| **43** | KRT18 | | 43 | IL6ST | | | |
| **44** | PRELP | | 44 | MMP2 | | | |
| **45** | A2M | | 45 | PCDH9 | | | |
| **46** | HSPB6 | | 46 | FAP | | | |
| **47** | CFH | | 47 | THBS2 | | | |
| **48** | SOD2 | | 48 | GPC6 | | | |
| **49** | THBS2 | | 49 | S100A6 | | | |
| **50** | PLTP | | 50 | NRP1 | | | |
| **51** | IFI6 | | 51 | COL1A2 | | | |
| **52** | NR2F1-AS1 | | 52 | IGFBP6 | | | |
| **53** | EMP1 | | 53 | PTGIS | | | |
| | | | 54 | SPON2 | | | |
| | | | 55 | DPYSL3 | | | |
| | | | 56 | HSPB6 | | | |
| | | | 57 | ABCA8 | | | |

| | | | | |
|---|---|---|---|---|
| | | | 58 | SERPINE2 |
| | | | 59 | ARHGAP15 |
| | | | 60 | EMP1 |
| | | | 61 | F3 |
| | | | 62 | APOA2 |
| | | | 63 | PDLIM3 |
| | | | 64 | DNAJC15 |
| | | | 65 | APOA1 |
| | | | 66 | VCAM1 |
| | | | 67 | CFH |
| | | | 68 | CYP1B1 |
| | | | 69 | THBS1 |
| | | | 70 | ITGBL1 |
| | | | 71 | ANK3 |
| | | | 72 | PRSS23 |
| | | | 73 | LTBP2 |
| | | | 74 | LAMA2 |
| | | | 75 | ELN |
| | | | 76 | CFI |
| | | | 77 | SOD2 |
| | | | 78 | CD44 |
| | | | 79 | BGN |
| | | | 80 | DCN |
| | | | 81 | COLEC10 |
| | | | 82 | PRELP |
| | | | 83 | IGFBP4 |
| | | | 84 | A2M |

**Table S7.** One-way ANOVA and Tukey test for multiple comparisons of the module scores shown in Figure 4C. (**A**) The gene signatures for each cluster (Mes2, MyoF_T1, MyoF_P) were compared with four groups of human liver cells: vascular smooth muscle cells (VSMC); HSC; myofibroblasts; and mesothelia (Meso). The mean module score (Mean) and ratio of module score for each cell type relative to that with the highest mean (Fold) are shown. (**B, C**) Pairwise comparisons between HSC and three other types of human liver cells (VSMC, MyoF and Meso) produce significantly different results ($P<1\times10^{-10}$); and pairwise comparisons between MyoF and the three other types of human liver cells (VSMC, HSC and Meso) also produce significantly different results ($P<1\times10^{-10}$). Taken together, the ANOVA and pairwise comparison results indicate that the gene signature of myofibroblasts in liver is most similar to MyoF_T1 and MyoF_P in microHOs, and that the gene signature of HSC is most similar to Mes2.

**Table S7A.** The Mean of four liver cell types and different clusters and their relative Fold.

| **Cluster:** | **Mes2** | | **MyoF_T1** | | **MyoF_P** | |
|---|---|---|---|---|---|---|
| **Cell types** | **Mean** | **Fold** | **Mean** | **Fold** | **Mean** | **Fold** |
| VSMC | 0.12 | 2.49 | 0.18 | 5.11 | 0.11 | 6.86 |
| HSC | **0.31** | 1.00 | 0.32 | 2.91 | 0.25 | 3.09 |
| MyoF | 0.06 | 4.80 | **0.92** | 1.00 | **0.78** | 1.00 |
| Meso | -0.07 | -4.48 | 0.49 | 1.89 | 0.39 | 2.00 |

**Table S7B.** One-way ANOVA analysis for the four cell types in different clusters**.**

| **Cluster** | **Source of Variation** | **Sum of Squares** | **Degrees of Freedom** | **Mean Square** | ***F*-Calculated** | ***P*-Value** |
|---|---|---|---|---|---|---|
| **Mes2** | Cell Types | 28.54 | 3 | 9.51 | 371 | $<2x10^{-16}$ |
| | Residuals | 59.03 | 2299 | 0.026 | | |
| **MyoF_T1** | Cell Types | 137.27 | 3 | 45.76 | 2834 | $<2x10^{-16}$ |
| | Residuals | 37.12 | 2299 | 0.02 | | |
| **MyoF_P** | Cell Types | 110.5 | 3 | 36.83 | 3850 | $<2x10^{-16}$ |
| | Residuals | 22 | 2299 | 0.01 | | |

**Table S7C.** Tukey test for multiple comparisons of the module scores.

| **Cluster** | **Pairwise Comparison** | **Mean Difference** | ***P*-adjusted** | **95% Confidence Interval Lower Bound** | **Upper Bound** |
|---|---|---|---|---|---|
| **Mes2** | HSC *vs* VSMC | 0.19 | $<1\times10^{-10}$ | 0.17 | 0.20 |
| | MyoF *vs* HSC | -0.24 | $<1\times10^{-10}$ | -0.27 | -0.22 |
| | Meso *vs* HSC | -0.38 | $<1\times10^{-10}$ | -0.42 | -0.34 |
| **MyoF_T1** | MyoF *vs* VSMC | 0.74 | $<1\times10^{-10}$ | 0.72 | 0.76 |
| | MyoF *vs* HSC | 0.61 | $<1\times10^{-10}$ | 0.59 | 0.62 |
| | Meso *vs* MyoF | -0.44 | $<1\times10^{-10}$ | -0.47 | -0.34 |
| **MyoF_P** | HSC *vs* VSMC | 0.67 | $<1\times10^{-10}$ | 0.65 | 0.69 |
| | MyoF *vs* HSC | 0.53 | $<1\times10^{-10}$ | 0.51 | 0.55 |
| | Meso *vs* HSC | -0.39 | $<1\times10^{-10}$ | -0.41 | -0.36 |

**Table S8**. Summary of the GSEA results assessing the correlation between the MyoF_T1 and MyoF_P gene signatures with NASH **(A**) or hepatocellular carcinoma-associated (HCC) **(B**) non-fibrotic and fibrotic liver tissues. Genes whose expression levels were increased in MyoF_T1 (n=527) or MyoF_P (n=1716) relative to Mes2 were used to form myofibroblast-specific gene expression signatures. The expression data was obtained from (A) early (non-fibrotic) stage 1 and late (fibrotic) stage 4 NASH liver tissue (GSE13525 [2]) or (B) from resected HCC liver tissue, which was classified by pathologists as fibrotic or non-fibrotic (GSE6764 [8]). GSEA analyses revealed that the MyoF_T1 signature was strongly associated with stage 4 NASH, but not with early stage 1 NASH liver tissue; and with liver fibrosis caused by HCC, but not with non-fibrotic HCC liver tissue. The MyoF_P signature had positive associations with stage 4 NASH and HCC-induced fibrotic liver tissue, but those associations were not statistically significant. The expression score (ES), normalized expression score (NES), nominal p-value, and false discovery rate (FDR q-value) are shown for each comparison.

**A. NASH Analyses (GSE135251)**

Comparison: Stage 4 (Fibrosis)

| | ES | NES | P-value | FDR q-value |
|---|---|---|---|---|
| MyoF_T1 | 0.56 | 1.56 | 0.007 | 0.018 |
| MyoF_P | 0.45 | 1.32 | 0.10 | 0.15 |

Comparison: Stage 1 (No Fibrosis)

| | ES | NES | P-value | FDR q-value |
|---|---|---|---|---|
| MyoF_T1 | -0.30 | -0.85 | 0.66 | 1 |
| MyoF_P | -0.29 | -0.82 | 0.72 | 0.87 |

**B. HCC Analyses (GSE6764**)

Comparison: Fibrotic Liver

| | ES | NES | P-value | FDR q-value |
|---|---|---|---|---|
| MyoF_T1 | 0.51 | 1.71 | 0.0019 | 0.05 |
| MyoF_P | 0.36 | 1.48 | 0.03 | 0.06 |

Comparison: Non-fibrotic liver

| | ES | NES | P-value | FDR q-value |
|---|---|---|---|---|
| MyoF_T1 | -0.36 | -1.32 | 0.10 | 0.16 |
| MyoF_P | -0.34 | -1.34 | 0.05 | 0.22 |

**Table S9.** The results of 2-way ANOVA analyses of the data shown in the indicated figures are shown. For these analyses, time and treatment group was treated as categorical variables. The degrees of freedom (DF), the sum of the squares (Sum Sq) and mean squared (mean Sq) of the variance, calculated F-value, and the probability (Pr(>F) from the 2-way ANOVA are shown for the treatment group, time, and treatment-by-time analyses. A Pr(>F) < 0.01 indicates that there was a significant effect of the indicated variable.

| **Figure 2D** | **DF** | **Sum Sq** | **Mean Sq** | **F value** | **Pr(>F)** |
|---|---|---|---|---|---|
| Treatment | 3 | 6.11E+14 | 2.04E+14 | 110.79 | <2.00E-16 |
| Time | 4 | 1.47E+16 | 3.67E+14 | 19.98 | 4.92E-13 |
| Treatment by Time | 12 | 2.48E+14 | 2.07E+13 | 11.23 | 1.66E-15 |
| Residuals | 140 | 2.57E+14 | 1.84E+12 | | |

| **Figure 3A** | **Df** | **Sum Sq** | **Mean Sq** | **F value** | **Pr(>F)** |
|---|---|---|---|---|---|
| Treatment | 6 | 2.96E+14 | 4.93E+13 | 14.087 | 1.24E-12 |
| Time | 2 | 1.08E+15 | 5.38E+14 | 153.846 | <2.00E-16 |
| Treatment by Time | 12 | 1.44E+14 | 1.20E+13 | 3.431 | 0.000185 |
| Residuals | 147 | 5.14E+14 | 3.50E+12 | | |

| **Figure 3B** | **Df** | **Sum Sq** | **Mean Sq** | **F value** | **Pr(>F)** |
|---|---|---|---|---|---|
| Treatment | 5 | 1.15E+14 | 2.30E+13 | 13.919 | 1.63E-10 |
| Time | 2 | 7.34E+14 | 3.67E+14 | 222.459 | <2.00E-16 |
| Treatment by Time | 10 | 7.65E+13 | 7.65E+12 | 4.637 | 1.69E-05 |
| Residuals | 109 | 1.80E+14 | 1.65E+12 | | |

| **Figure 5B** | **Df** | **Sum Sq** | **Mean Sq** | **F value** | **Pr(>F)** |
|---|---|---|---|---|---|
| Treatment | 3 | 1.22E+17 | 4.06E+16 | 75.096 | <2.00E-16 |
| Time | 4 | 2.54E+16 | 6.36E+15 | 11.75 | 2.95E-08 |
| Treatment by Time | 12 | 5.57E+16 | 4.64E+15 | 8.578 | 4.56E-12 |
| Residuals | 140 | 7.57E+16 | 5.41E+14 | | |

| **Figure 5C** | **Df** | **Sum Sq** | **Mean Sq** | **F value** | **Pr(>F)** |
|---|---|---|---|---|---|
| Treatment | 3 | 1.06E+15 | 3.52E+14 | 54.89 | <2.00E-16 |
| Time | 2 | 5.33E+14 | 2.66E+14 | 41.51 | 1.02E-12 |
| Treatment by Time | 6 | 6.49E+14 | 1.08E+14 | 16.86 | 4.78E-12 |
| Residuals | 72 | 4.62E+14 | 6.42E+12 | | |

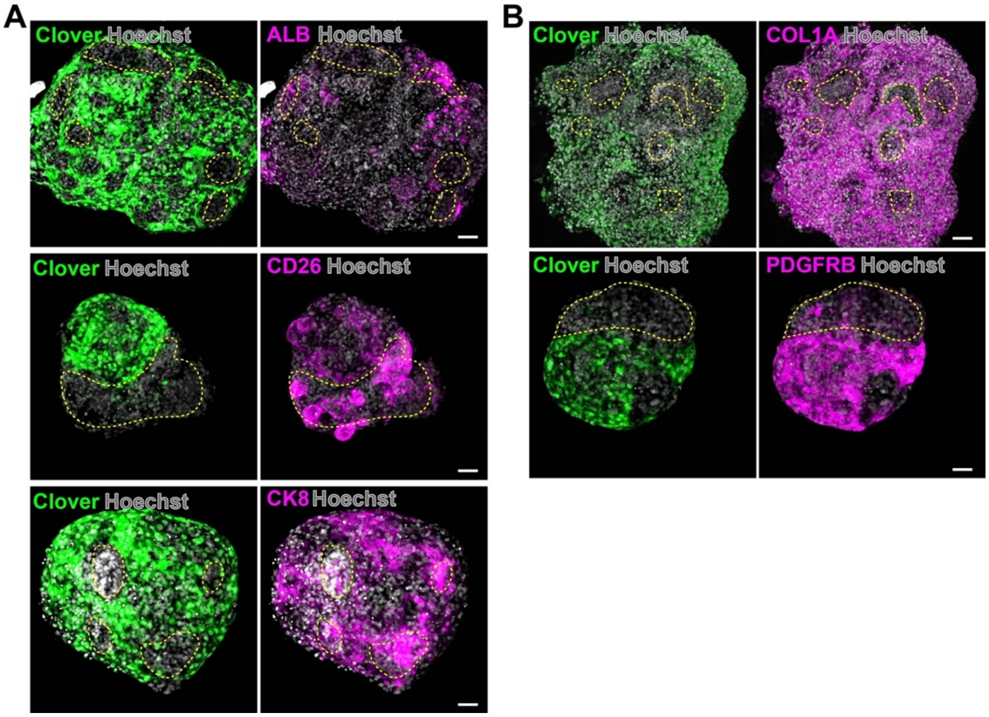

**Figure S1**. Immunofluorescent images of day 20 *COL1A1*-P2A Clover HOs stained with antibodies to Clover, Albumin (Alb), CD26 and CK8 (**A**); or antibodies to Clover, COL1A and PDGFRB (**B**). As seen in the merged images, the Clover$^+$ cells are distinct from the ALB$^+$ and CD26$^+$ and/or CK8$^+$ hepatocytes and cholangiocytes; while PDGFRB and COL1A are co-expressed in the Clover$^+$ cells. The dotted lines surround areas that are not stained with Clover and the Hoechst staining identifies nuclei. Scale bars, 50 $\mu$m.

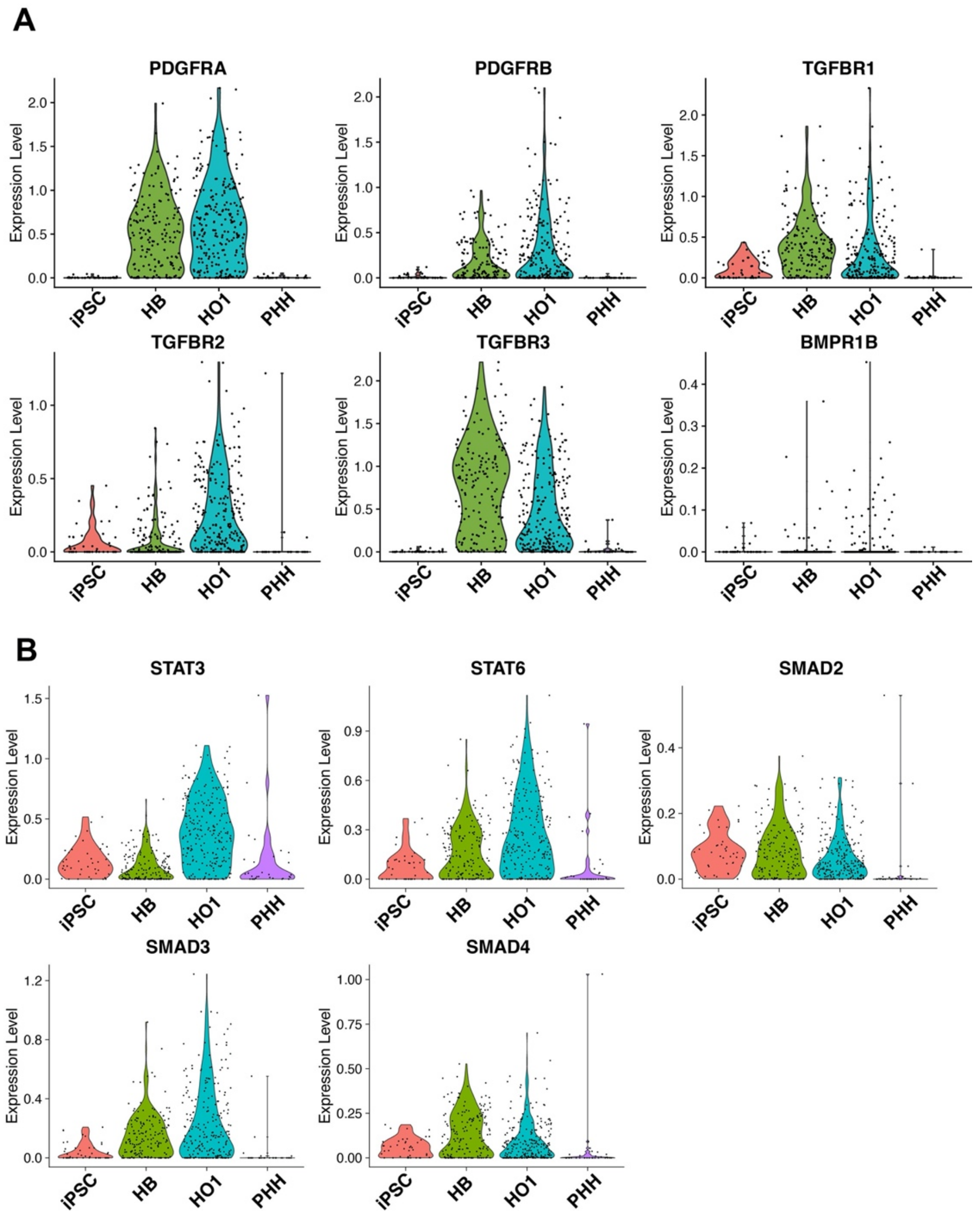

**Figure S2**. (**A, B**) Violin plots showing the level of expression of mRNAs encoding the receptors for PDGF (*PDGFRA, PDGFRB*) and TGF$\beta$1 (*TGFBR1, TGFBR2, TGFBR3, BMPR1B*); and of *STAT3*, *STAT6*, and *SMAD2-4* mRNAs in developing and mature HO cultures. Previously obtained scRNA-Seq data [30] was generated from iPSC (day 0), day 9 hepatoblast (HB) and day 21 mature organoid cultures. For comparison purposes, scRNA-Seq data obtained from primary human hepatocytes (PHH) is also shown.

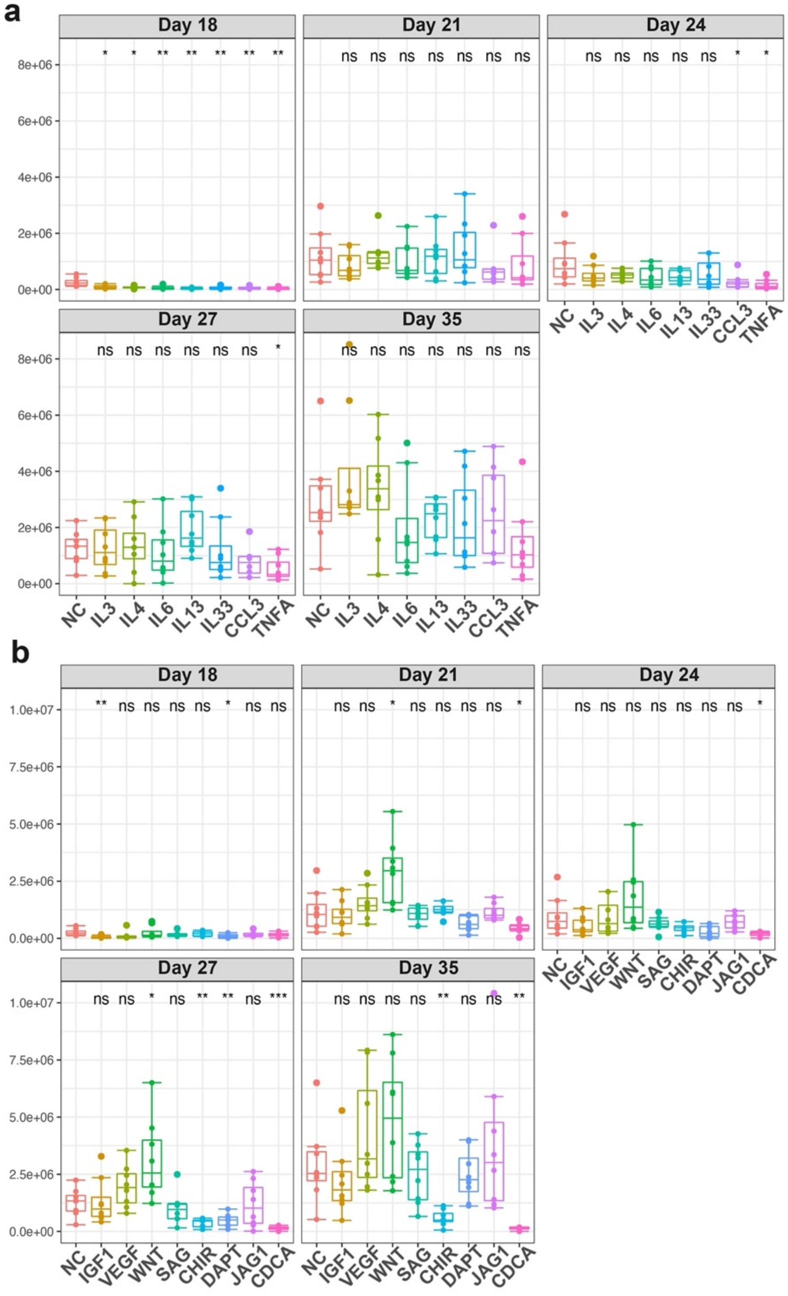

**Figure S3**. *COL1A1*-P2A Clover HOs were incubated with (**A**) 100 ng/ml IL-3, 100 ng/ml IL-4, 100 ng/ml IL-6, 100 ng/ml IL13, 100 ng/ml IL-33, 100 ng/ml CCL3, 50 ng/ml TNFa; or (**B**) 100 ng/ml IGF1, 50 ng/ml VEGF, 100 ng/ml Wnt3a, 10 μM SAG (Smoothened agonist), 3 μM CHIR99021, 10 μM DAPT (Gamma-Secretase Inhibitor), 100 ng/ml JAG1, 10 μM chenodeoxycholic acid (CDCA), or no addition (NC). Culture fluorescence, which indicates the amount of COL1A1$^{+}$ cells in the HOs, was serially measured on days 18 through 35. Each dot represents a measurement made on one HO, the thick line is the median of 8 organoids analyzed per condition, and the box plot shows the 25 to 75% range for all measurements per condition. With the possible exception of Wnt3a, none of these added agent caused a significant and or a sustained increase in COL1A1$^{+}$ cells in any of the organoid cultures. CDCA induced a decrease in COL1A1$^{+}$ cells.

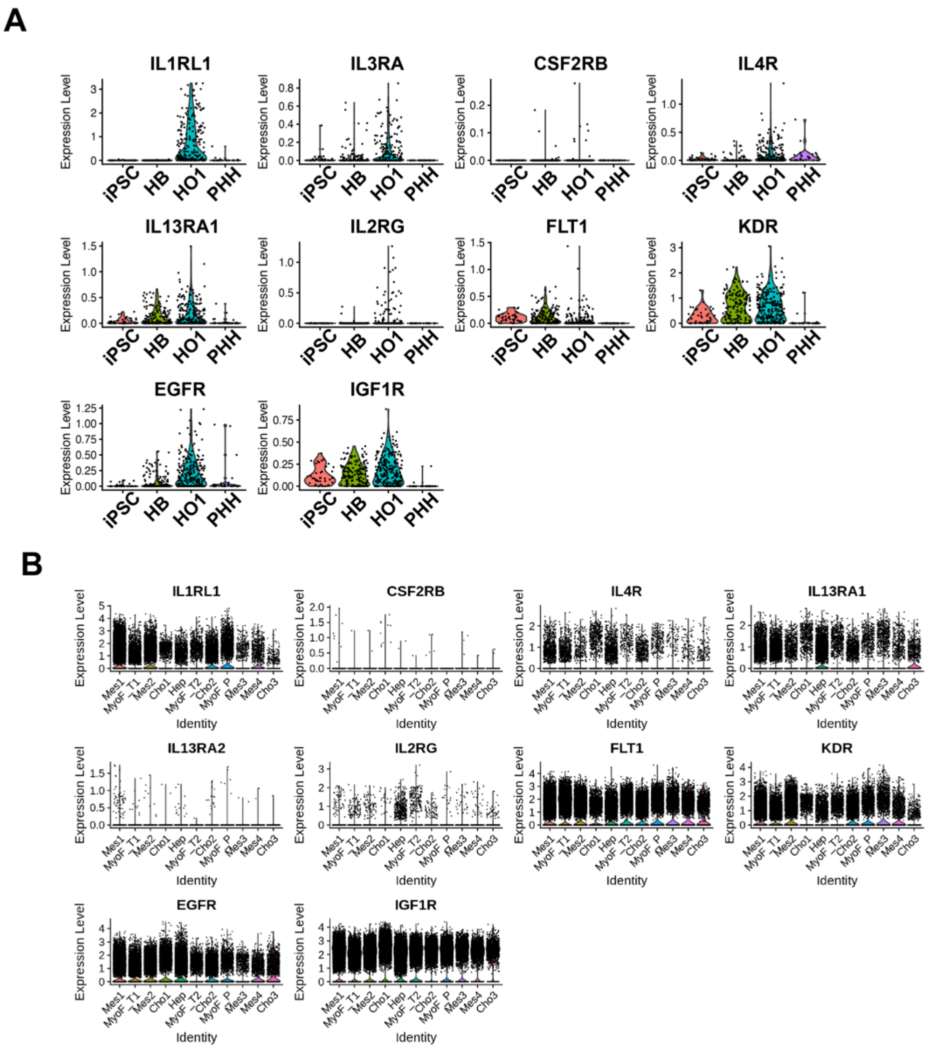

**Figure S4**. (**A**) Violin plots showing the level of expression of mRNAs for the following receptors during HO development: *IL1RL1, IL4R, IL13RA1, IL2RG, FLT1, KDR,* and *IGF1R*. scRNA-Seq data was generated from iPSC (day 0), day 9 hepatoblast (HB) and day 21 mature organoid (HO1) cultures. For comparison purposes, scRNA-Seq obtained from primary human hepatocytes (PHH) is also shown. This scRNA-Seq dataset was obtained from [30]. (**B**) Violin plots showing the level of mRNA expression for the receptors shown in (A) in the 11 cell clusters identified in day 21 control, PDGF- and TGFβ-treated microHOs using the scRNA-Seq data generated in this paper.

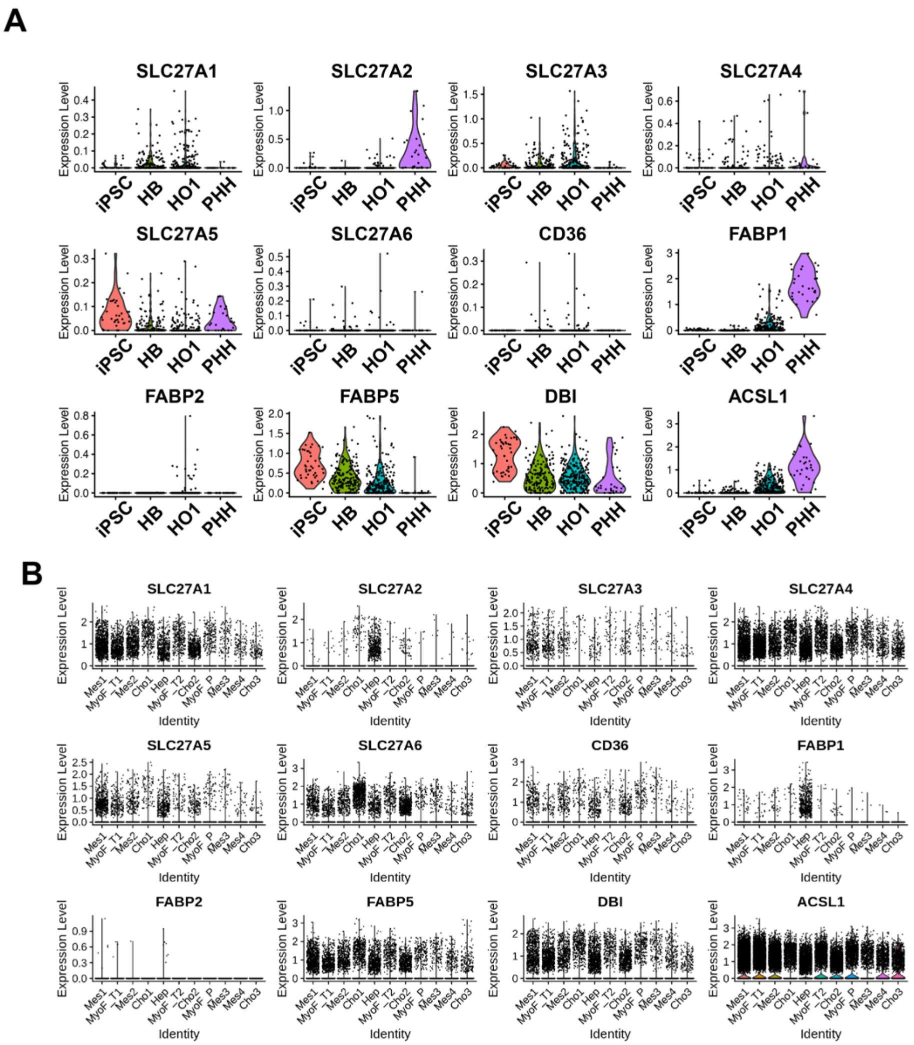

**Figure S5**. (**A**) Violin plots showing the level of expression of mRNAs for the following fatty acid transport proteins during HO development: *SLC27A1-6, CD36, FABP1-2, FABP5, DBI* and *ACSLI.* The previously obtained scRNA-Seq data [30] was generated from iPSC (day 0), day 9 hepatoblast (HB) and day 21 mature organoid cultures. For comparison purposes, scRNA-Seq obtained from primary human hepatocytes (PHH) is also shown. (**B**) Violin plots showing the level of mRNA expression for the for the fatty acid transport proteins shown in (A) in the eleven cell clusters identified in day 21 control, PDGF- and TGFβ-treated microHOs using the scRNA-Seq data generated in this paper.

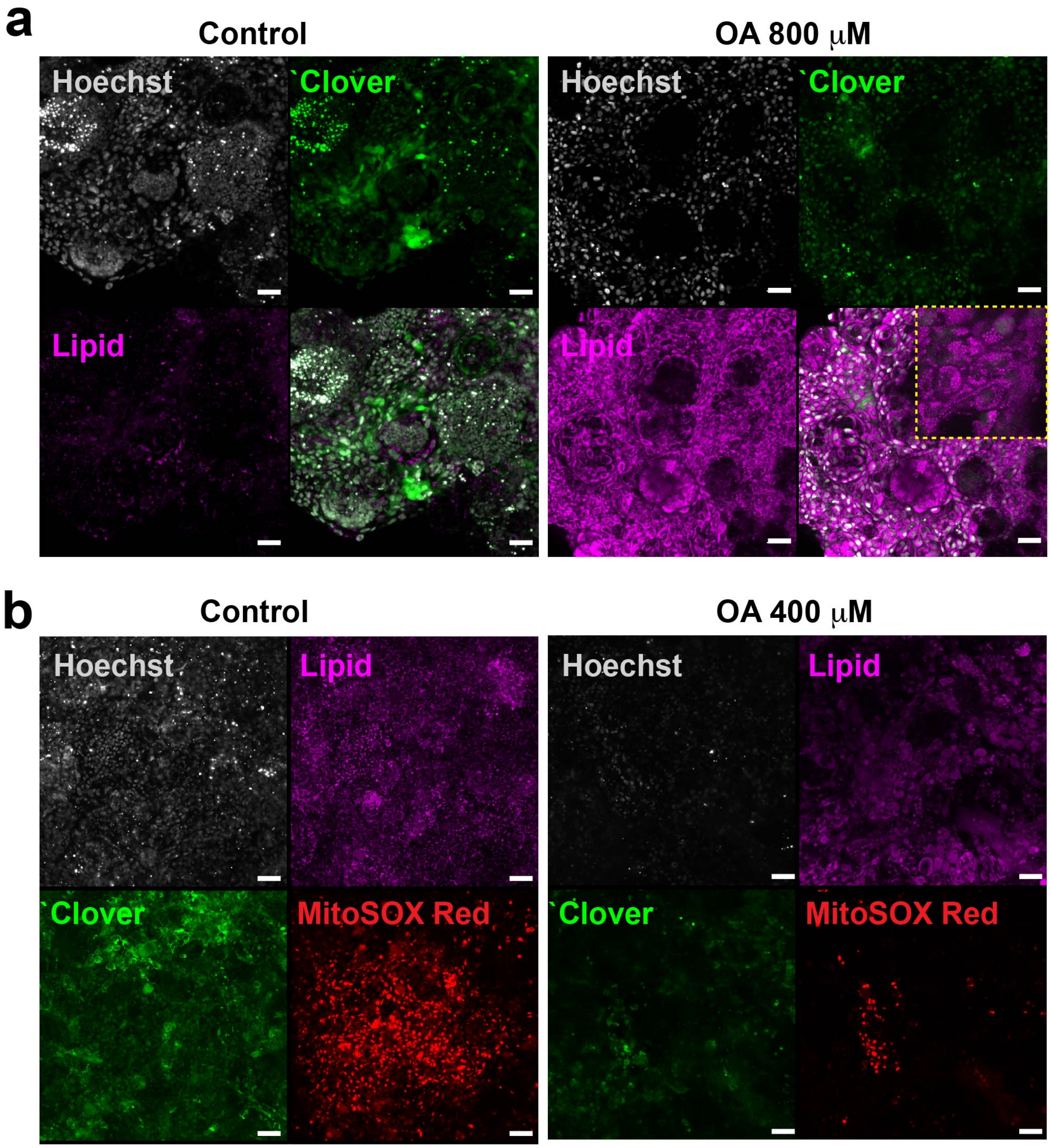

**Figure S6**. Oleic acid (OA) induces toxicity in HOs. Either 800 (**A**) or 400 (**B**) uM OA was added to HO cultures on day 13, and the HOs were analyzed by the HCS system on day 24. Although there was a marked increase in intracellular fatty acids within the HOs that were incubated with OA (LipidSpot), OA addition did not increase in COL1A1 expression (Clover). Moreover, there was a marked decrease in cell viability within the HOs that were incubated with OA, as shown by the MitoSox Red analysis.

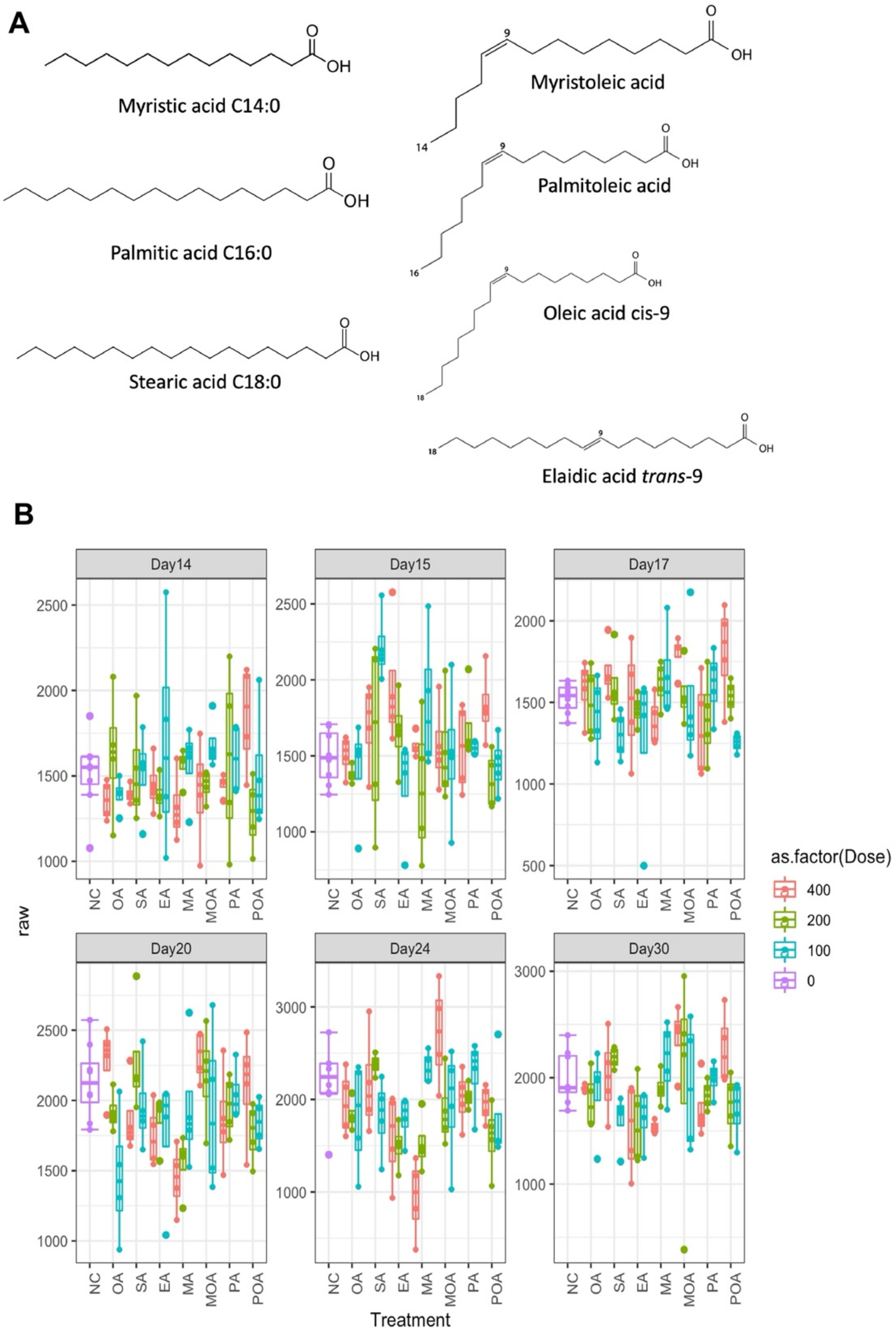

**Figure S7**. Free fatty acids do not increase collagen expression in HOs. (**A**) The structures of the free fatty acids whose effect on collagen expression in HOs was analyzed. Unsaturated FFAs (myristic (MA), palmitic (PA) and stearic (SA)) and their corresponding FFA with cis-9 (Myristoleic (MOA), palmitoleic (POA), and oleic (OA)) or trans-9 (elaidic (EA)) double bonds were analyzed. (**B**) *COL1A1*-P2A Clover HOs were incubated with 0, 100, 200 or 400 uM of the indicated free fatty acid on day 13. The fluorescence in these cultures, which indicates the amount of COL1A1$^{+}$ cells in the HOs, was serially measured on days 14 through 30. Each dot represents a measurement made on one HO, the thick line is the median of 8 organoids analyzed per condition, and the box plot shows the 25 to 75% range for all measurements per condition. The free fatty acids did not induce a significant increase in COL1A1$^{+}$ in any of the cultures.

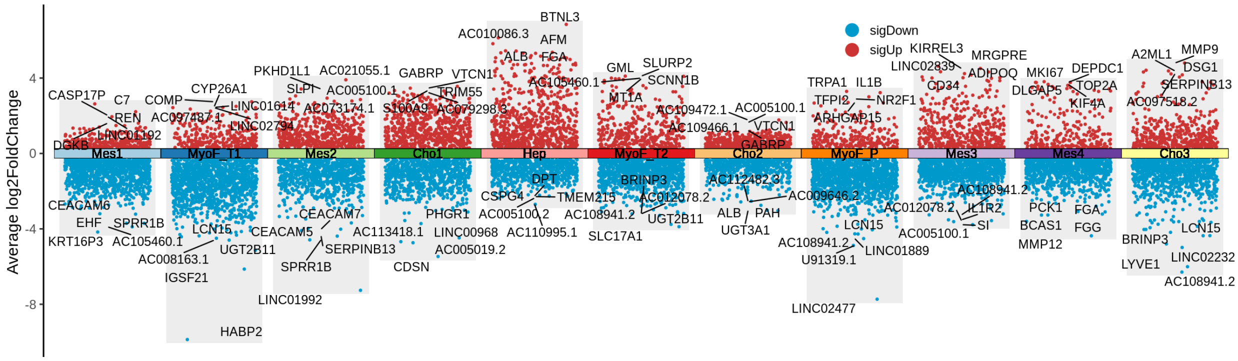

**Figure S8**. A combination volcano plot showing the top 5 DEGs for each of the 11 clusters. The red (or blue) dots indicate up-regulated (or down-regulated genes. As examples, Mes4 is the most proliferative cluster, and *MKI67* and *TOP2A* were among its DEGs; and the DEGs for the Hep cluster included hepatocyte specific markers (*ALB* and *FGA*).

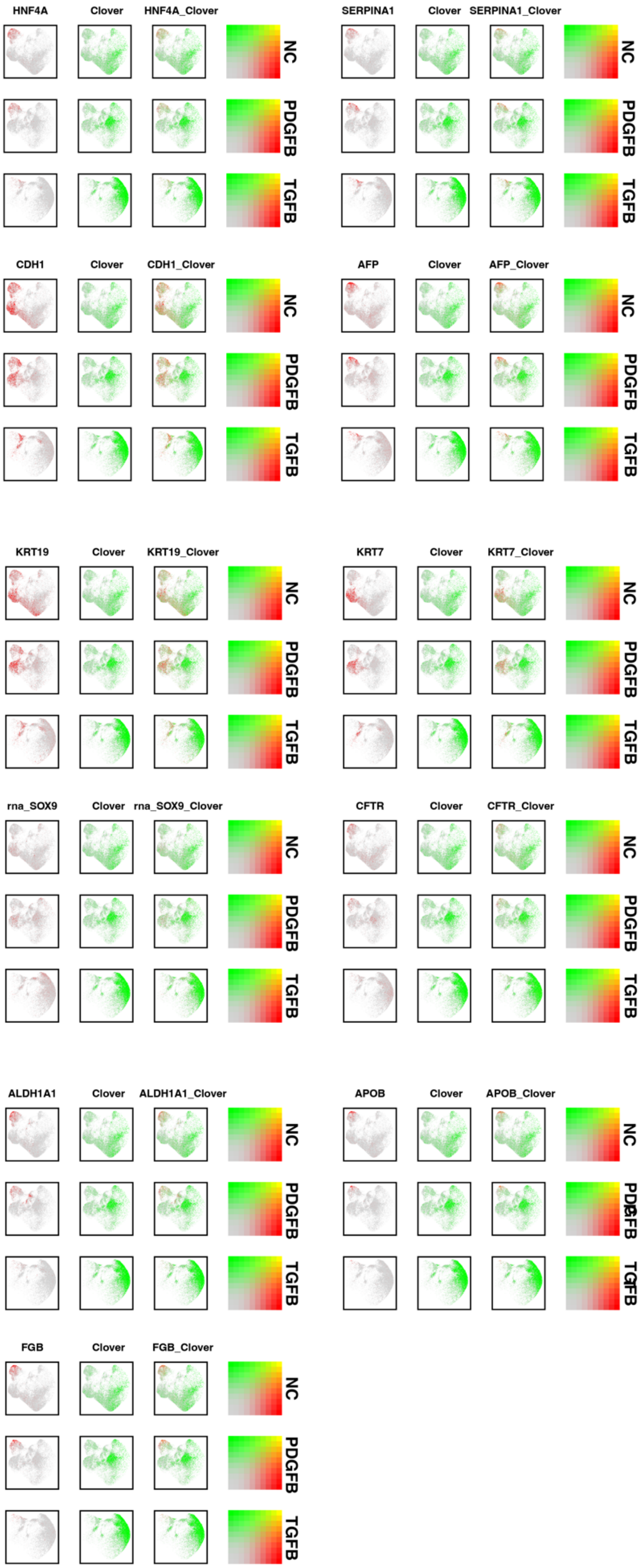
HNF4A
Clover
HNF4A_Clover
NC
PDGFB
TGFB
SERPINA1
Clover
SERPINA1_Clover
NC
PDGFB
TGFB
CDH1
Clover
CDH1_Clover
NC
PDGFB
TGFB
AFP
Clover
AFP_Clover
NC
PDGFB
TGFB
KRT19
Clover
KRT19_Clover
NC
PDGFB
TGFB
KRT7
Clover
KRT7_Clover
NC
PDGFB
TGFB
rna_SOX9
Clover
rna_SOX9_Clover
NC
PDGFB
TGFB
CFTR
Clover
CFTR_Clover
NC
PDGFB
TGFB
ALDH1A1
Clover
ALDH1A1_Clover
NC
PDGFB
TGFB
APOB
Clover
APOB_Clover
NC
PDGFB
TGFB
FGB
Clover
FGB_Clover
NC
PDGFB
TGFB

**Figure S9**. Feature plots show the level of expression of each indicated mRNA with *Clover* mRNA in the u-MAP plots shown in Fig 4. The uMAP plots for the normal control (NC), PDGF-, and TGFβ-treated microHOs are shown separately. The colors represent level of expression of the two mRNAs in each row as shown in the adjacent color threshold diagram. The mRNAs found in cholangiocytes (*KRT19*, *KRT7*, *CFTR),* hepatocytes *(SERPIN1A)* or hepatocytes and cholangiocytes (*HNF4A, AFP, CDH1, ALDHA1, APOB, SOX9, FGB*) were not expressed in the cells that expressed *Clover* mRNA in the microHOs.

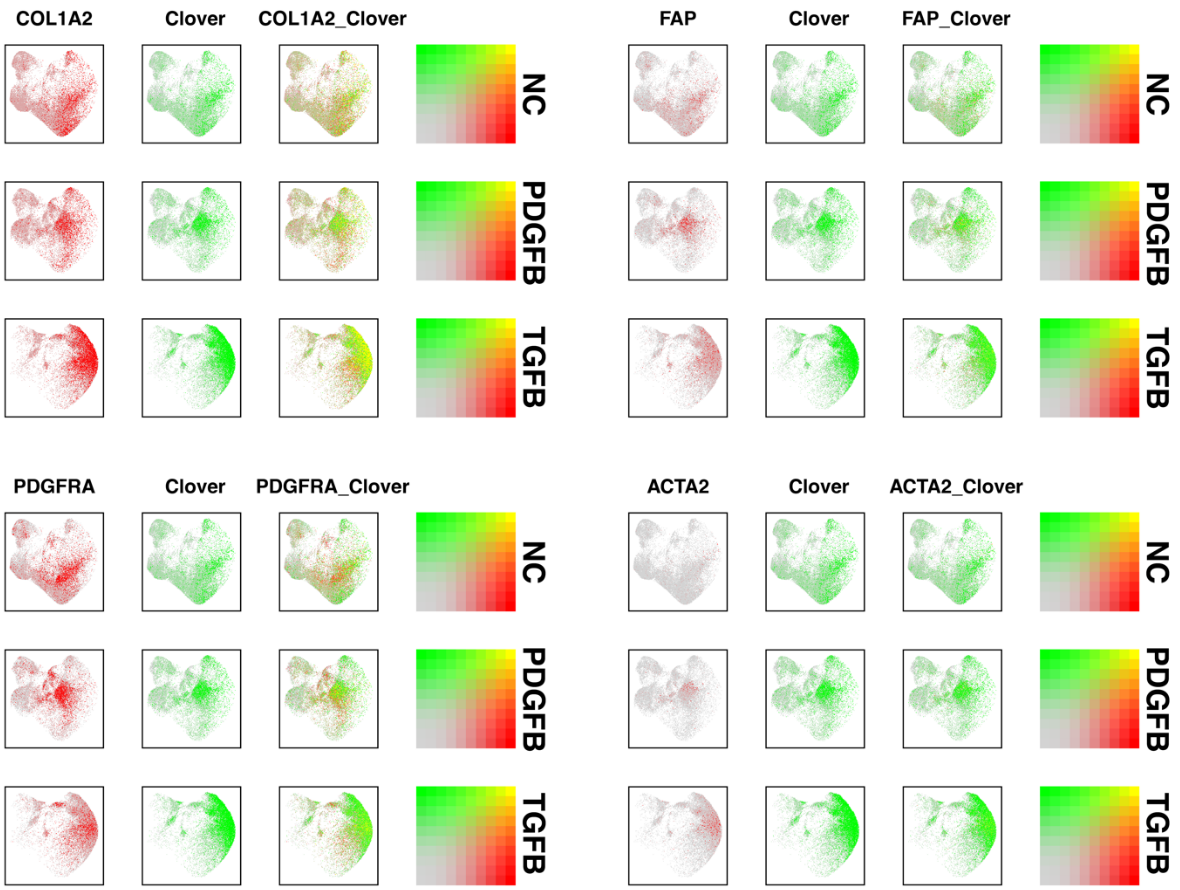

**Figure S10**. Feature plots show the level of expression of *COL1A2, PDGFRA, FAP or ACTA2* mRNAs with *Clover* mRNA in the u-MAP plots shown in Fig 8B. The UMAP plots for the normal control (NC), PDGF-, and TGFβ-treated microHOs are shown separately. The colors represent level of expression of the two mRNAs in each row as shown in the adjacent color threshold diagram. *COL1A2 and Clover* mRNAs have an overlapping pattern of expression; they are expressed in myofibroblasts and in mesenchymal cells. In contrast, *ACTA2, PDGFRA* and *FAP* mRNAs are predominantly expressed in the myofibroblasts (MyoF-T1/2, MyoF_P) in PDGF- or TGFβ-treated microHOs.

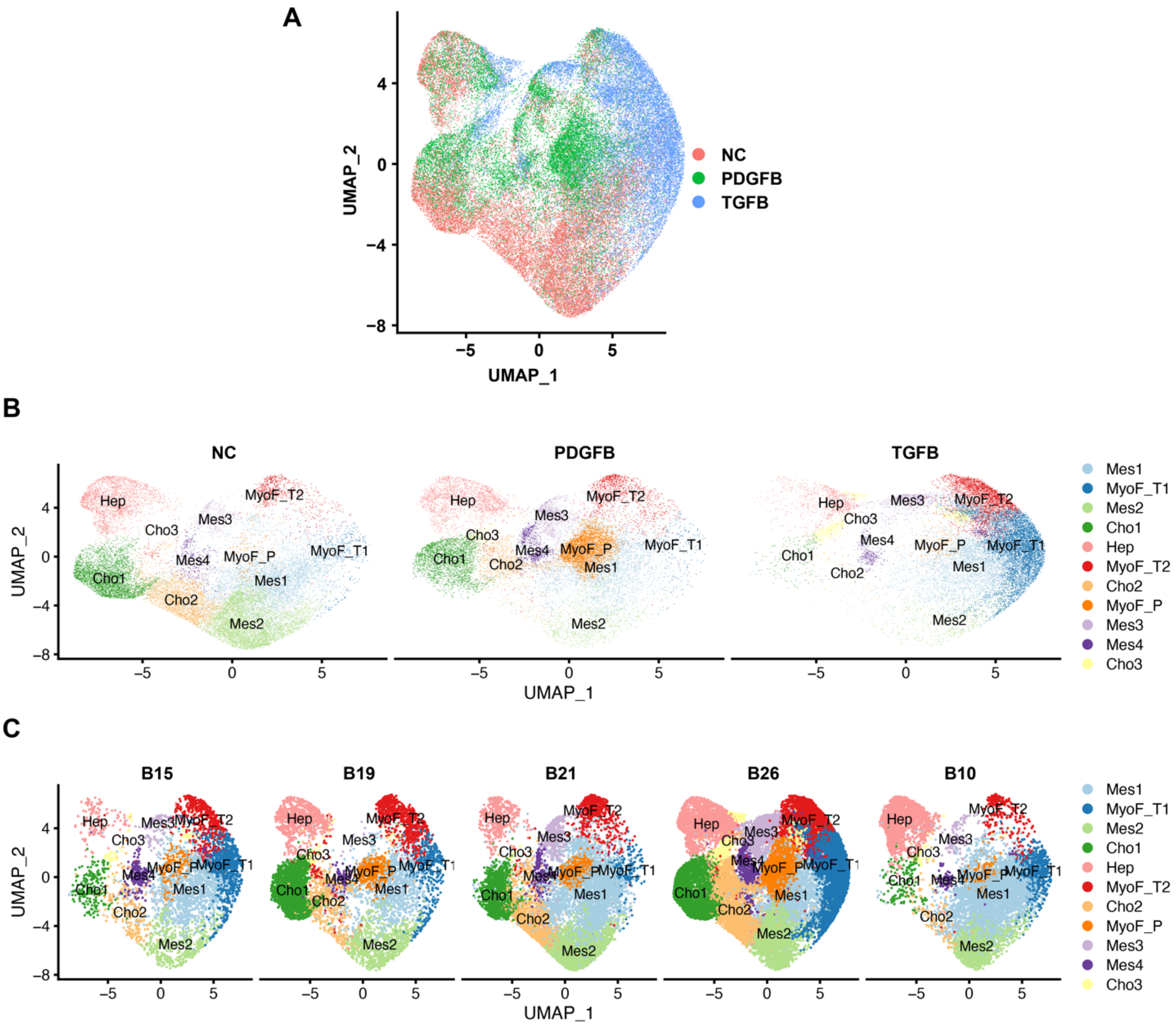

**Figure S11.** (**A**) A combined sample u-MAP plot shows the scRNA-seq data obtained from the cells in day 21 control (NC), PDGFβ-, or TGFβ1-treated microHO cultures. Each of the different type of microHO is indicated by the color shown on the right of the panel. (**B**) Separate u-MAP plots show the scRNA-seq data obtained from the cells in day 21 control (NC), PDGFβ-, or TGFβ1-treated microHO cultures. (**C**) The u-MAP plots for each of the five batches of microHOs that were separately analyzed by scRNA-Seq analysis. Each cluster is indicated by a different color as indicated on the right side of panels B and C.

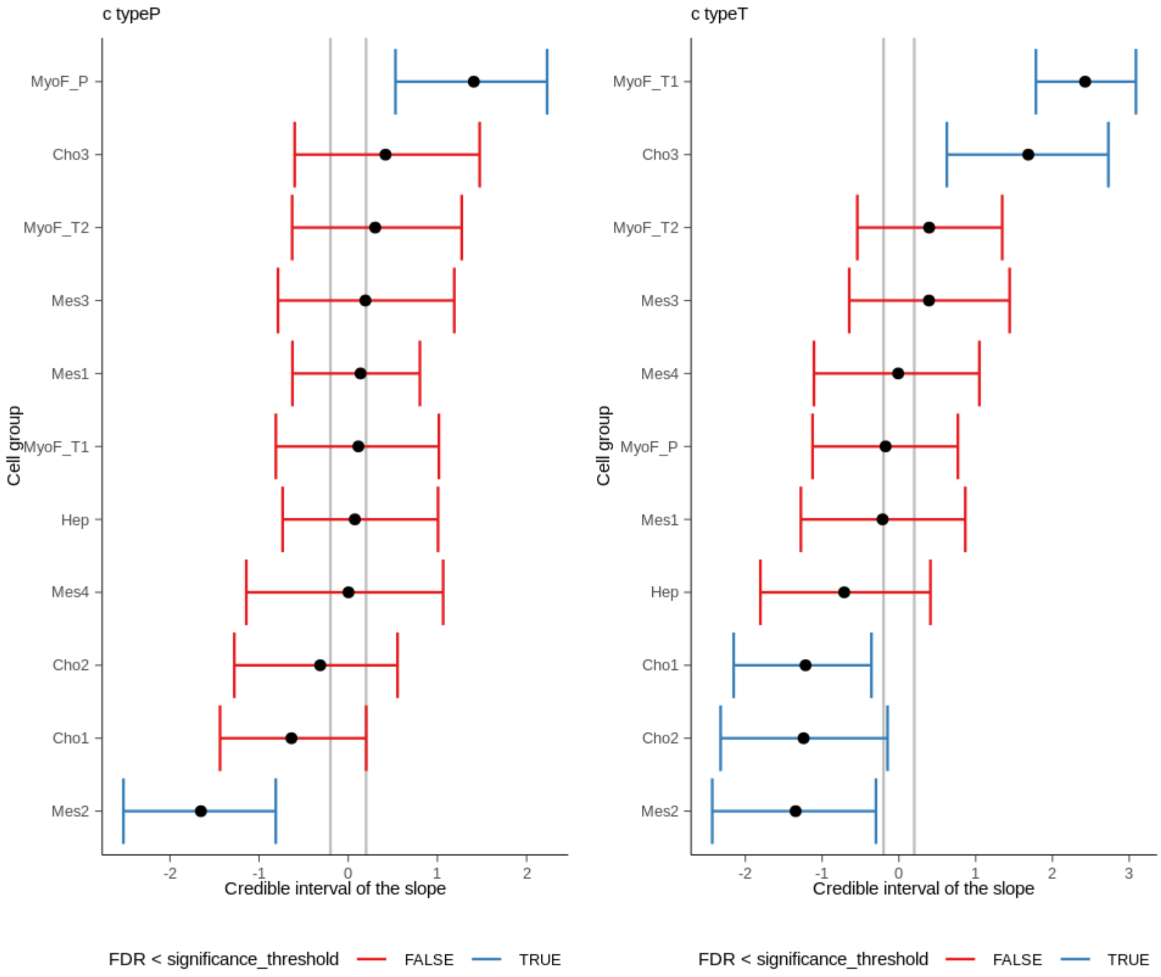

**Figure S12.** (**A**) Plots show the credible intervals determined by sccomp for each cell cluster in PDGFB-treated (Left) or TGFβ-treated (Right) microHOs vs NC microHOs. A blue color indicates that the treatment caused a significant change in the cell percentage.

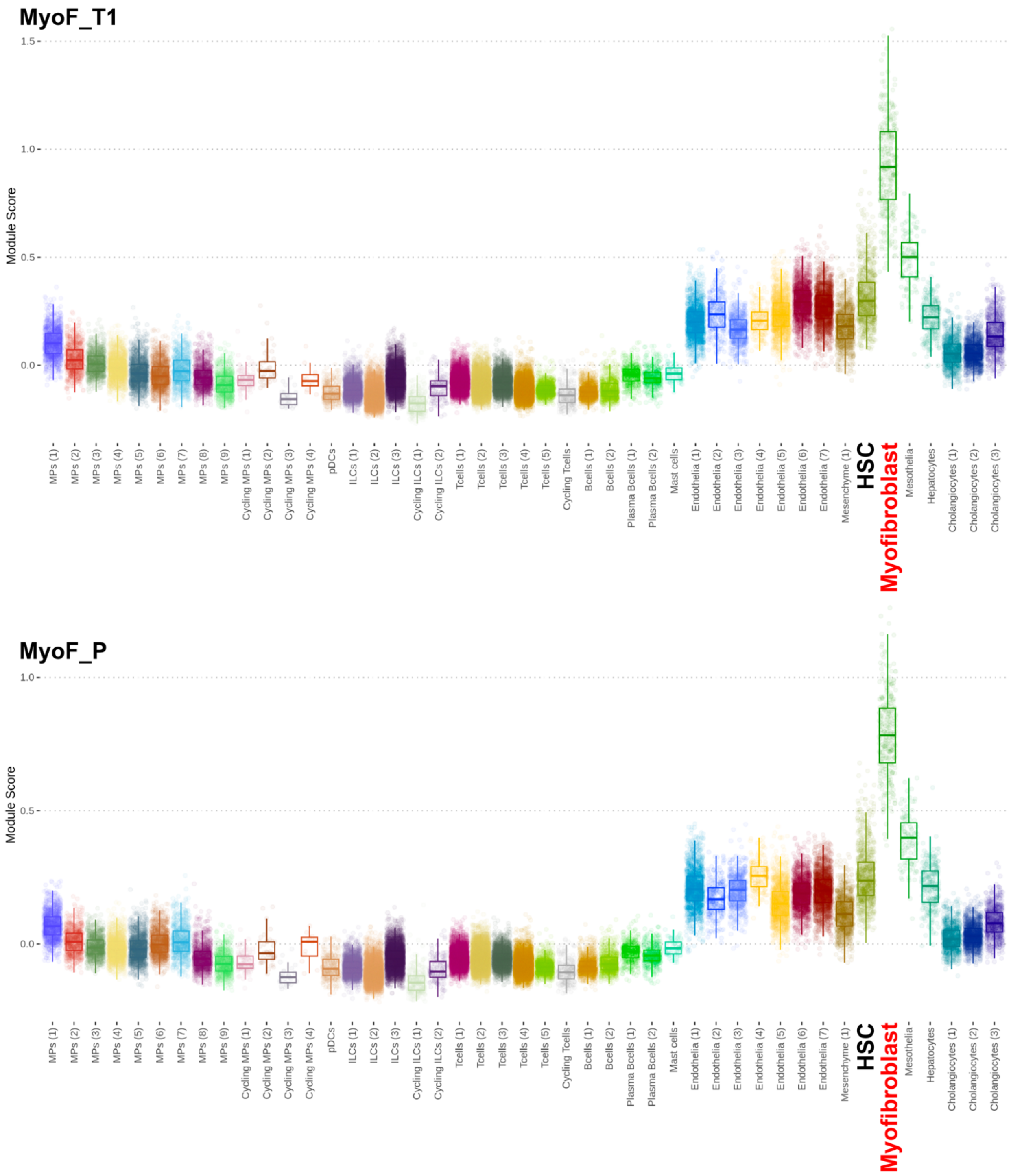

**Figure S13.** Among the multiple different types of cells present in normal and cirrhotic human liver, the MyoF_T1 and MyoF_P transcriptomes are most like that of myofibroblasts. The MyoF_T1 (n=53 genes) or MyoF_P (n=84 genes) signatures were compared with the transcriptomes of the various cell types in normal and cirrhotic human liver tissue (GSE136103 [22]), which include macrophages (MP) and innate lymphoid cells (ILC). ). Each dot shows the modulus score obtained when the gene signatures for the cell type indicated on the x-axis was compared with that of MyoF_T1 or MyoF_P. The thick line shows the mean, the boxplots show the 25 to 75% range, and the vertical line shows the Minimum (Q0 or 0th percentile) and

Maximum (Q4 or 100th percentile) of the scores. The MyoF_T1 and MyoF_P signatures are most enriched in the myofibroblasts in human liver tissue. The dotted lines represent actual reference module scores.

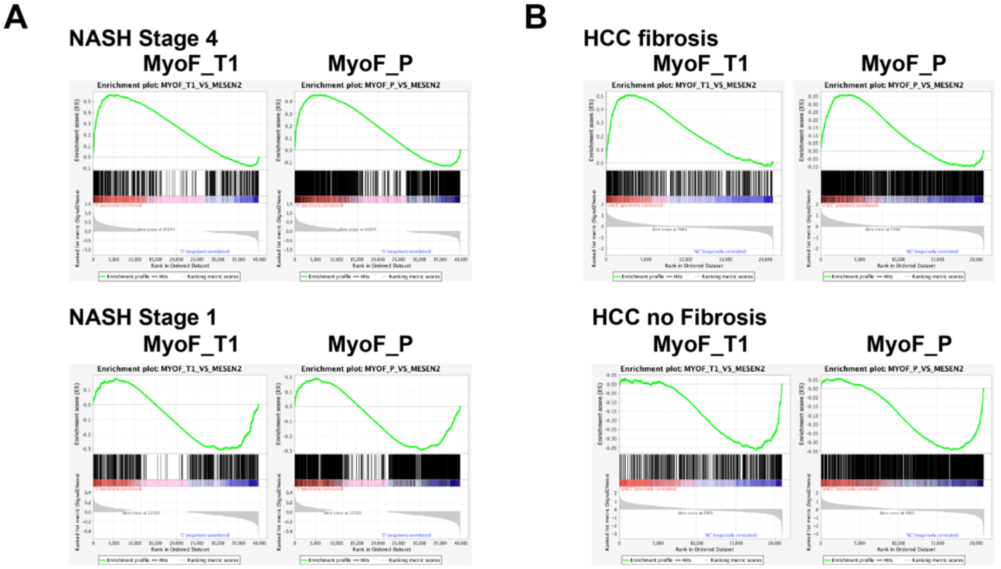

**Figure S14**. GSEA results assessing the correlation between the MyoF_T1 and MyoF_P gene signatures with that of non-fibrotic and fibrotic liver tissues caused by NASH **(A**) or hepatocellular carcinoma (HCC) **(B**). Genes whose expression levels were increased in MyoF_T1 (n=527) or MyoF_P (n=1716) relative to Mes2 (**Table S*)** were used to form myofibroblast-specific gene expression signatures. GSEA was performed using expression data obtained from (A) early (non-fibrotic) stage 1 and late (fibrotic) stage 4 NASH liver tissue (GSE13525 [2]) or (B) from resected HCC liver tissue, which was classified by histologic analysis as fibrotic or non-fibrotic (GSE6764 [8]). GSEA analyses revealed that the MyoF_T1 (NSE 1.7; false discovery rate (FDR) $3.3x10^{-4}$) and MyoF_P (1.4; FDR 0.013) signatures were strongly associated with stage 4 NASH liver tissue, but not with early (NES -0.97 and -0.95, FDR 1 and 0.99, respectively) stage 1 NASH liver tissue; and the MyoF_T1 (NES 2.77, FDR 0) and MyoF_P (NES 1.8, FDR $4.4x10^{-4}$) signatures were associated with liver fibrosis caused by HCC, whereas the MyoF_T1 (NES -1.98, FDR 0) and MyoF_P (NES -1.97, FDR 0) signatures were not associated with non-fibrotic HCC liver tissue.

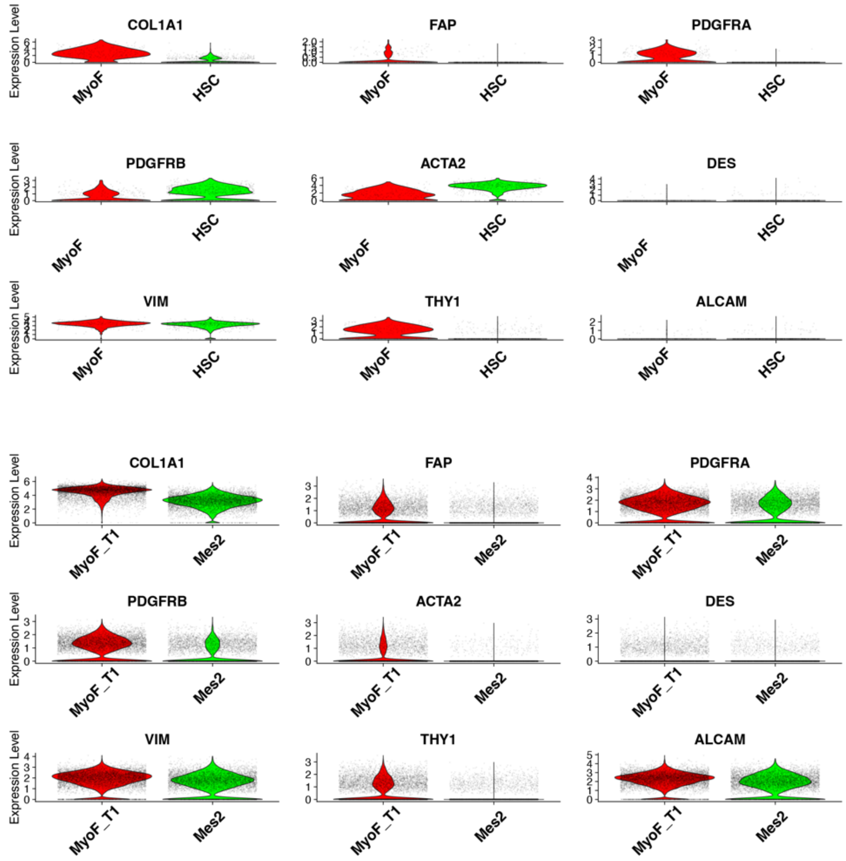

**Figure S15.** The pattern of expression of mesenchyme and MyoF-specific genes in hepatic stellate cells (HSC) and MyoF in human liver is mostly retained in Mes2 and MyoF_T1 cells in microHOs. scRNA-Seq data was analyzed to generate violin plots showing the level of expression of *COL1A1, FAP, PDGFRA, PDGFRB, ACTA2, DES, VIM, Thy1* and *ALCAM* mRNAs in HSC and MyoF in cirrhotic human liver (GSE136103; Top panel) and in Mes2 and MyoF_T1 cells in microHOs (Bottom panel). The cell cluster is indicated on the x-axis; and the y-axis shows the natural log transformed and normalized level of expression of each mRNA. Each dot shows the expression level in one cell.

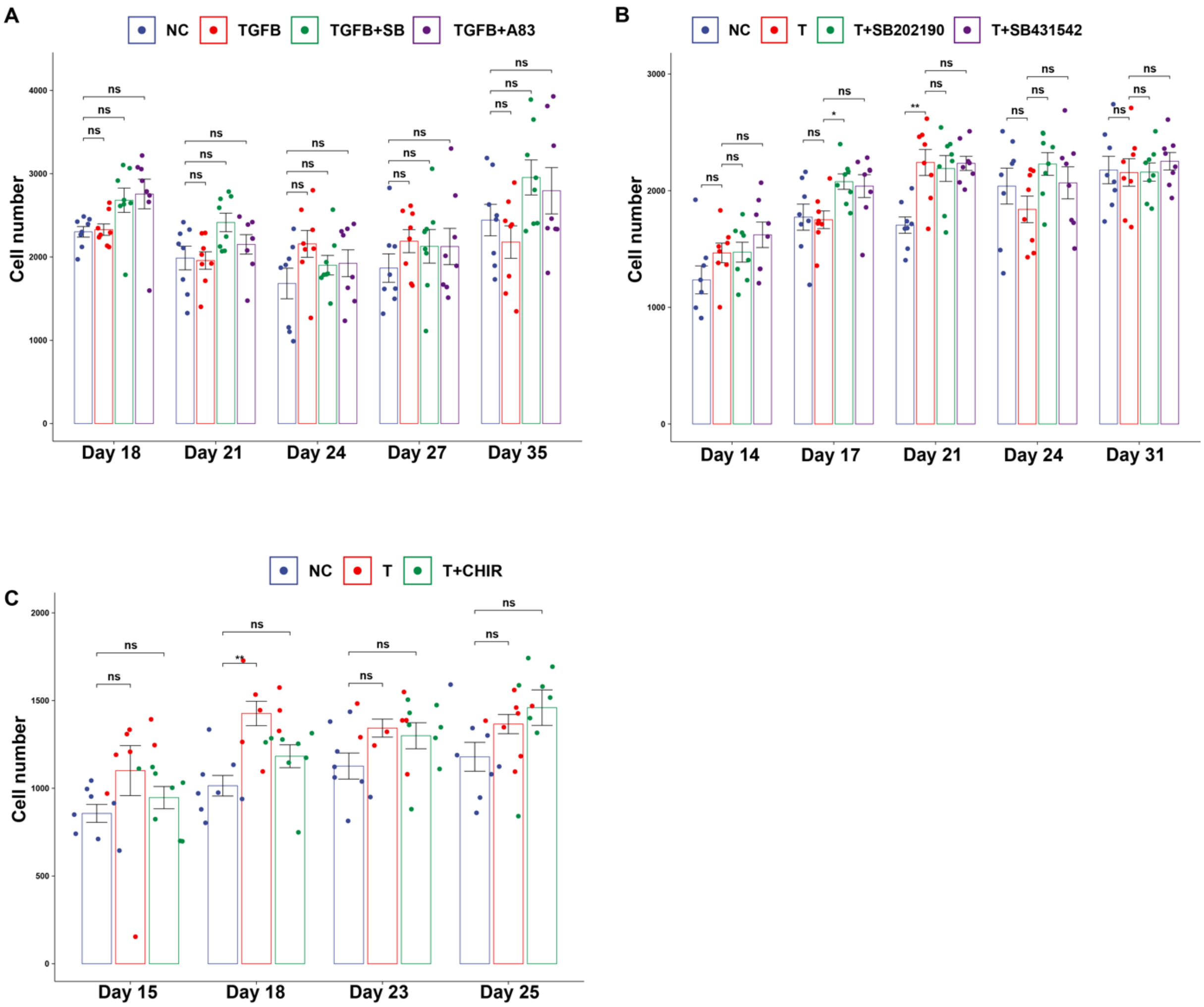

**Figure S16.** microHO viability was not affected by addition of TGFβ1, nor by any of the other tested drugs. microHOs were prepared and incubated with no addition (NC), 50 ng/ml TGFβ1 (T), or TGFβ1 ± the indicated inhibitor on day 13. The tested drugs were: (**A)** 10 uM TGFRβ1 tyrosine kinase inhibitors A83-01 or SB431542 (SB); (**B**) 10 uM TGFRβ1 inhibitor (SB431542) or 10 uM p38 α inhibitor (SB202190); and (**C**) 10 uM CHIR99021 (CHIR). The number of cells in each microHO was measured by counting the number of nuclei (stained with Hoechst 33342) on days 14 to 35. Each bar is the average of 8 individual measurements ± the SE, and individual datapoints are shown as dots. None of these additions caused a significant change (ns) in the number of cells in the treated microHOs relative to control microHOs.

**Table S4**. Twenty mRNAs that are differentially expressed in each of the 11 cell clusters identified in microHOs. The gene symbol, p-value, adjusted p-value, and fold differential expression ($log_2$) for each differentially expressed gene are shown. Pct.1 and Pct.2 indicates the contribution of that feature to defining the indicated cluster based upon principal component analysis.

| | cluster | gene | p_val | avg_log2FC | pct.1 | pct.2 | p_val_adj |
|---|---|---|---|---|---|---|---|
| **1** | 0 | *COL14A1* | 0.00 | 0.74 | 0.77 | 0.55 | 0 |
| **2** | 0 | *ADAMTS1* | 0.00 | 0.69 | 0.87 | 0.6 | 0 |
| **3** | 0 | *KCND2* | 0.00 | 0.65 | 0.75 | 0.53 | 1.41073166685289E-295 |
| **4** | 0 | *DLK1* | 0.00 | 0.64 | 0.97 | 0.88 | 0 |
| **5** | 0 | *IL1RL1* | 0.00 | 0.63 | 0.6 | 0.4 | 6.07272463504603E-113 |
| **6** | 0 | *COL6A3* | 0.00 | 0.58 | 0.99 | 0.93 | 0 |
| **7** | 0 | *IL6ST* | 0.00 | 0.57 | 0.99 | 0.95 | 0 |
| **8** | 0 | *ITGA1* | 0.00 | 0.56 | 0.93 | 0.74 | 0 |
| **9** | 0 | *ZNF804A* | 0.00 | 0.55 | 0.85 | 0.66 | 9.89305658045823E-202 |
| **10** | 0 | *CLMP* | 0.00 | 0.55 | 0.92 | 0.68 | 0 |
| **11** | 0 | *PDGFRA* | 0.00 | 0.54 | 0.91 | 0.76 | 0 |
| **12** | 0 | *SOX5* | 0.00 | 0.54 | 1 | 0.99 | 0 |
| **13** | 0 | *DCN* | 0.00 | 0.54 | 0.98 | 0.89 | 0 |
| **14** | 0 | *LAMA2* | 0.00 | 0.53 | 0.96 | 0.8 | 0 |
| **15** | 0 | *LUM* | 0.00 | 0.51 | 0.96 | 0.82 | 0 |
| **16** | 0 | *SERPINE2* | 0.00 | 0.49 | 0.99 | 0.93 | 0 |
| **17** | 0 | *DPP4* | 0.00 | 0.49 | 0.97 | 0.85 | 0 |
| **18** | 0 | *COL1A2* | 0.00 | 0.48 | 1 | 0.99 | 2.39332526066761E-224 |
| **19** | 0 | *RSPO2* | 0.00 | 0.47 | 0.9 | 0.69 | 0 |
| **20** | 0 | *TGFBR3* | 0.00 | 0.45 | 0.88 | 0.65 | 0 |
| **21** | 1 | *CILP* | 0.00 | 2.11 | 0.98 | 0.8 | 0 |
| **22** | 1 | *TIMP3* | 0.00 | 1.99 | 1 | 0.96 | 0 |
| **23** | 1 | *COMP* | 0.00 | 1.96 | 0.87 | 0.67 | 0 |
| **24** | 1 | *ITGA11* | 0.00 | 1.52 | 0.88 | 0.64 | 0 |
| **25** | 1 | *COL1A1* | 0.00 | 1.49 | 1 | 0.98 | 0 |

| | | | | | | | |
|---|---|---|---|---|---|---|---|
| **26** | 1 | *AC097487.1* | 0.00 | 1.48 | 0.65 | 0.59 | 0 |
| **27** | 1 | *TRABD2B* | 0.00 | 1.43 | 0.9 | 0.64 | 0 |
| **28** | 1 | *MXRA5* | 0.00 | 1.37 | 0.96 | 0.79 | 0 |
| **29** | 1 | *TIMP1* | 0.00 | 1.32 | 1 | 0.97 | 0 |
| **30** | 1 | *CCN2* | 0.00 | 1.32 | 0.86 | 0.67 | 0 |
| **31** | 1 | *ABTB2* | 0.00 | 1.32 | 0.98 | 0.76 | 0 |
| **32** | 1 | *LTBP2* | 0.00 | 1.30 | 0.91 | 0.67 | 0 |
| **33** | 1 | *NOX4* | 0.00 | 1.27 | 0.93 | 0.76 | 0 |
| **34** | 1 | *EGFP* | 0.00 | 1.27 | 0.96 | 0.82 | 0 |
| **35** | 1 | *SMYD3* | 0.00 | 1.27 | 0.99 | 0.93 | 0 |
| **36** | 1 | *AF165147.1* | 0.00 | 1.23 | 0.81 | 0.71 | 0 |
| **37** | 1 | *SPON1* | 0.00 | 1.22 | 0.75 | 0.55 | 0 |
| **38** | 1 | *SERPINE2* | 0.00 | 1.22 | 1 | 0.94 | 0 |
| **39** | 1 | *MICAL2* | 0.00 | 1.17 | 0.96 | 0.79 | 0 |
| **40** | 1 | *CLU* | 0.00 | 1.17 | 0.92 | 0.73 | 0 |
| **41** | 2 | *CNTN5* | 0.00 | 2.15 | 0.88 | 0.64 | 0 |
| **42** | 2 | *LINC01088* | 0.00 | 2.05 | 0.97 | 0.71 | 0 |
| **43** | 2 | *LINC02388* | 0.00 | 2.05 | 0.93 | 0.66 | 0 |
| **44** | 2 | *POSTN* | 0.00 | 1.70 | 0.95 | 0.74 | 0 |
| **45** | 2 | *NPNT* | 0.00 | 1.67 | 0.92 | 0.73 | 0 |
| **46** | 2 | *ARHGAP6* | 0.00 | 1.63 | 0.88 | 0.59 | 0 |
| **47** | 2 | *SLIT3* | 0.00 | 1.58 | 0.91 | 0.78 | 0 |
| **48** | 2 | *DIRC3* | 0.00 | 1.48 | 0.91 | 0.68 | 0 |
| **49** | 2 | *PKHD1L1* | 0.00 | 1.44 | 0.76 | 0.51 | 0 |
| **50** | 2 | *CLSTN2* | 0.00 | 1.35 | 0.82 | 0.62 | 0 |
| **51** | 2 | *BCO2* | 0.00 | 1.35 | 0.94 | 0.69 | 0 |
| **52** | 2 | *PDPN* | 0.00 | 1.29 | 0.98 | 0.87 | 0 |
| **53** | 2 | *CPS1* | 0.00 | 1.28 | 0.94 | 0.7 | 0 |
| **54** | 2 | *KCNMA1* | 0.00 | 1.26 | 0.94 | 0.8 | 0 |
| **55** | 2 | *LINC00894* | 0.00 | 1.26 | 0.92 | 0.68 | 0 |
| **56** | 2 | *PWWP3B* | 0.00 | 1.25 | 0.87 | 0.59 | 0 |

| | | | | | | | |
|---|---|---|---|---|---|---|---|
| **57** | 2 | *COL11A1* | 0.00 | 1.25 | 0.98 | 0.87 | 0 |
| **58** | 2 | *KDR* | 0.00 | 1.24 | 0.87 | 0.59 | 0 |
| **59** | 2 | *PRLR* | 0.00 | 1.24 | 0.96 | 0.77 | 0 |
| **60** | 2 | *GHR* | 0.00 | 1.23 | 0.88 | 0.68 | 0 |
| **61** | 3 | *UNC5C* | 0.00 | 2.45 | 0.96 | 0.62 | 0 |
| **62** | 3 | *GRHL2* | 0.00 | 2.36 | 0.91 | 0.64 | 0 |
| **63** | 3 | *ITGB6* | 0.00 | 2.32 | 0.96 | 0.7 | 0 |
| **64** | 3 | *GABRP* | 0.00 | 2.29 | 0.86 | 0.58 | 0 |
| **65** | 3 | *PDE4D* | 0.00 | 2.20 | 0.96 | 0.81 | 0 |
| **66** | 3 | *ARHGAP29* | 0.00 | 2.11 | 0.89 | 0.51 | 0 |
| **67** | 3 | *PLD5* | 0.00 | 2.08 | 0.86 | 0.67 | 0 |
| **68** | 3 | *ROR1* | 0.00 | 2.08 | 0.86 | 0.56 | 0 |
| **69** | 3 | *AC015522.1* | 0.00 | 2.00 | 0.98 | 0.77 | 0 |
| **70** | 3 | *PRTG* | 0.00 | 1.93 | 0.83 | 0.51 | 0 |
| **71** | 3 | *NRXN3* | 0.00 | 1.88 | 0.89 | 0.66 | 0 |
| **72** | 3 | *ST6GALNAC3* | 0.00 | 1.85 | 0.89 | 0.66 | 0 |
| **73** | 3 | *NEBL* | 0.00 | 1.85 | 0.91 | 0.75 | 0 |
| **74** | 3 | *ANKS1A* | 0.00 | 1.81 | 0.88 | 0.77 | 0 |
| **75** | 3 | *EYA1* | 0.00 | 1.76 | 0.67 | 0.53 | 0 |
| **76** | 3 | *NAALADL2* | 0.00 | 1.75 | 0.84 | 0.65 | 0 |
| **77** | 3 | *SEMA5A* | 0.00 | 1.74 | 0.95 | 0.73 | 0 |
| **78** | 3 | *PTN* | 0.00 | 1.67 | 0.8 | 0.55 | 0 |
| **79** | 3 | *THSD4* | 0.00 | 1.65 | 0.84 | 0.56 | 0 |
| **80** | 3 | *PATJ* | 0.00 | 1.64 | 0.82 | 0.63 | 0 |
| **81** | 4 | *AFP* | 0.00 | 3.58 | 0.72 | 0.53 | 0 |
| **82** | 4 | *CHST9* | 0.00 | 1.90 | 0.77 | 0.39 | 0 |
| **83** | 4 | *APOB* | 0.00 | 1.82 | 0.51 | 0.46 | 1.57013013564819E-97 |
| **84** | 4 | *CEACAM6* | 0.00 | 1.79 | 0.65 | 0.44 | 0 |
| **85** | 4 | *FGB* | 0.00 | 1.71 | 0.55 | 0.45 | 7.57761739439617E-259 |
| **86** | 4 | *LINC00511* | 0.00 | 1.68 | 0.87 | 0.54 | 0 |
| **87** | 4 | *NTN4* | 0.00 | 1.68 | 0.71 | 0.55 | 0 |

| | | | | | | | |
|---|---|---|---|---|---|---|---|
| **88** | 4 | *SERPINA1* | 0.00 | 1.58 | 0.53 | 0.43 | 3.26816794412982E-195 |
| **89** | 4 | *DNAJC15* | 0.00 | 1.47 | 0.91 | 0.73 | 0 |
| **90** | 4 | *CDH17* | 0.00 | 1.44 | 0.58 | 0.44 | 0 |
| **91** | 4 | *CDH2* | 0.00 | 1.38 | 0.95 | 0.79 | 0 |
| **92** | 4 | *FOXP2* | 0.00 | 1.37 | 0.73 | 0.55 | 0 |
| **93** | 4 | *EPCAM* | 0.00 | 1.26 | 0.75 | 0.52 | 0 |
| **94** | 4 | *APOA1* | 0.00 | 1.25 | 0.62 | 0.37 | 0 |
| **95** | 4 | *MARCHF1* | 0.00 | 1.23 | 0.64 | 0.44 | 0 |
| **96** | 4 | *CDH1* | 0.00 | 1.23 | 0.77 | 0.55 | 0 |
| **97** | 4 | *CCSER1* | 0.00 | 1.21 | 0.8 | 0.54 | 0 |
| **98** | 4 | *MAML3* | 0.00 | 1.20 | 0.9 | 0.75 | 0 |
| **99** | 4 | *LINC00278* | 0.00 | 1.20 | 0.78 | 0.59 | 0 |
| **100** | 4 | *LCP1* | 0.00 | 1.16 | 0.76 | 0.55 | 0 |
| **101** | 5 | *PI16* | 0.00 | 2.48 | 0.92 | 0.74 | 0 |
| **102** | 5 | *C3* | 0.00 | 2.00 | 0.74 | 0.59 | 0 |
| **103** | 5 | *IGFBP6* | 0.00 | 1.93 | 0.81 | 0.67 | 0 |
| **104** | 5 | *HTRA3* | 0.00 | 1.86 | 0.77 | 0.63 | 0 |
| **105** | 5 | *GABRB3* | 0.00 | 1.69 | 0.4 | 0.47 | 0.645287107 |
| **106** | 5 | *SLC24A2* | 0.00 | 1.49 | 0.75 | 0.65 | 0 |
| **107** | 5 | *KRT14* | 0.00 | 1.32 | 0.65 | 0.58 | 0 |
| **108** | 5 | *PRSS23* | 0.00 | 1.30 | 0.83 | 0.69 | 0 |
| **109** | 5 | *TIMP3* | 0.00 | 1.21 | 1 | 0.96 | 0 |
| **110** | 5 | *ATP10A* | 0.00 | 1.19 | 0.78 | 0.69 | 0 |
| **111** | 5 | *ANK2* | 0.00 | 1.19 | 0.85 | 0.73 | 0 |
| **112** | 5 | *TNS1* | 0.00 | 1.17 | 0.87 | 0.73 | 0 |
| **113** | 5 | *CSPG4* | 0.00 | 1.16 | 0.7 | 0.59 | 0 |
| **114** | 5 | *CD55* | 0.00 | 1.14 | 0.72 | 0.67 | 4.49192035504069E-284 |
| **115** | 5 | *KLF3* | 0.00 | 1.10 | 0.78 | 0.71 | 0 |
| **116** | 5 | *CPAMD8* | 0.00 | 1.07 | 0.58 | 0.42 | 0 |
| **117** | 5 | *AC099520.1* | 0.00 | 1.05 | 0.5 | 0.5 | 2.92243869181999E-49 |
| **118** | 5 | *ABI3BP* | 0.00 | 1.04 | 0.76 | 0.67 | 0 |

| | | | | | | | |
|---|---|---|---|---|---|---|---|
| **119** | 5 | *ZBTB7C* | 0.00 | 1.03 | 0.57 | 0.58 | 3.96295910582397E-99 |
| **120** | 5 | *FLNC* | 0.00 | 1.02 | 0.73 | 0.61 | 0 |
| **121** | 6 | *UNC5C* | 0.00 | 0.84 | 0.94 | 0.63 | 0 |
| **122** | 6 | *GABRP* | 0.00 | 0.78 | 0.79 | 0.6 | 0 |
| **123** | 6 | *ST6GALNAC3* | 0.00 | 0.78 | 0.9 | 0.67 | 0 |
| **124** | 6 | *GRHL2* | 0.00 | 0.77 | 0.89 | 0.65 | 0 |
| **125** | 6 | *PRTG* | 0.00 | 0.71 | 0.82 | 0.53 | 0 |
| **126** | 6 | *PDE4D* | 0.00 | 0.69 | 0.96 | 0.82 | 0 |
| **127** | 6 | *ROR1* | 0.00 | 0.69 | 0.85 | 0.57 | 0 |
| **128** | 6 | *ARHGAP29* | 0.00 | 0.68 | 0.83 | 0.54 | 0 |
| **129** | 6 | *ANKS1A* | 0.00 | 0.65 | 0.91 | 0.77 | 0 |
| **130** | 6 | *ITGB6* | 0.00 | 0.65 | 0.93 | 0.71 | 0 |
| **131** | 6 | *PLD5* | 0.00 | 0.62 | 0.84 | 0.68 | 0 |
| **132** | 6 | *LINC02388* | 0.00 | 0.61 | 0.74 | 0.69 | 3.14703394653217E-146 |
| **133** | 6 | *EYA1* | 0.00 | 0.60 | 0.67 | 0.54 | 1.6275168304562E-266 |
| **134** | 6 | *DIAPH3* | 0.00 | 0.60 | 0.68 | 0.55 | 1.51594497149573E-217 |
| **135** | 6 | *NRXN3* | 0.00 | 0.59 | 0.88 | 0.67 | 0 |
| **136** | 6 | *FREM2* | 0.00 | 0.59 | 0.66 | 0.52 | 1.11716953189223E-242 |
| **137** | 6 | *KCNQ5* | 0.00 | 0.58 | 0.73 | 0.57 | 5.17999574470969E-237 |
| **138** | 6 | *AC015522.1* | 0.00 | 0.58 | 0.97 | 0.78 | 0 |
| **139** | 6 | *PTN* | 0.00 | 0.57 | 0.71 | 0.57 | 6.4862311734237E-293 |
| **140** | 6 | *SEMA5A* | 0.00 | 0.57 | 0.94 | 0.74 | 0 |
| **141** | 7 | *TFPI2* | 0.00 | 2.00 | 0.62 | 0.46 | 0 |
| **142** | 7 | *PAPPA* | 0.00 | 1.65 | 0.98 | 0.89 | 0 |
| **143** | 7 | *PDE5A* | 0.00 | 1.55 | 0.73 | 0.57 | 0 |
| **144** | 7 | *ADAMTS1* | 0.00 | 1.53 | 0.8 | 0.65 | 0 |
| **145** | 7 | *AREG* | 0.00 | 1.35 | 0.69 | 0.64 | 5.25882782978605E-270 |
| **146** | 7 | *ITGA1* | 0.00 | 1.26 | 0.88 | 0.77 | 0 |
| **147** | 7 | *ABCC4* | 0.00 | 1.23 | 0.71 | 0.64 | 0 |
| **148** | 7 | *MSC-AS1* | 0.00 | 1.20 | 0.55 | 0.51 | 1.60206265130991E-185 |
| **149** | 7 | *PIEZO2* | 0.00 | 1.13 | 0.62 | 0.48 | 3.62254977687614E-291 |

| | | | | | | | |
|---|---|---|---|---|---|---|---|
| **150** | 7 | *SUCLG2-AS1* | 0.00 | 1.12 | 0.73 | 0.65 | 0 |
| **151** | 7 | *CLMP* | 0.00 | 1.11 | 0.79 | 0.72 | 0 |
| **152** | 7 | *KYNU* | 0.00 | 1.07 | 0.45 | 0.5 | 1.09972579275846E-41 |
| **153** | 7 | *IGFBP5* | 0.00 | 1.06 | 0.86 | 0.8 | 0 |
| **154** | 7 | *ROBO2* | 0.00 | 1.05 | 0.84 | 0.71 | 0 |
| **155** | 7 | *RORB* | 0.00 | 1.05 | 0.64 | 0.65 | 1.71419981781713E-181 |
| **156** | 7 | *MASP1* | 0.00 | 0.98 | 0.6 | 0.52 | 2.14815931343417E-221 |
| **157** | 7 | *AL691420.1* | 0.00 | 0.94 | 0.69 | 0.74 | 1.90329965031377E-129 |
| **158** | 7 | *KCND2* | 0.00 | 0.94 | 0.6 | 0.58 | 1.12904013826135E-87 |
| **159** | 7 | *PAMR1* | 0.00 | 0.93 | 0.66 | 0.64 | 1.05279467805712E-135 |
| **160** | 7 | *MMP16* | 0.00 | 0.91 | 0.79 | 0.73 | 6.45407093018707E-251 |
| **161** | 8 | *CXCL14* | 0.00 | 2.54 | 0.82 | 0.61 | 0 |
| **162** | 8 | *KIRREL3* | 0.00 | 2.47 | 0.64 | 0.58 | 1.78087718527724E-275 |
| **163** | 8 | *NRP2* | 0.00 | 2.07 | 0.81 | 0.59 | 0 |
| **164** | 8 | *GSN* | 0.00 | 2.01 | 0.93 | 0.84 | 0 |
| **165** | 8 | *AC002463.1* | 0.00 | 1.95 | 0.66 | 0.56 | 0 |
| **166** | 8 | *CRHBP* | 0.00 | 1.89 | 0.53 | 0.52 | 2.68844774508409E-87 |
| **167** | 8 | *ATRNL1* | 0.00 | 1.84 | 0.67 | 0.58 | 7.4046528162926E-205 |
| **168** | 8 | *NRG3* | 0.00 | 1.81 | 0.78 | 0.51 | 0 |
| **169** | 8 | *CDH13* | 0.00 | 1.79 | 0.63 | 0.48 | 1.90104856701329E-243 |
| **170** | 8 | *MIAT* | 0.00 | 1.79 | 0.7 | 0.56 | 0 |
| **171** | 8 | *NTM* | 0.00 | 1.77 | 0.59 | 0.55 | 1.11151682293596E-121 |
| **172** | 8 | *PLCB4* | 0.00 | 1.76 | 0.74 | 0.59 | 0 |
| **173** | 8 | *OXCT1* | 0.00 | 1.76 | 0.61 | 0.56 | 1.22313326537527E-195 |
| **174** | 8 | *NPAS2* | 0.00 | 1.59 | 0.67 | 0.53 | 1.64759745644261E-255 |
| **175** | 8 | *PI16* | 0.00 | 1.56 | 0.8 | 0.76 | 2.48410109644837E-303 |
| **176** | 8 | *CD109* | 0.00 | 1.54 | 0.75 | 0.7 | 3.32247596107369E-263 |
| **177** | 8 | *LINC01592* | 0.00 | 1.53 | 0.54 | 0.55 | 4.53240260414655E-120 |
| **178** | 8 | *TEK* | 0.00 | 1.48 | 0.73 | 0.63 | 7.56466873318405E-289 |
| **179** | 8 | *PARD3B* | 0.00 | 1.45 | 0.76 | 0.7 | 1.08312332593575E-226 |
| **180** | 8 | *PODXL* | 0.00 | 1.43 | 0.76 | 0.7 | 2.30063348380999E-244 |

| | | | | | | | |
|---|---|---|---|---|---|---|---|
| **181** | 9 | *MKI67* | 0.00 | 2.57 | 0.94 | 0.59 | 0 |
| **182** | 9 | *TOP2A* | 0.00 | 2.44 | 0.89 | 0.56 | 0 |
| **183** | 9 | *CENPF* | 0.00 | 2.23 | 0.89 | 0.58 | 0 |
| **184** | 9 | *ASPM* | 0.00 | 2.09 | 0.86 | 0.53 | 0 |
| **185** | 9 | *TPX2* | 0.00 | 2.02 | 0.88 | 0.57 | 0 |
| **186** | 9 | *CENPE* | 0.00 | 1.62 | 0.78 | 0.44 | 0 |
| **187** | 9 | *DIAPH3* | 0.00 | 1.56 | 0.88 | 0.55 | 0 |
| **188** | 9 | *ANLN* | 0.00 | 1.56 | 0.82 | 0.49 | 0 |
| **189** | 9 | *PRC1* | 0.00 | 1.39 | 0.73 | 0.44 | 0 |
| **190** | 9 | *KIF11* | 0.00 | 1.37 | 0.77 | 0.49 | 0 |
| **191** | 9 | *NUSAP1* | 0.00 | 1.36 | 0.71 | 0.41 | 0 |
| **192** | 9 | *CIT* | 0.00 | 1.34 | 0.75 | 0.5 | 0 |
| **193** | 9 | *KPNA2* | 0.00 | 1.31 | 0.73 | 0.46 | 3.90947789658089E-290 |
| **194** | 9 | *CDK1* | 0.00 | 1.27 | 0.68 | 0.51 | 1.05324536288692E-221 |
| **195** | 9 | *KIF4A* | 0.00 | 1.26 | 0.71 | 0.49 | 0 |
| **196** | 9 | *NCAPG* | 0.00 | 1.19 | 0.7 | 0.5 | 6.08843872357421E-298 |
| **197** | 9 | *CEP128* | 0.00 | 1.18 | 0.79 | 0.55 | 3.67991367428791E-298 |
| **198** | 9 | *KIF14* | 0.00 | 1.17 | 0.66 | 0.45 | 1.75116932862194E-259 |
| **199** | 9 | *CKAP2L* | 0.00 | 1.16 | 0.69 | 0.51 | 7.57255848194669E-251 |
| **200** | 9 | *MIR924HG* | 0.00 | 1.10 | 0.72 | 0.57 | 6.10640620425961E-203 |
| **201** | 10 | *MMP9* | 0.00 | 2.77 | 0.67 | 0.53 | 8.08397127732943E-193 |
| **202** | 10 | *LAMC2* | 0.00 | 2.72 | 0.97 | 0.74 | 0 |
| **203** | 10 | *KRT6A* | 0.00 | 2.63 | 0.89 | 0.67 | 0 |
| **204** | 10 | *LAMA3* | 0.00 | 2.51 | 0.8 | 0.65 | 2.88170788687901E-275 |
| **205** | 10 | *LAMB3* | 0.00 | 1.81 | 0.8 | 0.62 | 3.27852024209837E-276 |
| **206** | 10 | *KRT17* | 0.00 | 1.79 | 0.49 | 0.51 | 1.15843945877946E-40 |
| **207** | 10 | *COL17A1* | 0.00 | 1.77 | 0.68 | 0.59 | 5.38500051358062E-140 |
| **208** | 10 | *INPP4B* | 0.00 | 1.35 | 0.81 | 0.63 | 2.04825297095848E-212 |
| **209** | 10 | *KRT14* | 0.00 | 1.22 | 0.51 | 0.59 | 3.23324533571871E-21 |
| **210** | 10 | *TMEM132D* | 0.00 | 1.22 | 0.62 | 0.56 | 1.35736286299835E-92 |
| **211** | 10 | *SEMA3C* | 0.00 | 1.15 | 0.95 | 0.8 | 1.78100630981075E-230 |

| | | | | | | | |
|---|---|---|---|---|---|---|---|
| **212** | 10 | *THBS1* | 0.00 | 1.13 | 0.88 | 0.7 | 1.06459363129595E-135 |
| **213** | 10 | *DSC2* | 0.00 | 1.13 | 0.85 | 0.68 | 9.22506907450692E-173 |
| **214** | 10 | *ITGB4* | 0.00 | 1.12 | 0.49 | 0.45 | 3.83789725369348E-66 |
| **215** | 10 | *TNS4* | 0.00 | 1.09 | 0.42 | 0.52 | 1.58597324817715E-08 |
| **216** | 10 | *NTN4* | 0.00 | 1.09 | 0.69 | 0.56 | 8.74503754757718E-152 |
| **217** | 10 | *ITGA6* | 0.00 | 1.08 | 0.66 | 0.63 | 4.50495301171499E-66 |
| **218** | 10 | *LINC00511* | 0.00 | 1.08 | 0.84 | 0.57 | 1.39860819563529E-216 |
| **219** | 10 | *FRMD6* | 0.00 | 1.06 | 0.78 | 0.61 | 8.84377502264119E-144 |
| **220** | 10 | *CHST11* | 0.00 | 1.05 | 0.83 | 0.67 | 5.58462409190851E-165 |
